# Estimation of Non-null SNP Effect Size Distributions Enables the Detection of Enriched Genes Underlying Complex Traits

**DOI:** 10.1101/597484

**Authors:** Wei Cheng, Sohini Ramachandran, Lorin Crawford

**Affiliations:** Department of Ecology and Evolutionary Biology, Brown University, Providence, RI, USA; Center for Computational Molecular Biology, Brown University, Providence, RI, USA; Department of Biostatistics, Brown University, Providence, RI, USA; Center for Statistical Sciences, Brown University, Providence, RI, USA

## Abstract

Traditional univariate genome-wide association studies generate false positives and negatives due to difficulties distinguishing associated variants from variants with spurious nonzero effects that do not directly influence the trait. Recent efforts have been directed at identifying genes or signaling pathways enriched for mutations in quantitative traits or case-control studies, but these can be computationally costly and hampered by strict model assumptions. Here, we present gene-*ε*, a new approach for identifying statistical associations between sets of variants and quantitative traits. Our key insight is that enrichment studies on the gene-level are improved when we reformulate the genome-wide SNP-level null hypothesis to identify spurious small-to-intermediate SNP effects and classify them as non-causal. gene-*ε* efficiently identifies enriched genes under a variety of simulated genetic architectures, achieving greater than a 90% true positive rate at 1% false positive rate for polygenic traits. Lastly, we apply gene-*ε* to summary statistics derived from six quantitative traits using European-ancestry individuals in the UK Biobank, and identify enriched genes that are in biologically relevant pathways.

**Author Summary:** Enrichment tests augment the standard univariate genome-wide association (GWA) framework by identifying groups of biologically interacting mutations that are enriched for associations with a trait of interest, beyond what is expected by chance. These analyses model local linkage disequilibrium (LD), allow many different mutations to be disease-causing across patients, and generate biologically interpretable hypotheses for disease mechanisms. However, existing enrichment analyses are hampered by high computational costs, and rely on GWA summary statistics despite the high false positive rate of the standard univariate GWA framework. Here, we present the gene-level association framework gene-*ε* (pronounced “genie”), an empirical Bayesian approach for identifying statistical associations between sets of mutations and quantitative traits. The central innovation of gene-*ε* is reformulating the GWA null model to distinguish between *(i)* mutations that are statistically associated with the disease but are unlikely to directly influence it, and *(ii)* mutations that are most strongly associated with a disease of interest. We find that, with our reformulated SNP-level null hypothesis, our gene-level enrichment model outperforms existing enrichment methods in simulation studies and scales well for application to emerging biobank datasets. We apply gene-*ε* to six quantitative traits in the UK Biobank and recover novel and functionally validated gene-level associations.

## Introduction

Over the last decade, there has been an evolving debate about the types of insight genome-wide single-nucleotide polymorphism (SNP) genotype data offer into the genetic architecture of complex traits [1–5]. In the traditional genome-wide association (GWA) framework, individual SNPs are tested independently for association with a trait of interest. While this approach can have drawbacks [2, 3, 6], more recent approaches that combine SNPs within a region have gained power to detect biologically relevant genes and pathways enriched for correlations with complex traits [7–14]. Reconciling these two observations is crucial for biomedical genomics.

In the traditional GWA model, each SNP is assumed to either (*i*) directly influence (or perfectly tag a variant that directly influences) the trait of interest; or (*ii*) have no affect on the trait at all (see Fig. 1A). Throughout this manuscript, for simplicity, we refer to SNPs under the former as “associated” and those under latter as “non-associated”. These classifications are based on ordinary least squares (OLS) effect size estimates for each SNP in a regression framework, where the null hypothesis assumes that the true effects of non-associated SNPs are zero (*H*_0_ : *β*_*j*_ = 0). The traditional GWA model is agnostic to trait architecture, and is underpowered with a high false-positive rate for “polygenic” traits or traits which are generated by many mutations of small effect [5, 15–17].

**Figure 1.**
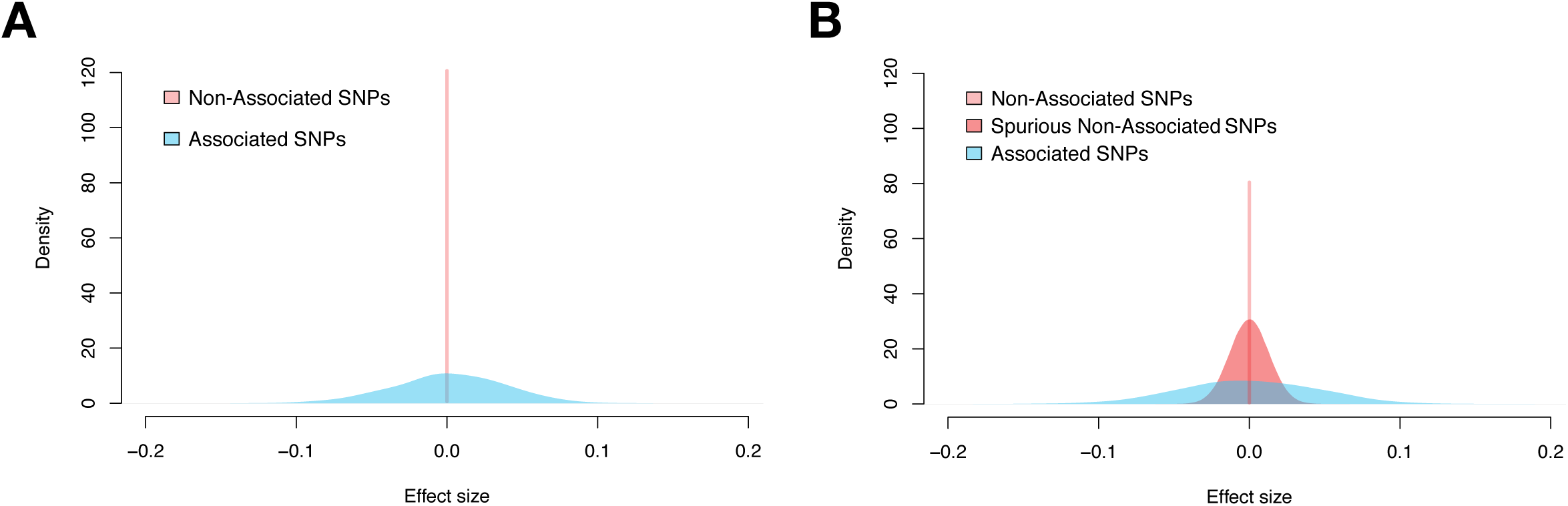
Illustration of null hypothesis assumptions for the distribution of GWA SNP-level effect sizes according to different views on underlying genetic architectures. The effect sizes of “non-associated” (pink), “spurious non-associated” (red), and “associated” (blue) SNPs were drawn from normal distributions with successively larger variances. **(A)** The traditional GWA model of complex traits simply assumes SNPs are associated or non-associated. Under the corresponding null hypothesis, associated SNPs are likely to emit nonzero effect sizes while non-associated SNPs will have effect sizes of zero. When there are many causal variants, we refer to the traits as polygenic. **(B)** Under our reformulated GWA model, there are three categories: associated SNPs, non-associated SNPs that emit spurious nonzero effect sizes, and non-associated SNPs with effect sizes of zero. We propose a multi-component framework (see also [18]), in which null SNPs can emit different levels of statistical signals based on (*i*) different degrees of connectedness (e.g., through linkage disequilibrium), or (*ii*) its regulated gene interacts with an enriched gene. While truly associated SNPs are still more likely to emit large effect sizes than SNPs in the other categories, null SNPs can have intermediate effect sizes. Here, our goal is to treat spurious SNPs with small-to-intermediate nonzero effects as being non-associated with the trait of interest.

Suppose that in truth each SNP in a GWA dataset instead belongs to one of *three* categories depending on the underlying distribution of their effects on the trait of interest: (*i*) associated SNPs; (*ii*) non-associated SNPs that emit spurious nonzero statistical signals; and (*iii*) non-associated SNPs with zero-effects (Fig. 1B) [18]. Associated SNPs may lie in enriched genes that directly influence the trait of interest. The phenomenon of a non-associated SNP emitting nonzero statistical signal can occur due to multiple reasons. For example, spurious nonzero SNP effects can be due to some varying degree of linkage disequilibrium (LD) with associated SNPs [19]; or alternatively, non-associated SNPs can have a trans-interaction effect with SNPs located within an enriched gene. In either setting, spurious SNPs can emit small-to-intermediate statistical noise (in some cases, even appearing indistinguishable from truly associated SNPs), thereby confounding traditional GWA tests (Fig. 1B). Hereafter, we refer to this noise as “epsilon-genic effects” (denoted in shorthand as “*ε*-genic effects”). There is a need for a computational framework that has the ability to identify mutations associated with a wide range of traits, regardless of whether narrow-sense heritability is sparsely or uniformly distributed across the genome.

Here, we develop a new and scalable quantitative approach for testing aggregated sets of SNP-level GWA summary statistics for enrichment of associated mutations in a given quantitative trait. In practice, our approach can be applied to any user-specified set of genomic regions, such as regulatory elements, intergenic regions, or gene sets. In this study, for simplicity, we refer to our method as a gene-level test (i.e., an annotated collection of SNPs within the boundary of a gene). The key contribution of our approach is that gene-level association tests should treat spurious SNPs with *ε*-genic effects as non-associated variants. Conceptually, this requires assessing whether SNPs explain more than some “epsilon” proportion of the phenotypic variance. In this generalized model, we reformulate the GWA null hypothesis to assume *approximately* no association for spurious non-associated SNPs where

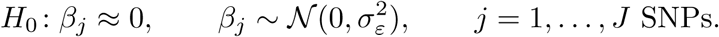

Here, 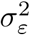 denotes a “SNP-level null threshold” and represents the maximum proportion of phenotypic variance explained (PVE) that is contributed by spurious non-associated SNPs. This null hypothesis can be equivalently restated as 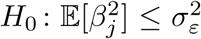 (Fig. 1B). Non-enriched genes are then defined as genes that only contain SNPs with *ε*-genic effects (i.e., 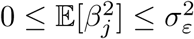 for every *j*-th SNP within that region). Enriched genes, on the other hand, are genes that contain at least one associated SNP (i.e., 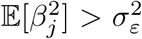 for at least one SNP *j* within that region). By accounting for the presence of spurious *ε*-genic effects (i.e., through different values of 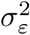 which the user can subjectively control), our approach flexibly constructs an appropriate GWA SNP-level null hypothesis for a wide range of traits with genetic architectures that land anywhere on the polygenic spectrum (see Materials and Methods).

We refer to our gene-level association framework as “gene-*ε*” (pronounced “genie”). gene-*ε* leverages our modified SNP-level null hypothesis to lower false positive rates and increases power for identifying gene-level enrichment within GWA studies. This happens via two key conceptual insights. First, gene-*ε* regularizes observed (and inflated) GWA summary statistics so that SNP-level effect size estimates are positively correlated with the assumed generative model of complex traits. Second, it examines the distribution of regularized effect sizes to offer the user choices for an appropriate SNP-level null threshold 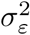 to distinguish associated SNPs from spurious non-associated SNPs. This makes for an improved and refined hypothesis testing strategy for identifying enriched genes underlying complex traits. With detailed simulations, we assess the power of gene-*ε* to identify significant genes under a variety of genetic architectures, and compare its performance against multiple competing approaches [7, 10, 12, 14, 20]. We also apply gene-*ε* to the SNP-level summary statistics of six quantitative traits assayed in individuals of European ancestry from the UK Biobank [21].

## Results

### Overview of gene-*ε*

The gene-*ε* framework requires two inputs: GWA SNP-level effect size estimates, and an empirical linkage disequilibrium (LD, or variance-covariance) matrix. The LD matrix can be estimated directly from genotype data, or from an ancestry-matched set of samples if genotype data are not available to the user. We use these inputs to both estimate gene-level contributions to narrow-sense heritability *h*^2^, and perform gene-level enrichment tests. After preparing the input data, there are three steps implemented in gene-*ε*, which are detailed below (Fig. 2).

**Figure 2.**
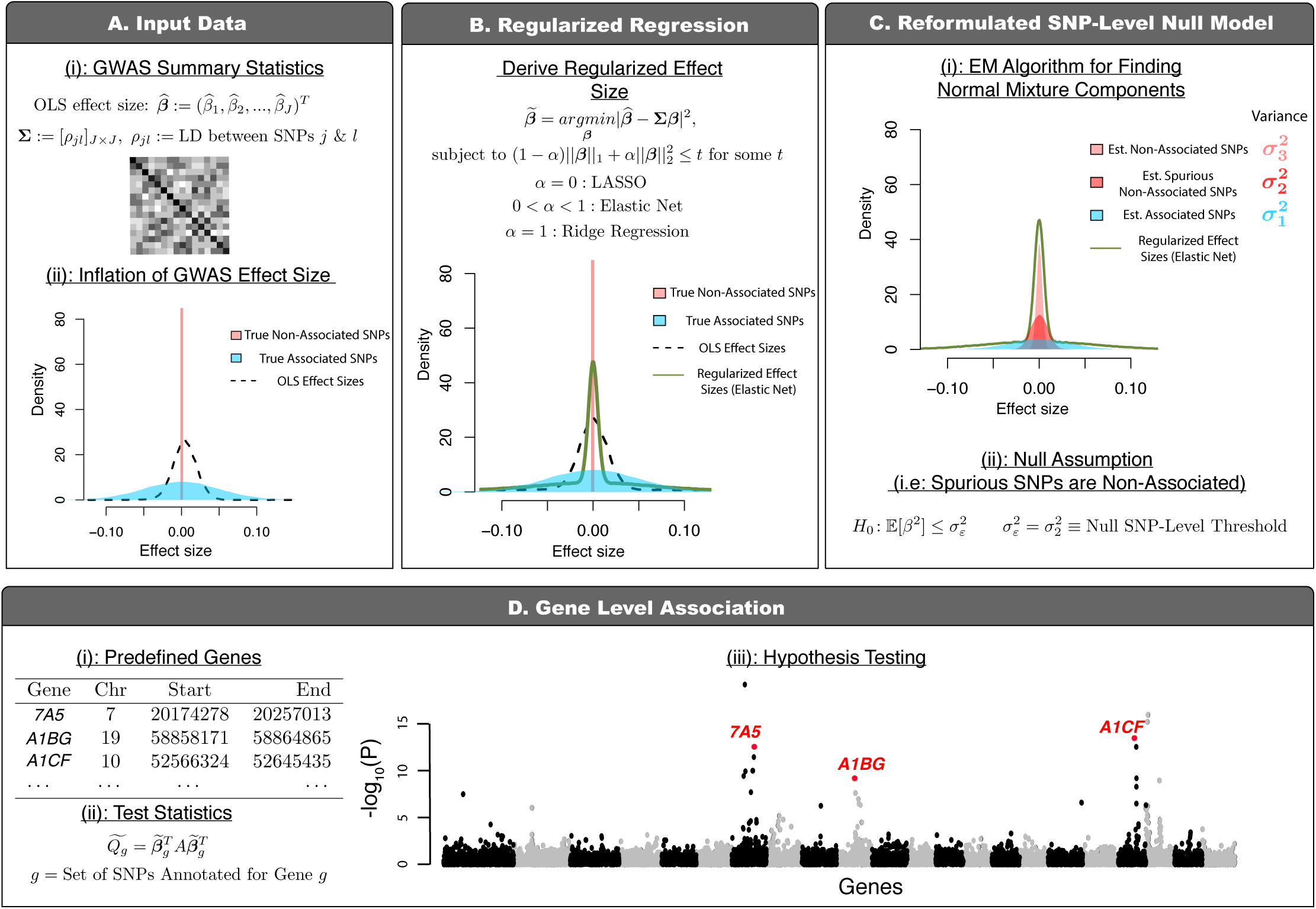
Schematic overview of gene-*ε*: our new gene-level association approach accounting for spurious nonzero SNP-level effects. **(A)** gene-*ε* takes SNP-level GWA marginal effect sizes (OLS estimates 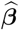) and a linkage disequilibrium (LD) matrix (**Σ**) as input. It is well-known that OLS effect size estimates are inflated due to LD (i.e., correlation structures) among genome-wide genotypes. **(B)** gene-*ε* first uses its inputs to derive regularized effect size estimates 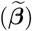 through shrinkage methods (LASSO, Elastic Net and Ridge Regression; we explore performance of each solution under a variety of simulated trait architectures in Supporting Information). **(C)** A unique feature of gene-*ε* is that it treats SNPs with spurious nonzero effects as non-associated. gene-*ε* assumes a reformulated null distribution of SNP-level effects 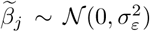, where 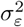 is the SNP-level null threshold and represents the maximum proportion of phenotypic variance explained (PVE) by a spurious or non-associated SNP. This leads to the reformulated SNP-level null hypothesis 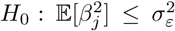. To infer an appropriate 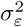, gene-*ε* fits a *K*-mixture of normal distributions over the regularized effect sizes with successively smaller variances 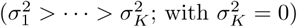. In this study (without loss of generality), we assume that associated SNPs will appear in the first set, while spurious and non-associated SNPs appear in the latter sets. By definition, the SNP-level null threshold is then 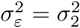. **(D)** Lastly, gene-*ε* computes gene-level association test statistics 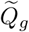 using quadratic forms and corresponding *P* -values using Imhof’s method. This assumes the common gene-level null *H*_0_ : *Q*_*g*_ = 0, where the null distribution of *Q*_*g*_ is dependent upon the SNP-level null threshold 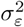. For more details, see Materials and Methods.

First, we shrink the observed GWA effect size estimates via regularized regression (Figs. 2A and B; Eq. (4) in Materials and Methods). This shrinkage step reduces the inflation of OLS effect sizes for spurious SNPs [22], and increases their correlation with the assumed generative model for the trait of interest (particularly for traits with high heritability; Fig. S1). When assessing the performance of gene-*ε* in simulations, we considered different types of regularization for the effect size estimates: the Least Absolute Shrinkage And Selection Operator (gene-*ε*-LASSO) [23], the Elastic Net solution (gene-*ε*-EN) [24], and Ridge Regression (gene-*ε*-RR) [25]. We also assessed our framework using the observed ordinary least squares (OLS) estimates without any shrinkage (gene-*ε*-OLS) to serve as motivation for having regularization as a step in the framework.

Second, we fit a *K*-mixture Gaussian model to all regularized effect sizes genome-wide with the goal of classifying SNPs as associated, non-associated with spurious statistical signal, or non-associated with zero-effects (Figs. 1B and 2C; see also [18]). Each successive Gaussian mixture component has distinctly smaller variances 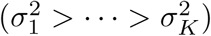 with the *K*-th component fixed at 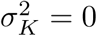. Estimating these variance components helps determine an appropriate *k*-th category to serve as the cutoff for SNPs with null effects (i.e., choosing some variance component 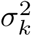 to be the null threshold 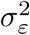). The gene-*ε* software allows users to determine this cutoff subjectively. Intuitively, enriched genes are likely to contain important variants with relatively larger effects that are categorized in the early-to-middle mixture components. Since the biological interpretation of the middle components may not be consistent across trait architectures, we take a conservative approach in our selection of a cutoff when determining associated SNPs. Without loss of generality, we assume non-null SNPs appear in the first mixture component with the largest variance, while null SNPs appear in the latter components. By this definition, non-associated SNPs with spurious *ε*-genic or zero-effects then have PVEs that fall at or below the variance of the second component (i.e., 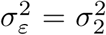 and 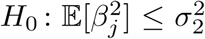 for the *j*-th SNP). gene-*ε* allows for flexibility in the number of Gaussians that specify the range of null and non-null SNP effects. To achieve genome-wide scalability, we estimate parameters of the *K*-mixture model using an expectation-maximization (EM) algorithm.

Third, we group the regularized GWA summary statistics according to gene boundaries (or user-specified SNP-sets) and compute a gene-level enrichment statistic based on a commonly used quadratic form (Fig. 2D) [7, 12, 20]. In expectation, these test statistics can be naturally interpreted as the contribution of each gene to the narrow-sense heritability. We use Imhof’s method [26] to derive a *P* -value for assessing evidence in support of an association between a given gene and the trait of interest. Details for each of these steps can be found in Materials and Methods, as well as in Supporting Information.

### Performance Comparisons in Simulation Studies

To assess the performance of gene-*ε*, we simulated complex traits under multiple genetic architectures using real genotype data on chromosome 1 from individuals of European ancestry in the UK Biobank (Materials and Methods). Following quality control procedures, our simulations included 36,518 SNPs (Supporting Information). Next, we used the NCBI’s Reference Sequence (RefSeq) database in the UCSC Genome Browser [27] to annotate SNPs with the appropriate genes. Simulations were conducted using two different SNP-to-gene assignments. In the first, we directly used the UCSC annotations which resulted in 1,408 genes to be used in the simulation study. In the second, we augmented the UCSC gene boundaries to include SNPs within ±50kb, which resulted in 1,916 genes in the simulation study. For both cases, we assumed a linear additive model for quantitative traits, while varying the following parameters: sample size (*N* = 5,000 or 10,000); narrow-sense heritability (*h*^2^ = 0.2 or 0.6); and the percentage of enriched genes (set to 1% or 10%). In each scenario, we considered traits being generated with and without additional population structure. In the latter setting, traits are simulated while also using the top ten principal components of the genotype matrix as covariates to create stratification. Regardless of the setting, GWA summary statistics were computed by fitting a single-SNP univariate linear model (via OLS) without any control for population structure. Comparisons were based on 100 different simulated runs for each parameter combination.

We compared the performance of gene-*ε* against that of five competing gene-level association or enrichment methods: SKAT [20], VEGAS [7], MAGMA [10], PEGASUS [12], and RSS [14] (Supporting Information). As previously noted, we also explored the performance of gene-*ε* while using various degrees of regularization on effect size estimates, with gene-*ε*-OLS being treated as a baseline. SKAT, VEGAS, and PEGASUS are frequentist approaches, in which SNP-level GWA *P* -values are drawn from a correlated chi-squared distribution with covariance estimated using an empirical LD matrix [28]. MAGMA is also a frequentist approach in which gene-level *P* -values are derived from distributions of SNP-level effect sizes using an *F* -test [10]. RSS is a Bayesian model-based enrichment method which places a likelihood on the observed SNP-level GWA effect sizes (using their standard errors and LD estimates), and assumes a spike-and-slab shrinkage prior on the true SNP effects [29]. Conceptually, SKAT, MAGMA, VEGAS, and PEGASUS assume null models under the traditional GWA framework, while RSS and gene-*ε* allow for traits to have architectures with more complex SNP effect size distributions.

For all methods, we assess the power and false discovery rates (FDR) for identifying correct genes at a Bonferroni-corrected threshold (*P* = 0.05*/*1408 genes = 3.55 × 10^−5^ and *P* = 0.05*/*1916 genes = 2.61 × 10^−5^, depending on if the ±50kb buffer was used) or median probability model (posterior enrichment probability > 0.5; see [30]) (Tables S1-S16). We also compare their ability to rank true positives over false positives via receiver operating characteristic (ROC) and precision-recall curves (Figs. 3 and S2-S16). While we find gene-*ε* and RSS have the best tradeoff between true and false positive rates, RSS does not scale well for genome-wide analyses (Table 1). In many settings, gene-*ε* has similar power to RSS (while maintaining a considerably lower FDR), and generally outperforms RSS in precision-versus-recall. gene-*ε* also stands out as the best approach in scenarios where the observed OLS summary statistics were produced without first controlling for confounding stratification effects in more heritable traits (i.e., *h*^2^ = 0.6). Computationally, gene-*ε* gains speed by directly assessing evidence for rejecting the gene-level null hypothesis, whereas RSS must compute the posterior probability of being an enriched gene (which can suffer from convergence issues; Supporting Information). For context, an analysis of just 1,000 genes takes gene-*ε* an average of 140 seconds to run on a personal laptop, while RSS takes around 9,400 seconds to complete.

**Table 1.**
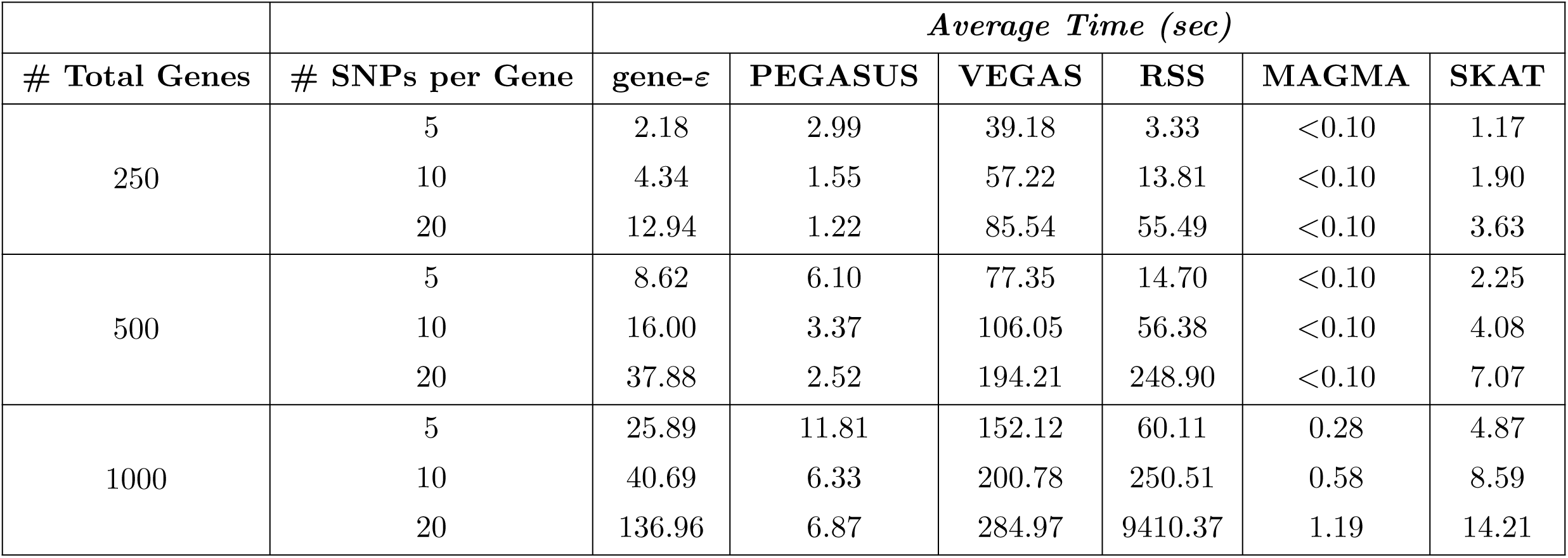
Computational time for running gene-*ε* and other gene-level association approaches, as a function of the total number genes analyzed and the number of SNPs within each gene. Methods compared include: gene-*ε*, PEGASUS [12], VEGAS [7], RSS [14], MAGMA [10], and SKAT [20]. Here, we simulated 10 datasets for each pair of parameter values (number of genes analyzed, and number of SNPs within each gene). Each table entry represents the average computation time (in seconds) it takes each approach to analyze a dataset of the size indicated. Run times were measured on a MacBook Pro (Processor: 3.1-gigahertz (GHz) Intel Core i5, Memory: 8GB 2133-megahertz (MHz) LPDDR3). Only a single core on the machine was used. PEGASUS, SKAT, and MAGMA are score-based methods and, thus, are expected to take the least amount of time to run. Both gene-*ε* and RSS are regression-based methods, but gene-*ε* is scalable in both the number of genes and the number of SNPs per gene. The increased computational burden of RSS results from its need to do Bayesian posterior inference; however, gene-*ε* is able to scale because it leverages regularization and point estimation for hypothesis testing.

| # Total Genes | # SNPs per Gene | *Average Time (sec)* |  |  |  |  |  |
| --- | --- | --- | --- | --- | --- | --- | --- |
| | | gene-ε | PEGASUS | VEGAS | RSS | MAGMA | SKAT |
| 250 | 5 | 2.18 | 2.99 | 39.18 | 3.33 | <0.10 | 1.17 |
|  | 10 | 4.34 | 1.55 | 57.22 | 13.81 | <0.10 | 1.90 |
|  | 20 | 12.94 | 1.22 | 85.54 | 55.49 | <0.10 | 3.63 |
| 500 | 5 | 8.62 | 6.10 | 77.35 | 14.70 | <0.10 | 2.25 |
|  | 10 | 16.00 | 3.37 | 106.05 | 56.38 | <0.10 | 4.08 |
|  | 20 | 37.88 | 2.52 | 194.21 | 248.90 | <0.10 | 7.07 |
| 1000 | 5 | 25.89 | 11.81 | 152.12 | 60.11 | 0.28 | 4.87 |
|  | 10 | 40.69 | 6.33 | 200.78 | 250.51 | 0.58 | 8.59 |
|  | 20 | 136.96 | 6.87 | 284.97 | 9410.37 | 1.19 | 14.21 |

**Figure 3.**
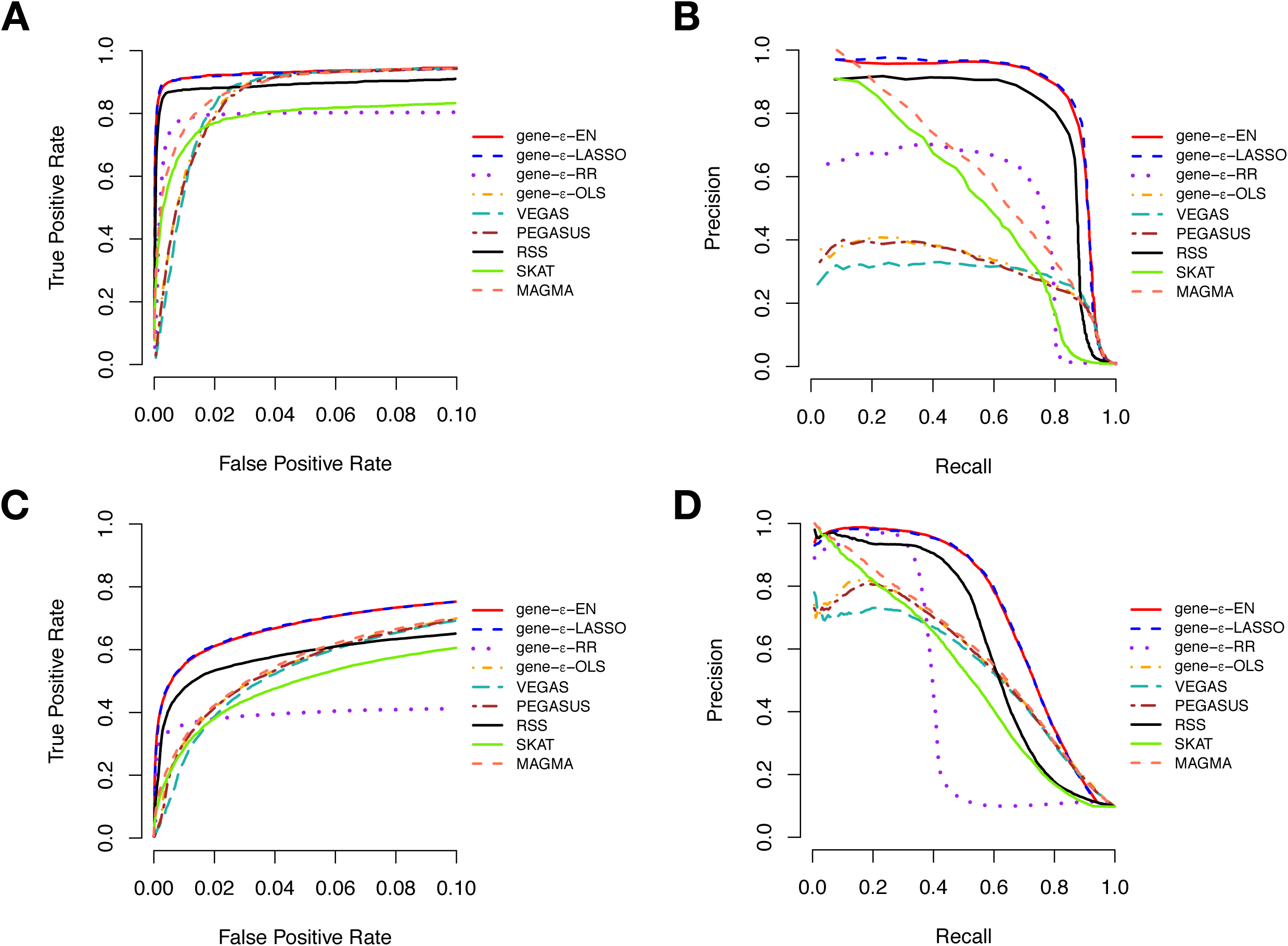
Receiver operating characteristic (ROC) and precision-recall curves comparing the performance of gene-*ε* and competing approaches in simulations (*N* = 10, 000; *h*^*2*^ = 0.6). We simulate complex traits under different genetic architectures and GWA study scenarios, varying the following parameters: narrow sense heritability, proportion of associated genes, and sample size (Supporting Information). Here, the sample size *N* = 10, 000 and the narrow-sense heritability *h*^2^ = 0.6. We compute standard GWA SNP-level effect sizes (estimated using ordinary least squares). Results for gene-*ε* are shown with LASSO (blue), Elastic Net (EN; red), and Ridge Regression (RR; purple) regularizations. We also show the results of gene-*ε* without regularization to illustrate the importance of this step (labeled OLS; orange). We further compare gene-*ε* with five existing methods: PEGASUS (brown) [12], VEGAS (teal) [7], the Bayesian approach RSS (black) [14], SKAT (green) [20], and MAGMA (peach) [10]. **(A, C)** ROC curves show power versus false positive rate for each approach of sparse (1% associated genes) and polygenic (10% associated genes) architectures, respectively. Note that the upper limit of the x-axis has been truncated at 0.1. **(B, D)** Precision-Recall curves for each method applied to the simulations. Note that, in the sparse case (1% associated genes), the top ranked genes are always true positives, and therefore the minimal recall is not 0. All results are based on 100 replicates.

When using GWA summary statistics to identify genotype-phenotype associations, modeling the appropriate trait architecture is crucial. As expected, all methods we compared in this study have relatively more power for traits with high *h*^2^. However, our simulation studies confirm the expectation that the max utility for methods assuming the traditional GWA framework (i.e., SKAT, MAGMA, VEGAS, and PEGASUS) is limited to scenarios where heritability is low, phenotypic variance is dominated by just a few enriched genes with large effects, and summary statistics are not confounded by population structure (Figs. S2, S3, S9, and S10). RSS, gene-*ε*-EN, and gene-*ε*-LASSO robustly outperform these methods for the other trait architectures (Figs. 3, S4-S8, and S11-S16). One major reason for this result is that shrinkage and penalized regression methods appropriately correct for inflation in GWA summary statistics (Fig. S1). For example, we find that the regularization used by gene-*ε*-EN and gene-*ε*-LASSO is able to recover effect size estimates that are almost perfectly correlated (*r*^2^ > 0.9) with the true effect sizes used to simulate sparse architectures (e.g., simulations with 1% enriched genes). In Figs. S17-S24, we show a direct comparison between gene-*ε* with and without regularization to show how inflated SNP-level summary statistics directly affect the ability to identify enriched genes across different trait architectures. Regularization also allows gene-*ε* to preserve type 1 error when traits are generated under the null hypothesis of no gene enrichment. Importantly, our method is relatively conservative when GWA summary statistics are less precise and derived from studies with smaller sample sizes (e.g., *N* = 5,000; Table S17).

### Characterizing Genetic Architecture of Quantitative Traits in the UK Biobank

We applied gene-*ε* to 1,070,306 genome-wide SNPs and six quantitative traits — height, body mass index (BMI), mean red blood cell volume (MCV), mean platelet volume (MPV), platelet count (PLC), waisthip ratio (WHR) — assayed in 349,414 European-ancestry individuals in the UK Biobank (Supporting Information) [21]. After quality control, we regressed the top ten principal components of the genotype data onto each trait to control for population structure, and then we derived OLS SNP-level effect sizes using the traditional GWA framework. For completeness, we then analyzed these GWA effect size estimates with the four different implementations of gene-*ε*. In the main text, we highlight results under the Elastic Net solution; detailed findings with the other gene-*ε* approaches can be found in Supporting Information.

While estimating *ε*-genic effects, gene-*ε* provides insight into to the genetic architecture of a trait (Table S18). For example, past studies have shown human height to have a higher narrow-sense heritability (estimates ranging from 45-80%; [6, 31–39]). Using Elastic Net regularized effect sizes, gene-*ε* estimated approximately 11% of SNPs in the UK Biobank to be statistically associated with height. This meant approximately 110,000 SNPs had marginal PVEs 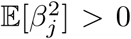 (Materials and Methods). This number is similar to the 93,000 and 100,000 height associated variants previously estimated by Goldstein [40] and Boyle et al. [4], respectively. Additionally, gene-*ε* identified approximately 2% of SNPs to be “causal” (meaning they had PVEs greater than the SNP-level null threshold, 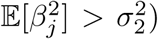; again similar to the Boyle et al. [4] estimate of 3.8% causal SNPs for height using data from the GIANT Consortium [32], and the Lello et al. [41] estimate of 3.1% causal SNPs for height using European-ancestry individuals in the UK Biobank.

Compared to body height, narrow-sense heritability estimates for BMI have been considered both high and low (estimates ranging from 25-60%; [31, 33, 34, 36, 37, 39, 42–45]). Such inconsistency is likely due to difference in study design (e.g., twin, family, population-based studies), many of which have been known to produce different levels of bias [44]. Here, our results suggest BMI to have a lower narrow-sense heritability than height, with a slightly different distribution of null and non-null SNP effects. Specifically, we found BMI to have 13% associated SNPs and 6% causal SNPs.

In general, we found our genetic architecture characterizations in the UK Biobank to reflect the same general themes we saw in the simulation study. Less aggressive shrinkage approaches (e.g., OLS and Ridge) are subject to misclassifications of associated, spurious, and non-associated SNPs. As a result, these methods struggle to reproduce well-known narrow-sense heritability estimates from the literature, across all six traits. This once again highlights the need for computational frameworks that are able to appropriately correct for inflation in summary statistics.

### gene-*ε* Identifies Refined List of Genetic Enrichments

Next, we applied gene-*ε* to the summary statistics from the UK Biobank and generated genome-wide gene-level association *P* -values (panels A and B of Figs. 4 and S25-S29). As in the simulation study, we conducted two separate analyses using two different SNP-to-gene annotations: *(i)* we used the RefSeq database gene boundary definitions directly, or *(b)* we augmented the gene boundaries by adding SNPs within a ±50 kilobase (kb) buffer to account for possible regulatory elements. A total of 14,322 genes were analyzed when using the UCSC boundaries as defined, and a total of 17,680 genes were analyzed when including the 50kb buffer. The ultimate objective of gene-*ε* is to identify enriched genes, which we define as containing at least one associated SNP and achieving a gene-level association *P* -value below a Bonferroni-corrected significance threshold (in our two analyses, *P* = 0.05*/*14322 genes = 3.49 × 10^−6^ and *P* = 0.05*/*17680 genes 2.83 × 10^−6^, respectively; Tables S19-S24). As a validation step, we compared gene-*ε P* -values to RSS posterior enrichment probabilities for each gene. We also used the gene set enrichment analysis tool Enrichr [46] to identify dbGaP categories with an overrepresentation of significant genes reported by gene-*ε* (panels C and D of Figs. 4 and S25-S29). A comparison of gene-level associations and gene set enrichments between the different gene-*ε* approaches are also listed (Tables S25-S27).

**Figure 4.**
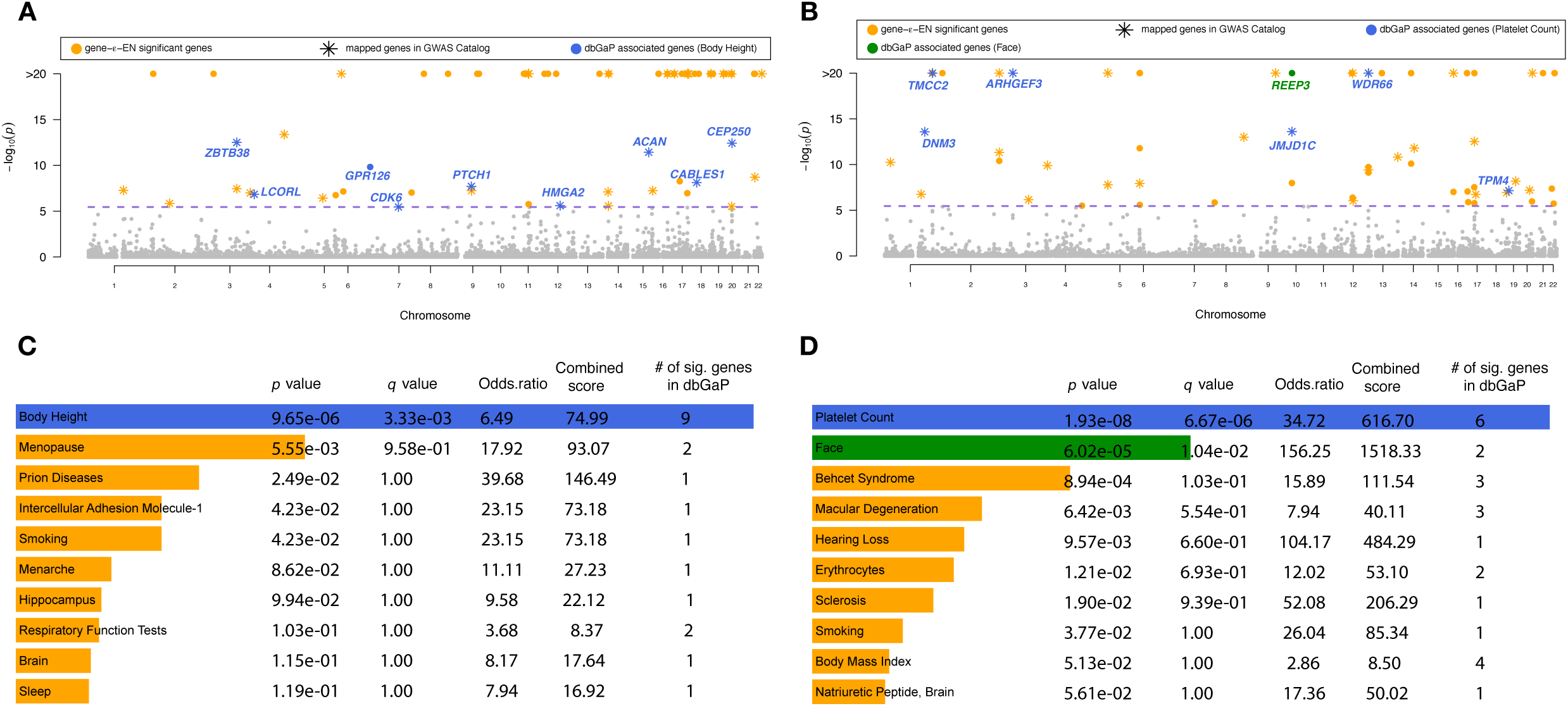
Gene-level association results from applying gene-*ε* to body height (panels A and C) and mean platelet volume (MPV; panels B and D), assayed in European-ancestry individuals in the UK Biobank. Body height has been estimated to have a narrow-sense heritability *h*^2^ in the range of 0.45 to 0.80 [6, 31–39]; while, MPV has been estimated to have *h*^2^ between 0.50 and [33, 34, 90]. Manhattan plots of gene-*ε* gene-level association *P* -values using Elastic Net regularized effect sizes for **(A)** body height and **(B)** MPV. The purple dashed line indicates a log-transformed Bonferroni-corrected significance threshold (*P* = 3.49 × 10^−6^ correcting for 14,322 autosomal genes analyzed). We color code all significant genes identified by gene-*ε* in orange, and annotate genes overlapping with the database of Genotypes and Phenotypes (dbGaP). In **(C)** and **(D)**, we conduct gene set enrichment analysis using Enrichr [46, 91] to identify dbGaP categories enriched for significant gene-level associations reported by gene-*ε*. We highlight categories with *Q*-values (i.e., false discovery rates) less than 0.05 and annotate corresponding genes in the Manhattan plots in **(A)** and **(B)**, respectively. For height, the only significant dbGAP category is “Body Height”, with nine of the genes identified by gene-*ε* appearing in this category. For MPV, the two significant dbGAP categories are “Platelet Count” and “Face” — the first of which is directly connected to trait [57, 92, 93].

Many of the candidate enriched genes we identified by applying gene-*ε* were not previously annotated as having trait-specific associations in either dbGaP or the GWAS catalog (Fig. 4); however, many of these same candidate genes have been identified by past publications as related to the phenotype of interest (Table 2). It is worth noting that multiple genes would not have been identified by standard GWA approaches since the top SNP in the annotated region had a marginal association below a genome-wide threshold (see Table 2 and highlighted rows in Tables S19-S24). Additionally, 45% of the genes selected by gene-*ε* were also selected by RSS. For example, gene-*ε* reports *C1orf150* as having a significant gene-level association with MPV (*P* = 1 × 10^−20^ and RSS posterior enrichment probability of 1), which is known to be associated with germinal center signaling and the differentiation of mature B cells that mutually activate platelets [47–49]. Importantly, nearly all of the genes reported by gene-*ε* had evidence of overrepresentation in gene set categories that were at least related to the trait of interest. As expected, the top categories with Enrichr *Q*-values smaller than 0.05 for height and MPV were “Body Height” and “Platelet Count”, respectively. Even for the less heritable MCV, the top significant gene sets included hematological categories such as “Transferrin”, “Erythrocyte Indices”, “Hematocrit”, “Narcolepsy”, and “Iron” — all of which have verified and clinically relevant connections to trait [50–57].

**Table 2.** Top three newly identified candidate genes reported by gene-*ε* for the six quantitative traits studied in the UK Biobank (using imputed genotypes with gene boundaries defined by the NCBI’s RefSeq database in the UCSC Genome Browser [27]). We call these novel candidate genes because they are not listed as being associated with the trait of interest in either the GWAS catalog or dbGaP, and they have top posterior enrichment probabilities with the trait using RSS analysis. Each gene is annotated with past functional studies that link them to the trait of interest. We also report each gene’s overall trait-specific significance rank (out of 14,322 autosomal genes analyzed for each trait), as well as their heritability estimates from gene-*ε* using Elastic Net to regularize GWA SNP-level effect size estimates. The traits are: height; body mass index (BMI); mean corpuscular volume (MCV); mean platelet volume (MPV); platelet count (PLC); and waist-hip ratio (WHR). ^♣^: Enriched genes whose top SNP is not marginally significant according to a genome-wide Bonferroni-corrected threshold (*P* = 4.67 × 10^−8^ correcting for 1,070,306 SNPs analyzed; see highlighted rows in Supplementary Tables S19-S24 for complete list). *: Multiple genes were tied for this ranking.

| Trait | Gene | Chr | gene-ε $P$-Value | Rank | $h_g^2$ | Post. Prob. | Biological Relevance to Trait | Ref(s) |
| --- | --- | --- | --- | --- | --- | --- | --- | --- |
| Height | *EZH2* | 7 | $9.34 \times 10^{-8}$ | 61 | $7.23 \times 10^{-3}$ | 1.000 | Associated with diseases Adamantinoma of Long Bone and Weaver Syndrome (characterized by rapid growth). | [94] |
| Height | *C17orf42* | 17 | $5.38 \times 10^{-9}$ | 52 | $4.54 \times 10^{-3}$ | 1.000 | Known as the transcription elongation factor of mitochondria (TEFM) which regulates transcription and can affect body height. | [95] |
| Height | *KISS1R* | 19 | $1 \times 10^{-20}$ | 1* | $5.27 \times 10^{-4}$ | 0.970 | Associated with disorders of puberty and final height. | [96] |
| BMI | *ZC3H4* | 19 | $1.62 \times 10^{-14}$ | 20 | $7.84 \times 10^{-3}$ | 1.000 | BMI-inducer known to be associated with adiposity and obesity. | [97–100] |
| BMI | *PTOV1* | 19 | $1 \times 10^{-20}$ | 1* | $2.26 \times 10^{-3}$ | 0.990 | Found to be overexpressed in prostate adenocarcinomas which can be induced by obesity. | [101] |
| BMI | *FBXO45*♣ | 3 | $6.52 \times 10^{-7}$ | 23 | $1.82 \times 10^{-3}$ | 0.029 | Reported to be involved in children syndromic obesity. | [102] |
| MCV | *SLC24A1* | 15 | $1.74 \times 10^{-7}$ | 50 | $4.66 \times 10^{-3}$ | 0.140 | Encoded protein is involved in glucose transportation pathway and MCV is reported to be associated with glucose level. | [101] |
| MCV | *PDX1*♣ | 13 | $1 \times 10^{-20}$ | 1* | $2.31 \times 10^{-4}$ | 0.019 | Associated with Glycated hemoglobin which is affected by MCV | [103] |
| MCV | *RHOD* | 11 | $1 \times 10^{-20}$ | 1* | $3.35 \times 10^{-4}$ | 0.002 | Associated with Wiskott-Aldrich Syndrome which is characterized by abnormal immune system function (immune deficiency) and a reduced ability to form blood clots. | [101, 104] |
| MPV | *C1orf150* | 1 | $1 \times 10^{-20}$ | 1* | $3.44 \times 10^{-2}$ | 1.000 | Known as *GCSAML* which is involved with germinal center signaling and differentiation of mature B cells that mutually activate platelets. | [47–49] |
| MPV | *KIAA0922* | 4 | $3.20 \times 10^{-6}$ | 64 | $7.17 \times 10^{-3}$ | 1.000 | Known as *TMEM131L* which is associated with canonical Wnt signaling and can effect platelet formation. | [105, 106] |
| MPV | *TPT1*♣ | 13 | $1 \times 10^{-20}$ | 1* | $3.25 \times 10^{-4}$ | 0.051 | mRNA expression is identified in platelets. | [101] |
| PLC | *C1orf150* | 1 | $1 \times 10^{-20}$ | 1* | $2.51 \times 10^{-2}$ | 1.000 | Known as *GCSAML* which is involved with germinal center signaling and differentiation of mature B cells that mutually activate platelets. | [47–49] |
| PLC | *PSMD2* | 3 | $1.42 \times 10^{-9}$ | 29 | $7.40 \times 10^{-3}$ | 1.000 | Also known as the 26S proteasome which is found to be important for platelet production. | [101] |
| PLC | *APOB48R* | 16 | $1 \times 10^{-20}$ | 1* | $1.36 \times 10^{-3}$ | 0.003 | Involved in Lipoprotein metabolism pathway which can affect platelet. | [101] |
| WHR | *TFAP2B* | 6 | $3.92 \times 10^{-7}$ | 21 | $3.60 \times 10^{-3}$ | 1.000 | Dietary protein associated with weight maintenance. | [99, 107] |
| WHR | *WDR68* | 17 | $1.05 \times 10^{-7}$ | 20 | $1.10 \times 10^{-3}$ | 0.990 | Also known as *DCAF7* which has been shown to bind Huntingtin-associated protein 1 (HAP1) and affect weight. | [108] |
| WHR | *MLL* | 11 | $8.14 \times 10^{-8}$ | 19 | $2.43 \times 10^{-3}$ | 0.940 | Orthologous gene in mice that affects skeleton, body size, and growth. | [99, 109–111] |

Lastly, gene-*ε* also identified genes with rare causal variants. For example, *ZNF628* (which is not mapped to height in the GWAS catalog) was detected by gene-*ε* with a significant *P* -value of 1 × 10^−20^ (and *P* = 4.58 × 10^−8^ when the gene annotation included a 50kb buffer). Previous studies have shown a rare variant *rs147110934* within this gene to significantly affect adult height [38]. Rare and low-frequency variants are generally harder to detect under the traditional GWA framework. However, rare variants have been shown to be important for explaining the variation of complex traits [28, 39, 58–61]. With regularization and testing for spurious *ε*-genic effects, gene-*ε* is able to distinguish between rare variants that are causal and SNPs with larger effect sizes due various types of correlations. This only enhances the power of gene-*ε* to identify potential novel enriched genes.

## Discussion

During the past decade, it has been repeatedly observed that the traditional GWA framework can struggle to accurately differentiate between associated and spurious SNPs (which we define as SNPs that covary with associated SNPs but do not directly influence the trait of interest). As a result, the traditional GWA approach is prone to generating false positives, and detects variant-level associations spread widely across the genome rather than aggregated sets in disease-relevant pathways [4]. While this observation has spurred to many interesting lines of inquiry — such as investigating the role of rare variants in generating complex traits [9, 28, 58, 59], comparing the efficacy of tagging causal variants in different ancestries [62, 63], and integrating GWA data with functional -omics data [64–66] — the focus of GWA studies and studies integrating GWA data with other -omics data is still largely based on the role of individual variants, acting independently.

Here, our objective is to identify biologically significant underpinnings of the genetic architecture of complex traits by modifying the traditional GWA null hypothesis from *H*_0_ : *β*_*j*_ = 0 (i.e., the *j*-th SNP has zero statistical association with the trait of interest) to *H*_0_ : *β*_*j*_ ≈ 0. We accomplish this by testing for *ε*-genic effects: spurious small-to-intermediate effect sizes emitted by truly non-associated SNPs. We use an empirical Bayesian approach to learn the effect size distributions of null and non-null SNP effects, and then we aggregate (regularized) SNP-level association signals into a gene-level test statistic that represents the gene’s contribution to the narrow-sense heritability of the trait of interest. Together, these two steps reduce false positives and increase power to identify the mutations, genes, and pathways that directly influence a trait’s genetic architecture. By considering different thresholds for what constitutes a null SNP effect (i.e., different values of 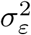 for spurious non-associated SNPs; Figs. 1 and 2), gene-*ε* offers the flexibility to construct an appropriate null hypothesis for a wide range of traits with genetic architectures that land anywhere on the polygenic spectrum. It is important to stress that while we repeatedly point to our improved ability distinguish “causal” variants in enriched genes, gene-*ε* is by no means a causal inference procedure. Instead, it is an association test which highlights genes in enriched pathways that are most likely to be associated with the trait of interest.

Through simulations, we showed the gene-*ε* framework outperforms other widely used gene-level association methods (particularly for highly heritable traits), while also maintaining scalability for genome-wide analyses (Figs. 3 and S2-S24, and Tables 1 and S1-17). Indeed, all the approaches we compared in this study showed improved performance when they used summary statistics derived from studies with larger sample sizes (i.e., simulations with *N* = 10, 000). This is because the quality of summary statistics also improves in these settings (via the asymptotic properties of OLS estimates). Nonetheless, our results suggest that applying gene-*ε* to summary statistics from previously published studies will increase the return made on investments in GWA studies over the last decade.

Like any aggregated SNP-set association method, gene-*ε* has its limitations. Perhaps the most obvious limitation is that annotations can bias the interpretation of results and lead to erroneous scientific conclusions (i.e., might cause us to highlight the “wrong” gene [14, 67, 68]). We observed some instances of this during the UK Biobank analyses. For example, when studying MPV, *CAPN10* only appeared to be a significant gene after its UCSC annotated boundary was augmented by a ±50kb buffer window (*P* = 1.85 × 10^−1^ and *P* = 1.17 × 10^−7^ before and after the buffer was added, respectively; Table S22). After further investigation, this result occurred because the augmented definition of *CAPN10* included nearly all causal SNPs from the significant neighboring gene *RNPEPL1* (*P* = 1 × 10^−20^ and *P* = 2.07 × 10^−9^ before and after the buffer window was added, respectively). While this shows the need for careful biological interpretation of the results, it also highlights the power of gene-*ε* to prioritize true genetic signal effectively.

Another limitation of gene-*ε* is that it relies on the user to determine an appropriate SNP-level null threshold 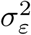 to serve as a cutoff between null and non-null SNP effects. In the current study, we use a *K*-mixture Gaussian model to classify SNPs into different categories and then (without loss of generality) we subjectively assume that associated SNPs only appear in the component with the largest variance (i.e., we choose 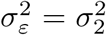). Indeed, there can be many scenarios where this particular threshold choice is not optimal. For example, if there is one very strongly associated locus, the current implementation of the algorithm will assign it to its own mixture component and all other SNPs will be assumed to be not associated with the trait, regardless of the size of their corresponding variances. As previously mentioned, one practical guideline would be to select 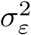 based on some *a priori* knowledge about a trait’s architecture. However, a more robust approach would be to select the SNP-null hypothesis threshold based on the data at hand. One way to do this would be to take a fully Bayesian approach and allow posterior inference on 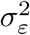 to be dependent upon how much heritability is explained by SNPs placed in the top few largest components of the normal mixture. Recently, sparse Bayesian parametric [69] and nonparametric [70] Gaussian mixture models have been proposed for improved polygenic prediction with summary statistics. Combining these modeling strategies with our modified SNP-level null hypothesis could make for a more unified and data-driven implementation of the gene-*ε* framework.

There are several other potential extensions for the gene-*ε* framework. First, in the current study, we only focused on applying gene-*ε* to quantitative traits (Figs. 4 and S25-S29, and Tables 2 and S18-S27). Future studies extending this approach to binary traits (e.g., case-control studies) should explore controlling for additional confounders that can occur within these phenotypes, such as ascertainment [71–73]. Second, we only focus on data consisting of common variants; however, it would be interesting to extend gene-*ε* for (*i*) rare variant association testing and (*ii*) studies that consider the combined effect between rare and common variants. A significant challenge, in either case, would be to adaptively adjust the strength of the regularization penalty on the observed OLS summary statistics for causal rare variants, so as to not misclassify them as spurious non-associated SNPs. Previous approaches with specific re-weighting functions for rare variants may help here [9, 28, 58] (Materials and Methods). A final related extension of gene-*ε* is to include information about standard errors when estimating *ε*-genic effects. In our analyses using the UK Biobank, some of the newly identified candidate genes contained SNPs that had large effect sizes but insignificant *P* -values in the original GWA analysis (after Bonferroni-correction; Tables 2 and S19-S24). While this could be attributed to the modified SNP-level null distribution assumed by gene-*ε*, it also motivates a regularization model that accounts for the standard error of effect size estimates from GWA studies [14, 22, 29].

## Acknowledgements

The authors would like to thank the Editor, Associate Editor, Doug Speed, and the other two anonymous referees for their constructive comments. The authors also thank Xiang Zhu (Stanford University) for help with the implementation of RSS, as well as Sam Smith (Brown University) for help with the management of the UK Biobank data. This research was conducted using the UK Biobank Resource under Application Number 22419, and part of this research was conducted using computational resources and services at the Center for Computation and Visualization (CCV), Brown University. S. Ramachandran also acknowledges support from a Natural Sciences Fellowship at the Swedish Collegium for Advanced Study (Spring 2019), and by the Erling-Persson Family Foundation and the Knut and Alice Wallenberg Foundation.

## Author Contributions

W.C., S.R., and L.C. conceived the methods. W.C. developed the software and carried out all analyses. W.C., S.R., and L.C. wrote and reviewed the manuscript.

## Materials and Methods

### Traditional Association Tests using Summary Statistics

gene-*ε* requires two inputs: genome-wide association (GWA) marginal effect size estimates 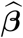, and an empirical linkage disequilibrium (LD) matrix **Σ**. We assumed the following generative linear model for complex traits

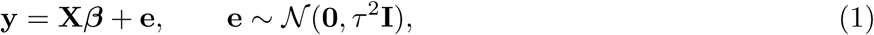

where **y** denotes an *N* -dimensional vector of phenotypic states for a quantitative trait of interest measured in *N* individuals; **X** is an *N* × *J* matrix of genotypes, with *J* denoting the number of single nucleotide polymorphisms (SNPs) encoded as {0, 1, 2} copies of a reference allele at each locus; ***β*** is a *J*-dimensional vector containing the additive effect sizes for an additional copy of the reference allele at each locus on **y**; **e** is a normally distributed error term with mean zero and scaled variance *τ* ^2^; and **I** is an *N* × *N* identity matrix. For convenience, we assumed that the genotype matrix (column-wise) and trait of interest have been mean-centered and standardized. We also treat ***β*** as a fixed effect. A central step in GWA studies is to infer ***β*** for each SNP, given both genotypic and phenotypic measurements for each individual sample. For every SNP *j*, gene-*ε* takes in the ordinary least squares (OLS) estimates based on Eq. (1)

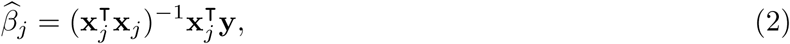

where **x**_*j*_ is the *j*-th column of the genotype matrix **X**, and 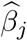 is the *j*-th entry of the vector 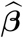. In traditional GWA studies, the null hypothesis for statistical association tests assumes *H*_0_ : *β*_*j*_ = 0 for all *j* = 1, …, *J* SNPs. It can be shown that two genotypic variants **x**_*j*_ and **x**_*j*_ in linkage disequilibrium (LD) will produce effect size estimates 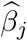 and 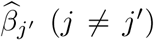 that are correlated [29]. This can lead to confounded statistical tests. For the applications considered here, the LD matrix is empirically estimated from external data (e.g., directly from GWA study data, or using an LD map from a population with similar genomic ancestry to that of the samples analyzed in the GWA study).

### Regularized Regression for GWA Summary Statistics

gene-*ε* uses regularization on the observed GWA summary statistics to reduce inflation of SNP-level effect size estimates and increase their correlation with the assumed generative model of complex traits. For large sample size *N*, note that the asymptotic relationship between the observed GWA effect size estimates 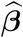 and the true coefficient values ***β*** is [18, 74, 75]

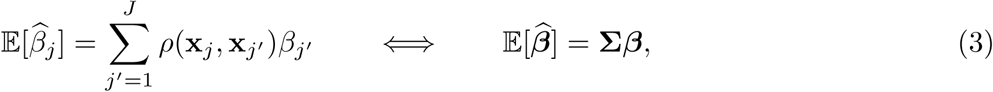

where **Σ**_*jj*′_ = *ρ*(**x**_*j*_, **x**_*j*′_) denotes the correlation coefficient between SNPs **x**_*j*_ and **x**_*j*′_. The above mirrors a high-dimensional regression model with the misestimated OLS summary statistics as the response variables and the LD matrix as the design matrix. Theoretically, the resulting output coefficients from this model are the desired true effect size estimates. Due to the multi-collinear structure of GWA data, we cannot reuse the ordinary least squares solution reliably [76]. Thus, we derive the general regularization

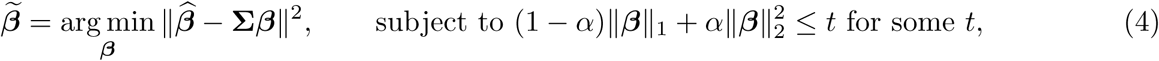

where, in addition to previous notation, the solution 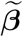 is used to denote the regularized solution of the observed GWA effect sizes 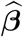; and ‖ • ‖_1_ and 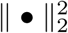 denote *L*_1_ and *L*_2_ penalties, respectively. The free regularization parameter *t* is chosen based off a grid [log *t*_min_, log *t*_max_] with 100 sequential steps of size Here, *t*_max_ is the minimum value such that all summary statistics are shrunk to zero. We then select the *t* that results in a model with an *R*^2^ within one standard error of the best fitted model. In other words, we choose the *t* that (*i*) results in a more sparse solution than the best fitted model, but (*ii*) cannot be distinguished from the best fitted model in terms of overall variance explained.

The term *α* in Eq. (4) distinguishes the type of regularization used, and can be chosen to induce various degrees of shrinkage on the effect size estimates. Specifically, *α* = 0 corresponds to the “Least Absolute Shrinkage and Selection Operator” or LASSO solution [23], *α* = 1 equates to Ridge Regression [25], while 0 < *α* < 1 results in the Elastic Net [24]. The LASSO solution forces some inflated coefficients to be zero; while the Ridge shrinks the magnitudes of all coefficients but does not set any of them to be exactly zero. Intuitively, the LASSO will create a regularized set of effect sizes where associated SNPs have larger effects, non-associated SNPs with spurious small-to-intermediate (or *ε*-genic) effects, and non-associated SNPs with zero-effects. It has been suggested that the *L*_1_-penalty can suffer from a lack of stability [77]. Therefore, in the main text, we also highlighted gene-*ε* using the Elastic Net (with *α* = 0.5). The Elastic Net is a convex combination of the LASSO and Ridge penalties, but still produces distinguishable sets of associated, spurious, and non-associated SNPs. Note that for large GWA studies (e.g., the UK Biobank analysis in the main text), it can be impractical to construct a genome-wide LD matrix; therefore, we regularize OLS effect size estimates based on partitioned chromosome specific LD matrices. Results comparing each of the gene-*ε* regularization implementations are given in the main text (Fig. 3) and Supporting Information (Figs. S2-S24 and Tables S1-18 and 25-27). We will describe how we approximate the null distribution for these regularized GWA summary statistics over the next two sections.

### Estimating the SNP-Level Null Threshold

The main innovation of gene-*ε* is to treat spurious SNPs with *ε*-genic effects as non-associated. This leads to reformulating the GWA SNP-level null hypothesis to assume non-associated SNPs can make small-to-intermediate contributions to the phenotypic variance. Formally, we write this as

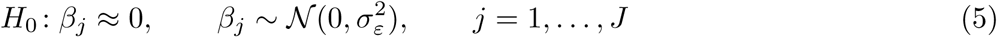

where 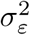 denotes the “SNP-level null threshold” and represents the maximum proportion of phenotypic variance explained (PVE) that is contributed by spurious SNPs. Based on Eq. (5), we equivalently say

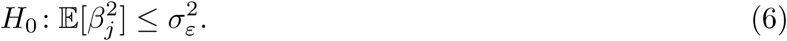

To estimate the threshold 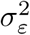 for null SNP-level effects, we use an empirical Bayesian approach and fit a *K*-mixture of normal distributions over the (regularized) effect size estimates [18],

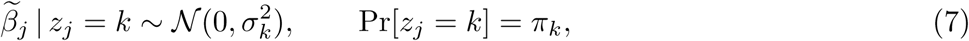

where *z*_*j*_ ∈ {1, …, *K*} is a latent variable representing the categorical membership for the *j*-th SNP. When summing over all components, Eq. (7) corresponds to the following marginal distribution

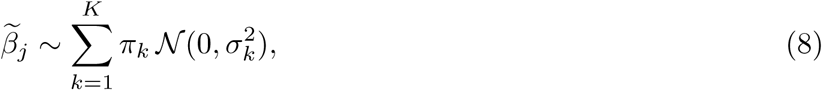

where *π*_*k*_ is a mixture weight representing the marginal (unconditional) probability that a randomly selected SNP belongs to the *k*-th component, with ∑_*k*_ *π*_*k*_ = 1. The above mixture allows for distinct clusters of nonzero effects through *K* different variance components 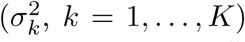 [18]. Here, we consider sequential fractions (*π*_1_, …, *π*_*K*_) of SNPs to correspond to distinctly smaller effects 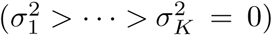 [18]. The goal of the mixture model is to “bin” each of the (regularized) SNP-level effects and determine an appropriate category *k* to serve as the cutoff for SNPs with null effects (i.e., choosing the threshold 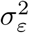 based on some 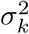). Such a threshold can be chosen based on *a priori* knowledge about the phenotype of interest. It is intuitive to assume that enriched genes will contain non-null SNPs that classify within the early-to-middle mixture components; unfortunately, the biological interpretations of the middle components may not be consistent across trait architectures. Therefore, without loss of generality in this paper, we take a conservative approach in our definition of associated SNPs within enriched genes. Here, we subjectively set the SNP-level null threshold as 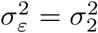. Thus, non-null SNPs are assumed to appear in the largest fraction (i.e., the alternative 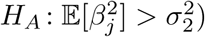, while null SNPs with belong to the latter groups (i.e., the null 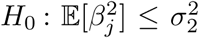. Given Eqs. (7) and (8), we write the joint log-likelihood for all *J* SNPs as the following

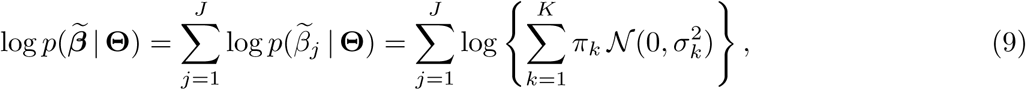

where 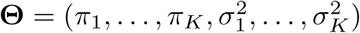 is the complete set of parameters for the mixture model. Since there is not a closed-form solution for the maximum likelihood estimate (MLE), so we use an expectation-maximization (EM) algorithm to estimate the parameters in **Θ** [78–80].

#### Derivation of the EM Algorithm

To derive an EM solution, we use Eqs. (7) and (8) to write the joint distribution of the *J*-regularized SNP-level effect sizes and the *J*-latent random variables **z** = (*z*_1_, …, *z*_*J*_), conditioned on the mixture parameters **Θ**,

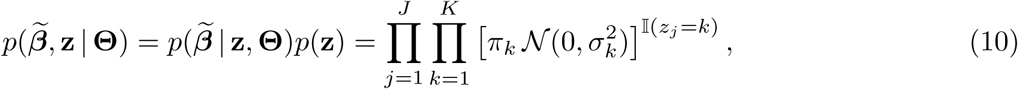

where 𝕀(*z*_*j*_ = *k*) is an indicator function and equates to one if *z*_*j*_ = *k* and zero otherwise. Taking the log of this distribution yields the following

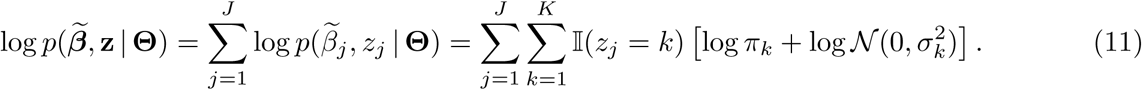

As opposed to Eq. (9), the augmented log-likelihood in Eq. (11) is a much simpler function for which to find a solution. The formal steps of the EM algorithm are now detailed below:

1. **E-Step: Update the Probability of Fraction Assignment.** In the E-step of the EM algorithm, we estimate the probability that the *j*-th SNP belongs to one of the *K* fraction groups. To begin, we use Bayes theorem to find

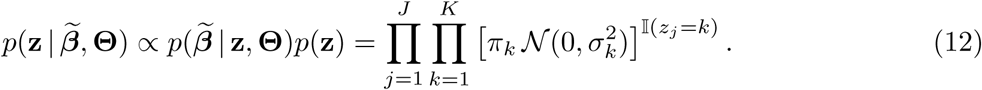 Next, we take the expectation of the complete log-likelihood 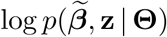, with respect to the condtional distribution 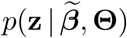, under current value of the mixture parameters 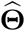. This yields

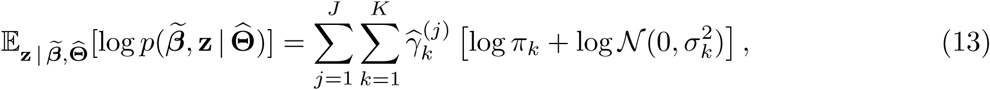

where 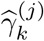 is referred to as the “responsibility of the *k*-th mixture component”, and is given as

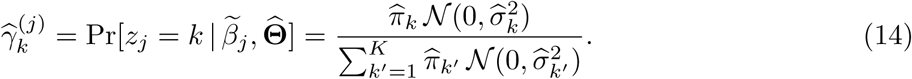 Intuitively, the EM algorithm uses the collection of these responsibility values to assign SNPs to one of the *K* fraction groups. This key step may be interpreted as determining the category of SNP effects (which is determined by identifying the *k*-th component with the largest 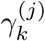 for each *j*-th SNP).
2. **M-Step: Update the Component Variances and Mixture Weights.** In the M-step of the EM algorithm, we now fix the responsibility values and maximize the expectation in Eq. (13), with respect to the parameters in 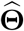. Namely, we compute the following closed-form solutions:

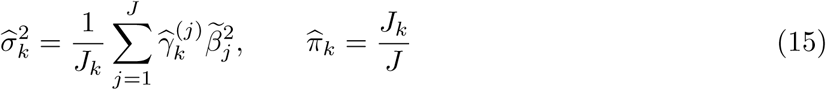

where 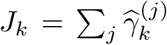 is the sum of the membership weights for the *k*-th mixture component and represents the number of SNPs assigned to that component. The 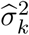 estimates are used to set the SNP-level null threshold 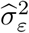.

The gene-*ε* software implements the above EM algorithm using the `mclust` [81] package in R. Results in the main text and Supporting Information are based on 100 iterations from 10 different parallel chains to ensure convergence. To implement the above algorithm, we use the `mclust` software package which can fit a Gaussian mixture with up to *K* = 10 distinct components (see Software Details). Here, the function will compare the Bayesian Information Criterion (BIC) approximation to the Bayes factor for each possible *K* [82], *and produces a resulting output for the K* value that has the largest BIC value. Note that since the EM updates do not involve any large LD matrices, the algorithm scales to be fit efficiently over all SNPs genome-wide.

### Regularized GWA Summary Statistics under the Null Hypothesis

With an estimate of the SNP-level null threshold 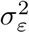, we now describe the probabilistic distribution of the regularized GWA summary statistics under the null hypothesis. Without loss of generality, we demonstrate this property using the general regularization approach where we fix *α* ∈ [0, 1] and have the following (approximate) closed form solution for the regularized effect size estimates [23–25]

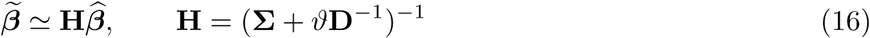

with *ϑ* ≥ 0 being a penalization parameter that has one-to-one correspondence with *t* in Eq. (4). Here, **H** is commonly referred to as the “linear shrinkage estimator” **[citation]**, where **D** is a diagonal weight matrix with nonzero elements dictated by the type of regularization that is being used. For example, **D** = **I** while performing ridge regression [25], and 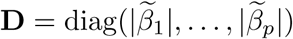 while using ridge-based approximations for the elastic net and lasso solutions [23, 24]. From Eq. (16), it is clear that 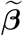 may be interpreted as a marginal estimator of SNP-level effects after accounting for LD structure. Using Eqs. (2)-(3), it is straightforward to show the (approximate) relationship between the regularized effect size estimates and the true coefficient values

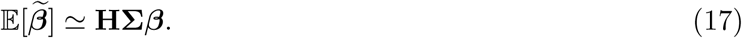

As described in the main text, the accuracy of this relationship is dependent upon both the sample size and narrow-sense heritability of the trait of interest (Fig. S1). Indeed, if **Σ** is full rank and regularization is no longer implemented (i.e., *ϑ* = 0), 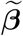 is simply the ordinary least squares solution for marginal GWA summary statistics with asymptotic variance-covariance 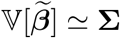 under the null model [18, 74, 75]. In the limiting case where the number of observations in a GWA study is large (i.e., *N* → ∞) and the trait of interest is highly heritable, 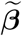 converges onto ***β*** in expectation; and thus is assumed to be independently and normally distributed under the null hypothesis with asymptotic variance 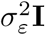 (previously discussed in Eq. (5)). As empirically demonstrated for synthetic traits in the current study, we are rarely in situations where we expect the regularized effect size estimates to have completely converged onto the true generative SNP-level coefficients (again see Fig. S1). This effectively means that we cannot expect each 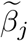 to be completely independent under the null hypothesis in practice. We accommodate this realization by assuming that under the null model

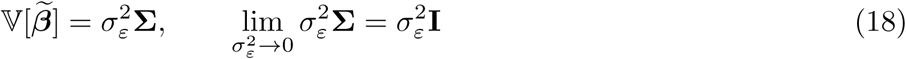

Our reasoning for the formulation above is that, for most quality controlled studies, SNPs in perfect LD will have been pruned such that *ρ*(**x**_*j*_, **x**_*j*′_) < *ρ*(**x**_*j*_, **x**_*j*_) for all *j* = *j*′ variants in the data. Therefore, when traits are generated under the idealized null scenario with large sample sizes and no genetic effects, the estimate of 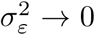 and the off-diagonals of 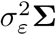 will approach zero quicker than the diagonal elements; thus, allowing the regularized 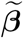 to asymptotically converge onto the true coefficients ***β***. When this scenario does not occur, we are able to appropriately deal with the remaining correlation structure (e.g., all the simulation scenarios explored in this work; see Figs. 3 and S2-S24, and Tables 1 and S1-17).

### Using the SNP-Level Null Threshold to Detect Enriched Genes

We now formalize the hypothesis test for identifying significantly enriched genes conditioned on the SNP-level null threshold 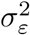, which we compute using the variance component estimates from the EM algorithm detailed in the previous section. The gene-*ε* gene-level test statistic is based on a quadratic form using GWA summary statistics, which is a common approach for generating gene-level test statistics for complex traits. Let gene (or genomic region) *g* represent a known set of SNPs *j* _*g*_ ∈ 𝒥; for example, 𝒥_*g*_ may include SNPs within the boundaries of *g* and/or within its corresponding regulatory region. Here, we conformably partition the regularized GWA effect size estimates 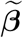 and define the gene-level test statistic

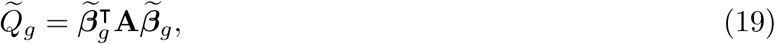

where **A** is an arbitrary symmetric and positive semi-definite weight matrix. We set to **A** = **I** to be the identity matrix for all analyses in the current study; hence, 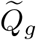 simplifies to a sum of squared SNP effects in the *g*-th gene. Indeed, similar quadratic forms have been implemented to assess the enrichment of mutations at the gene level [7, 12] and across general SNP-sets [9, 20, 28, 58]. A key feature of the gene-*ε* framework is to assess the statistics in Eq. (19) against a gene-level enrichment null hypothesis *H*_0_ : *Q*_*g*_ = 0 that is dependent on the SNP-level null threshold 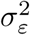. Due to the normality assumption for each SNP effect in Eq. (5), *Q*_*g*_ is theoretically assumed to follow a mixture of chi-square distributions,

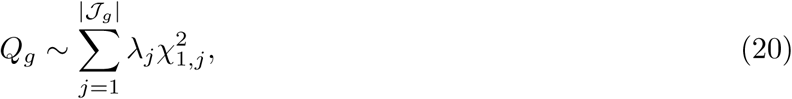

where |𝒥_*g*_| denotes the cardinality of the set of SNPs 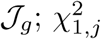 are standard chi-square random variables with one degree of freedom; and 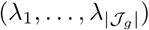 are the eigenvalues of the matrix [83, 84]

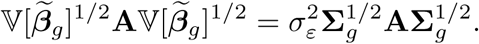

Again, in the current study, 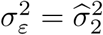 from the estimates in Eq. (15), and **Σ**_*g*_ denotes a subset of the LD matrix only containing SNPs annotated in the *g*-th SNP-set. Again, when **A** = **I**, the eigenvalues are based on a scaled version of the local gene-specific LD matrix. Several approximate and exact methods have been suggested to obtain *P* -values under a mixture of chi-square distributions. In this study, we use Imhof’s method [26] where we empirically compute an estimate of the weighted sum in Eq. (20) and compare this distribution to the observed test statistic in Eq. (19) (see Software Details). It is important to note here that the gene-level null hypothesis is the same for gene-*ε* and other similar competing enrichment methods [9, 12, 20, 28, 58]; the defining characteristic that sets gene-*ε* apart is that it assumes a different null distribution for effects on the SNP-level.

#### Estimating Gene Specific Contributions to the PVE

In the main text, we highlight some of the additional features of the gene-*ε* gene-level association test statistic. First, the expected enrichment for trait-associated mutations in a given gene is equal to the heritability explained by the SNPs contained in said gene. Formally, consider the expansion of Eq. (19) derived from the expectation of quadratic forms,

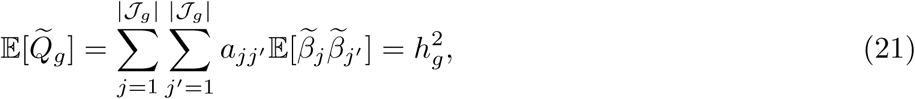

where denotes the heritability contributed by gene *g*. When **A** = **I** (as in the current study), the gene-*ε* hypothesis test for identifying enriched genes is based on the individual SNP contributions to the narrow-sense heritability (i.e., the sum of the expectation of squared SNP effects; see also [34])

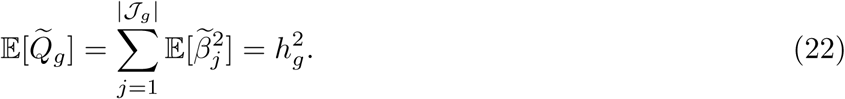

Alternatively, one could choose to re-weight these contributions by specifying **A** otherwise [12, 20, 83, 85, 86]. For example, if SNP *j* has a small effect size but is known to be functionally associated with the trait of interest, then increasing **A**_*jj*_ will reflect this knowledge. Specific weight functions have also been suggested for dealing with rarer variants [9, 28, 58].

### Simulation Studies

We used a simulation scheme to generate SNP-level summary statistics for GWA studies. First, we randomly select a set of enriched genes and assume that complex traits (under various genetic architectures) are generated via a linear model

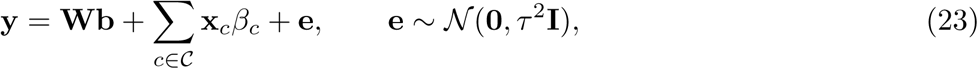

where **y** is an *N* -dimensional vector containing all the phenotypes; 𝒞 represents the set of causal SNPs contained within the associated genes; **x**_*c*_ is the genotype for the *c*-th causal SNP encoded as 0, 1, or 2 copies of a reference allele; *β*_*c*_ is the additive effect size for the *c*-th SNP; **W** is an *N* × *M* matrix of covariates representing additional population structure (e.g., the top ten principal components from the genotype matrix) with corresponding fixed effects **b**; and **e** is an *N* -dimensional vector of environmental noise. The phenotypic variance is assumed 𝕍[**y**] = 1. The effect sizes of SNPs in enriched genes are randomly drawn from standard normal distributions and then rescaled so they explain a fixed proportion of the narrow-sense heritability 𝕍[∑**x**_*c*_*β*_*c*_] = *h*^2^. The covariate coefficients are also drawn from standard normal distributions and then rescaled such that 𝕍[**Wb**] + 𝕍[**e**] = (1 − *h*^2^). GWA summary statistics are then computed by fitting a single-SNP univariate linear model via ordinary least squares (OLS): 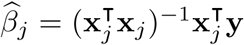 for every SNP in the data *j* = 1,…*J*. These effect size estimates, along with an LD matrix **Σ** computed directly from the full *N* × *J* genotype matrix **X**, are given to gene-*ε*. We also retain standard errors and *P* -values for implementation of the competing methods (VEGAS, PEGASUS, RSS, SKAT, and MAGMA). Given different model parameters, we simulate data mirroring a wide range of genetic architectures (Supporting Information).

## Software Details

Source code implementing gene-*ε* and tutorials are freely available at https://github.com/ramachandran-lab/ genee and was written in `R` (version 3.3.3). Within this software, regularization of the OLS SNP-level effect sizes is done using the package `glmnet` (version 2.0-16) [87]. For large datasets, such as the UK Biobank, the software also offers regularization using the `biglasso` (version 1.3-6) [88] to help with memory and scalability requirements. Note that selection of the free parameter *t* is done the same way using both the `glmnet` and `biglasso` packages. Both packages also take in an *α* ∈ [0, 1] to specify fitting the Ridge, Elastic Net or Lasso regularization to the OLS SNP-level effect sizes. The fitting of a *K*-mixture of Gaussian distributions for the estimation of the SNP-level null threshold 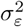 is done using the package `mclust` (version 5.4.3) [81]. Lastly, the package `CompQuadForm` (version 1.4.3) was used to compute gene-*ε* gene-level *P* -values with Imhof’s method [26, 89]. Comparisons in this work were made using software for MAGMA (version 1.07b; https://ctg.cncr.nl/software/magma), PEGASUS (version 1.3.0; https://github.com/ramachandran-lab/PEGASUS), RSS (version 1.0.0; https://github.com/stephenslab/rss), SKAT (version 1.3.2.1; https://www.hsph.harvard.edu/skat), VEGAS (version 2.0.0; https://vegas2.qimrberghofer.edu.au) which are also publicly available. See all other relevant URLs below.

**URLs**

gene-*ε* software, https://github.com/ramachandran-lab/genee; UK Biobank, https://www.ukbiobank.ac.uk; Database of Genotypes and Phenotypes (dbGaP), https://www.ncbi.nlm.nih.gov/gap; NHGRI-EBI GWAS Catalog, https://www.ebi.ac.uk/gwas/; UCSC Genome Browser, https://genome.ucsc.edu/index.html; Enrichr software, http://amp.pharm.mssm.edu/Enrichr/; SNP-set (Sequence) Kernel Association Test (SKAT) software, https://www.hsph.harvard.edu/skat; Multi-marker Analysis of GenoMic Annotation (MAGMA) software, https://ctg.cncr.nl/software/magma; Precise, Efficient Gene Association Score Using SNPs (PEGASUS) software, https://github.com/ramachandran-lab/PEGASUS; Regression with Summary Statistics (RSS) enrichment software, https://github.com/stephenslab/rss; Versatile Gene-based Association Study (VEGAS) version 2, https://vegas2.qimrberghofer.edu.au.

