## Supplementary Text, Figures, and Tables for "Estimation of Non-null SNP Effect Size Distributions Enables the Detection of Enriched Genes Underlying Complex Traits"

### S1 Data Quality Control Procedures

The results presented in the main text made use of imputed data released from the UK Biobank [1]. Quality control procedures for these data are as follows. First, we only studied individuals who self-identified as "white British" people. From this cohort, we further excluded individuals identified by the UK Biobank to have high heterozygosity, excessive relatedness, or aneuploidy (1,550 individuals removed). We also removed individuals whose kinship coefficient was greater than 0.0442 (i.e., close relatives). Next, we removed *(i)* monomorphic SNPs, *(ii)* ambiguous A/T or C/G SNPs, *(iii)* SNPs with minor allele frequency (MAF) less than 2.5%, *(iv)* SNPs not in Hardy-Weinberg Equilibrium (Fisher's exact test $P > 10^{-6}$), *(v)* SNPs with missingness greater than 1%, and *(vi)* SNPs in high linkage disequilibrium (using the flag `--indep-pairwise 50 5 0.9` with PLINK 1.9 [2]). After all QC steps, we had a final dataset of 349,414 individuals and 1,070,306 SNPs. Next, we used the NCBI's Reference Sequence (RefSeq) database in the UCSC Genome Browser [3] to annotate SNPs with the appropriate genes. Recall that in both the simulation studies and real data analysis, we define genes with boundaries in two ways: *(a)* we use the UCSC gene boundary definitions directly, or *(b)* we augment the gene boundaries by adding SNPs within a $\pm 50$ kilobase (kb) buffer to account for possible regulatory elements. Genes with only 1 SNP within their boundary were excluded from either analysis. A total of 14,322 autosomal genes were analyzed when using the UCSC boundaries, and a total of 17,680 autosomal genes were analyzed when including the 50kb buffer.

### S2 Simulation Setup and Scenarios

In our simulation studies, we used the following general simulation scheme to generate SNP-level summary statistics for GWA studies using real genotype data on chromosome 1 from individuals of European ancestry in the UK Biobank [1]. We will denote this genotype matrix as $\mathbf{X}$, with $\mathbf{x}_j$ denoting the genotypic vector for the $j$-th SNP. Following quality control procedures detailed in the previous section, our simulations included $J = 36{,}518$ SNPs distributed across genome. Again, we used the NCBI's RefSeq database in the UCSC Genome Browser to assign SNPs to genes. Simulations were conducted using two different SNP-to-gene assignments. In the first, we directly used the UCSC annotations which resulted in 1,408 genes to be used in the simulation study. In the second, we augmented the UCSC gene boundaries to include SNPs within $\pm 50$kb resulting in 1,916 genes for analysis. Regardless of annotation type, we simulated phenotypes by first assuming that the total phenotypic variance $\mathbb{V}[\mathbf{y}] = 1$ and that all observed genetic effects explained a fixed proportion of this value (i.e., narrow-sense heritability, $h^2$). Next, we randomly selected a certain percentage of enriched genes and denoted the sets of SNPs that they contained as $\mathcal{C}$. Within $\mathcal{C}$, we select causal SNPs in a way such that each associated gene at least contains one SNP with non-zero effect size. Quantitative continuous traits were then generated under the following two general linear models:

(i) Standard Model: $\mathbf{y} = \sum_{c \in \mathcal{C}} \mathbf{x}_c \beta_c + \mathbf{e}$

(ii) Population Stratification Model: $\mathbf{y} = \mathbf{W}\mathbf{b} + \sum_{c \in \mathcal{C}} \mathbf{x}_c \beta_c + \mathbf{e}$

where $\mathbf{y}$ is an $N$-dimensional vector containing all the phenotypes; $\mathbf{x}_c$ is the genotype for the $c$-th causal SNP encoded as 0, 1, or 2 copies of a reference allele; $\beta_c$ is the additive effect size for the $c$-th SNP; and $\mathbf{e} \sim \mathcal{N}(0, \tau^2 \mathbf{I})$ is an $N$-dimensional vector of normally distributed environmental noise. Additionally, in model *(ii)*, $\mathbf{W}$ is an $N \times M$ matrix of the top five principal components (PCs) from the genotype matrix and represents additional population structure with corresponding fixed effects $\mathbf{b}$. The effect sizes of SNPs in enriched genes are randomly drawn from standard normal distributions and then rescaled so they explain a fixed proportion of the narrow-sense heritability $\mathbb{V}[\sum \mathbf{x}_c \beta_c] = h^2$. The coefficients for the genotype PCs are also drawn from standard normal distributions and rescaled such that

$\mathbb{V}[\mathbf{Wb}] = 10\%$ of the total phenotypic variance, with the variance of all non-genetic effects contributing $\mathbb{V}[\mathbf{Wb}]+\mathbb{V}[\mathbf{e}] = (1-h^2)$. For any simulations conducted under model (ii), genotype PCs are not included in any of the model fitting procedures, and no other preprocessing normalizations were carried out to account for the additional population structure. More specifically, GWA summary statistics are then computed by fitting a single-SNP univariate linear model via ordinary least squares (OLS):

$$\widehat{\beta}_j = (\mathbf{x}_j^{\intercal}\mathbf{x}_j)^{-1}\mathbf{x}_j^{\intercal}\mathbf{y}; \tag{S1}$$

for every SNP in the data $j = 1, \ldots J$. These OLS effect size estimates, along with an empirically LD matrix $\boldsymbol{\Sigma}$ computed directly from the full $N \times J$ genotype matrix $\mathbf{X}$, are given to gene-$\varepsilon$ . We also retain standard errors and $P$-values for the implementation of competing methods (i.e., VEGAS, PEGASUS, RSS, SKAT, and MAGMA). Given the simulation procedure above, we simulate a wide range of scenarios for comparing the performance of gene-level association approaches by varying the following parameters:

- Number of individuals: $N$ = 5,000 and 10,000;
- Narrow-sense heritability: $h^2 = 0.2$ and 0.6;
- Percentage of enriched genes: 1% and 10%;

Furthermore, we set the number of causal SNPs with non-zero effects to be some fixed percentage of all SNPs located within the designated enriched genes. In the setting where we have 1,408 genes with boundaries defined strictly by RefSeq in UCSC Genome Browser, we set this percentage to be 0.125% in the 1% associated gene case, and 3% in the 10% associated gene case. In the setting where we have 1,916 genes with boundaries augmented by the ±50kb buffer, we set this percentage to be 0.125% in the 1% associated gene case, and 8% in the 10% associated gene case. Lastly, for each simulated dataset, we also selected some number of intergenic SNPs (i.e., SNPs not mapped to any gene) to have non-zero effect sizes. This was done to mimic genetic associations in unannotated regulatory elements. Specifically, 5 randomly selected intergenic SNPs were given non-zero contributions to the trait heritability in the 1% enriched genes case, and 30 intergenic SNPs were selected in the 10% enriched genes case.

All performance comparisons are based on 100 different simulated runs for each parameter combination. We computed gene-level $P$-values for the gene-$\varepsilon$ approaches, PEGASUS, VEGAS, SKAT, and MAGMA. For evaluting the performance of RSS, we compute posterior enrichment probabilities. For all approaches, we assessed:

- The power and false discovery rates when identifying enriched genes at a Bonferroni-corrected threshold ($P = 0.05/1,408$ genes $= 3.55 \times 10^{-5}$; $P = 0.05/1,916$ genes $= 2.61 \times 10^{-5}$ if the ±50kb buffer was used) or median probability model (posterior enrichment probability $> 0.5$) [4];
- The ability to rank true positive (TP) genes over false positives (FP) via receiver operating characteristic (ROC) and precision-recall curves.

All figures and tables show the mean performances (and standard deviations) across all simulated replicates.

### S3 Review of Other Gene-Level Association Methods

In this section, we give a comprehensive review of the three gene-level association tests that we compare with the gene-$\varepsilon$ approach. To facilitate the understanding of these summaries, we adapt notation from the original references that first introduced these methods to mirror the notation we use in this study.

**Precise, Efficient Gene Association Score Using SNPs (PEGASUS).** Consider a gene $g$ with $|\mathcal{J}_g|$ SNPs, where $|\mathcal{J}_g|$ represents the cardinality of the set of SNPs $\mathcal{J}_g$. Also assume that we have access to corresponding $|\mathcal{J}_g|$ GWA SNP-level $P$-values. We denote the $P$-values for SNPs within a given gene boundary as $\widehat{\mathbf{p}}_g = \{\widehat{p}_1, \ldots, \widehat{p}_{|\mathcal{J}_g|}\}$. PEGASUS computes a gene-level test statistic $\widehat{Q}_g$ via the following quadratic form

$$\widehat{Q}_g = \widehat{\boldsymbol{\beta}}_g^{\intercal}\mathbf{A}\widehat{\boldsymbol{\beta}}_g \tag{S2}$$

where $\widehat{\boldsymbol{\beta}}_g = F^{-1}(\widehat{\mathbf{p}}_g)$, and $F^{-1}(\bullet)$ is the quantile function of the standard chi-square distribution with one degree of freedom, and $\mathbf{A}$ is a predefined symmetric and positive semi-definite weight matrix. Probabilistically, under the null hypothesis, $\widehat{\boldsymbol{\beta}}_g$ is assumed to jointly follow a multivariate normal distribution with mean $\mathbf{0}$ and covariance matrix $\boldsymbol{\Sigma}_g$, where each matrix element $\rho(\mathbf{x}_j, \mathbf{x}_l)$ is the LD between the $j$-th and $l$-th SNPs contained within gene $g$. Therefore, also under the null hypothesis, $Q_g$ is assumed to follow a mixture of chi-square distributions,

$$Q_g \sim \sum_{j=1}^{|\mathcal{J}_g|} \lambda_j U_j^2 \tag{S3}$$

where each $U_j$ is a mutually independent standard normal variables, and $(\lambda_1, \ldots, \lambda_{|\mathcal{J}_g|})$ are the eigenvalues of the matrix product $\boldsymbol{\Sigma}_g\mathbf{A}$. $P$-values are computed numerically using Davies' exact method [5]. See [6] for more details. Note that in our implementation of PEGASUS, $\mathbf{A} = \mathbf{I}$ is set to be the identity matrix.

**Versatile Gene-based Association Study (VEGAS).** Again consider a gene $g$ with $|\mathcal{J}_g|$ SNPs. Under the null hypothesis, a non-associated gene will contain only non-causal SNPs and is assumed to be represented by a $|\mathcal{J}_g|$-dimensional multivariate normal vector $\boldsymbol{\beta}_g^* = (\beta_1^*, \ldots, \beta_{|\mathcal{J}_g|}^*)$ for which

$$\boldsymbol{\beta}_g^* \sim \mathcal{N}(\mathbf{0}, \boldsymbol{\Sigma}_g), \tag{S4}$$

where $\boldsymbol{\Sigma}_g$ is the LD matrix for all SNPs within gene $g$. VEGAS generates gene scores by: *(i)* simulating the random vector $\boldsymbol{\beta}_g^*$ upwards of one million times, *(ii)* transforming the elements of each vector into correlated chi-square variables with one degree of freedom where $q_j = \beta_j^{*2}$ and $\mathbf{Q}_g^* = (q_1, \ldots, q_{|\mathcal{J}_g|})$, *(iii)* acquiring realizations from the null distribution by summing over all the components in each $\mathbf{Q}_g^*$, and *(iv)* computing an empirical gene-level $P$-value based on the proportion of times an observed test statistic is smaller than the simulated null statistics $\Pr[\sum \widehat{\mathbf{Q}}_g < \sum \mathbf{Q}_g^*]$ across all simulations. See [7] for more details.

**Regression with Summary Statistics (RSS) Enrichment.** Consider a GWA study with $N$ individuals typed on $P$ SNPs. For the $j$-th SNP, assume that we are given corresponding effect sizes $\widehat{\beta}_j$ and standard error $\widehat{s}_j$ via a single-SNP linear model fit using OLS. RSS then implements the following likelihood to model the GWA summary statistics [8]

$$\widehat{\boldsymbol{\beta}} \sim \mathcal{N}(\widehat{\mathbf{S}}\boldsymbol{\Sigma}\widehat{\mathbf{S}}^{-1}\boldsymbol{\beta}, \widehat{\mathbf{S}}\boldsymbol{\Sigma}\widehat{\mathbf{S}}) \tag{S5}$$

where $\widehat{\mathbf{S}} = \text{diag}(\widehat{\mathbf{s}})$ is a $J \times J$ diagonal matrix of standard errors, $\boldsymbol{\Sigma}$ is again used to represent some empirical estimate of the LD matrix (i.e., using some external reference panel with ancestry matching the cohort of interest), and $\boldsymbol{\beta}$ are the true (unobserved) SNP-level effect sizes. To model gene-level enrichment, RSS assumes the following hierarchical prior structure on the true effect sizes

$$\beta_j \sim \pi_j\,\mathcal{N}(0, \sigma_\beta^2) + (1 - \pi_j)\,\delta_0, \tag{S6}$$

$$\sigma_\beta^2 = h^2 \left( \sum_{j=1}^{J} \pi_j \, N^{-1} \widehat{s}_j^{-2} \right)^{-1}, \tag{S7}$$

$$\pi_j = \left( 1 + 10^{-(\theta_0 + a_j \theta)} \right)^{-1}, \tag{S8}$$

where $\delta_0$ is point mass centered at zero, $h^2$ denotes the narrow-sense heritability of the trait, $a_j$ is an indicator detailing whether the $j$-th SNP is inside a particular gene, $\theta_0$ is the background proportion of trait-associated SNPs, and $\theta$ reflects the increase in probability (on the $\log_{10}$-odds scale) when a SNP within a gene has non-zero effect. Here, the authors follow earlier works [9] and place independent uniform grid priors on the hyper-parameters $\{h^2, \theta_0, \theta\}$. Note that, unlike other methods, RSS does not calculate a $P$-value for assessing gene-level association. Instead, RSS produces a posterior enrichment probability that at least one SNP in a given gene boundary is associated with the trait

$$P_g := 1 - \Pr\left[\beta_j = 0, \forall j \in \mathcal{J}_g \,|\, \mathbf{D}\right] \tag{S9}$$

where $\mathbf{D}$ represents all of the input data including the GWA summary statistics $\{\widehat{\boldsymbol{\beta}}, \widehat{\mathbf{s}}\}$, the estimated LD matrix $\boldsymbol{\Sigma}$, and any applicable SNP annotations or weights $\mathbf{a} = (a_1, \ldots, a_J)$. See [8, 10] for more details on preferred hyper-parameter settings. As noted in the main text, RSS is relies on a Markov chain Monte Carlo (MCMC) scheme for sampling posterior distributions and estimating model parameters. As a result, its algorithm can be subject to convergence issues if these (or the random seed) are not chosen properly.

**SNP-set (Sequence) Kernel Association Test (SKAT).** The implementation of SKAT required access to raw phenotype $\mathbf{y}$ and genotype $\mathbf{X}$ information for $N$ individuals typed on $J$ SNPs. To assess enrichment of the $|\mathcal{J}_g|$ variants within gene $g$, consider the linear model with sub-matrix $\mathbf{X}_g$

$$\mathbf{y} = \beta_0 + \mathbf{X}_g \boldsymbol{\beta}_g + \mathbf{e}, \qquad \mathbf{e} \sim \mathcal{N}(\mathbf{0}, \tau^2 \mathbf{I}) \tag{S10}$$

where $\beta_0$ is an intercept term, $\boldsymbol{\beta}_g = (\beta_1, \ldots, \beta_{|\mathcal{J}_g|})$ is a vector of regression coefficients for the SNPs within the gene of interest, and $\mathbf{e}$ is a normally distributed error term with mean zero and scaled variance $\tau^2$. For model flexibility, gene-specific SNP effects $\beta_j$ are assumed to follow an arbitrary distribution with mean zero and marginal variances $a_j \sigma_\beta^2$, where $\sigma_\beta^2$ is a variance component and $a_j$ is a pre-specified weight for the $j$-th SNP. To this end, SKAT uses a variance component scoring approach and tests the null hypothesis $H_0 : \boldsymbol{\beta} = \mathbf{0}$, or equivalently $H_0 : \sigma_\beta^2 = 0$. The corresponding gene-level test statistic $\widehat{Q}_g$ then takes on the familiar quadratic form

$$\widehat{Q}_g = (\mathbf{y} - \widehat{\beta}_0)^\intercal \mathbf{K}_g (\mathbf{y} - \widehat{\beta}_0) \tag{S11}$$

where $\widehat{\beta}_0$ is the predicted mean of trait under the null hypothesis, and is computed by projecting $\mathbf{y}$ onto the column space of the intercept (i.e., a vector of ones). The term $\mathbf{K}_g = \mathbf{X}_g \mathbf{A}_g \mathbf{A}_g \mathbf{X}_g^\intercal$ is commonly referred to as an $N \times N$ kernel matrix, where $\mathbf{A}_g = \text{diag}(a_1, \ldots, a_{|\mathcal{J}_g|})$ is used to denote a diagonal weight matrix that changes for each gene $g$. Each element of $\mathbf{K}_g$ is computed via the linear kernel function

$$k(\mathbf{x}_i, \mathbf{x}_{i'}) = \sum_{j=1}^{|\mathcal{J}_g|} a_j x_{ij} x_{i'j}. \tag{S12}$$

While implementing SKAT in this work, we follow previous works and set each weight to be $\sqrt{a_j} = \text{Beta}(\text{MAF}_j, 1, 25)$ — the beta distribution density function with pre-specified parameters evaluated at the sample minor allele frequency (MAF) for the $j$-th SNP in the gene region. For more details, see [11–14].

**Multi-marker Analysis of GenoMic Annotation (MAGMA).** In the current study, gene analyses with MAGMA also required access to raw phenotype $\mathbf{y}$ and genotype $\mathbf{X}$ information for $N$ individuals typed on $J$ SNPs. This approach is based on a multiple principal components regression model. In the first step, MAGMA projects the sub-genotype matrix for a gene $\mathbf{X}_g$ onto its principal components. Next, it prunes away PCs with very small eigenvalues, and then uses those reduced vectors as predictors for the phenotype in the linear regression model. Consider the following linear regression and singular-value decomposition (SVD) of the genotype matrix

$$\mathbf{y} = \beta_0 + \mathbf{X}\boldsymbol{\beta} + \mathbf{e}, \qquad \mathbf{X} = \mathbf{U}\boldsymbol{\Lambda}\mathbf{V}^{\intercal}, \qquad \mathbf{e} \sim \mathcal{N}(\mathbf{0}, \tau^2\mathbf{I}) \tag{S13}$$

where, in addition to the aforementioned notation, $\boldsymbol{\Lambda}$ is an $N \times J$ rectangular diagonal matrix of singular values, and $\mathbf{U}$ and $\mathbf{V}$ are $N \times N$ and $J \times J$ matrices of orthogonal unit vectors, respectively. For numerical stability and reduction of computational complexity, vectors corresponding to small eigenvalues can be truncated. Therefore, without loss of generality, MAGMA considers $\mathbf{V}$ and $\boldsymbol{\Lambda}$ to be of dimensions $J^* \times J^*$ and $N \times J^*$, respectively. Here, $J^*$ denotes the top eigenvalues explaining 99.9% of the cumulative variance in $\mathbf{X}_g$. By defining $\mathbf{G} = \mathbf{U}\boldsymbol{\Lambda}$, the model above simplifies to

$$\mathbf{y} = \beta_0 + \mathbf{G}\boldsymbol{\vartheta} + \mathbf{e}, \qquad \mathbf{e} \sim \mathcal{N}(\mathbf{0}, \tau^2\mathbf{I}) \tag{S14}$$

where $\boldsymbol{\vartheta} = \mathbf{V}^{\intercal}\boldsymbol{\beta}$ represents the lower-dimensional genetic effect. To derive a $P$-value for a single gene's association with the phenotype, MAGMA uses an F-test under the null hypothesis $H_0 : \boldsymbol{\vartheta} = \mathbf{0}$ or, equivalently, $H_0 : \mathbf{V}^{\intercal}\boldsymbol{\beta} = \mathbf{0}$. See [15] for more details.

### Supplementary Figures

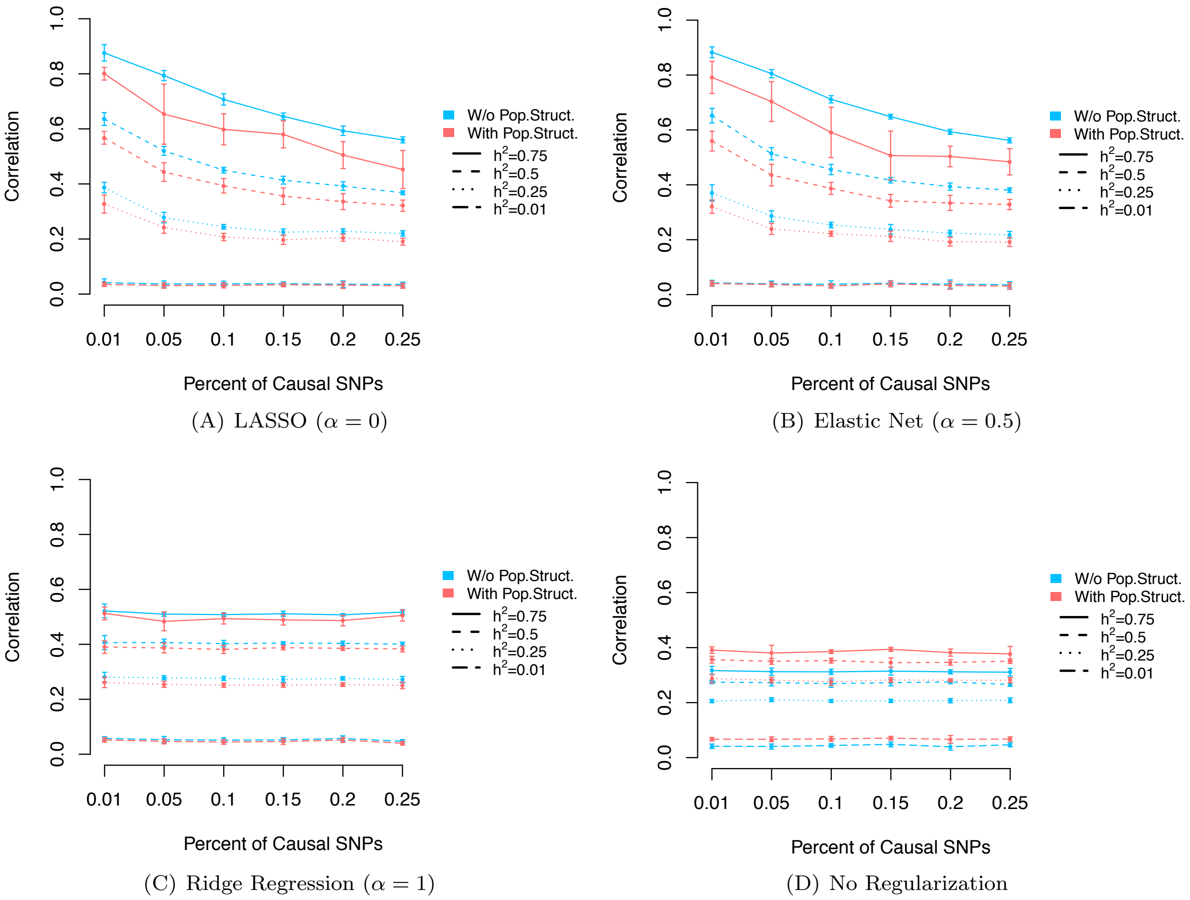

(A) LASSO ($\alpha = 0$)

(B) Elastic Net ($\alpha = 0.5$)

(C) Ridge Regression ($\alpha = 1$)

(D) No Regularization

**Figure S1. Simulation study results showing the Pearson correlation between various degrees of gene-$\varepsilon$ regularized SNP-level effect size estimates and the true effect sizes that generated the complex traits.** Assessed regularization techniques are the **(A)** LASSO [16], **(B)** Elastic Net [17], **(C)** Ridge Regression [18], and **(D)** no regularization of ordinary least squares (OLS) effect sizes which serves as a baseline. Here, we take real genotype data on chromosome 19 from $N = 5,000$ randomly chosen individuals of European ancestry in the UK Biobank (see Section S1). We then assumed a simple linear additive model for quantitative traits while varying the narrow-sense heritability ($h^2 = \{0.01, 0.05, 0.10, 0.15, 0.20, 0.25\}$). We considered two scenarios where traits are generated with and without additional population structure (colored as pink and blue lines, respectively). In the former setting, phenotypes are simulated while also using the top five principal components (PCs) of the genotype matrix as covariates to create stratification. These PCs contributed to 10% of the phenotypic variance. In both settings, GWA SNP-level effect sizes were derived via OLS without accounting for any additional structure. The y-axis shows Pearson correlation between gene-$\varepsilon$ regularized effect sizes and the truth. On the x-axis of each plot, we vary the number of causal SNPs for each trait (i.e., {1, 5, 10, 15, 20, 25}%). Results are based on ten replicates (see Section S2), with the error bars representing standard errors across runs.

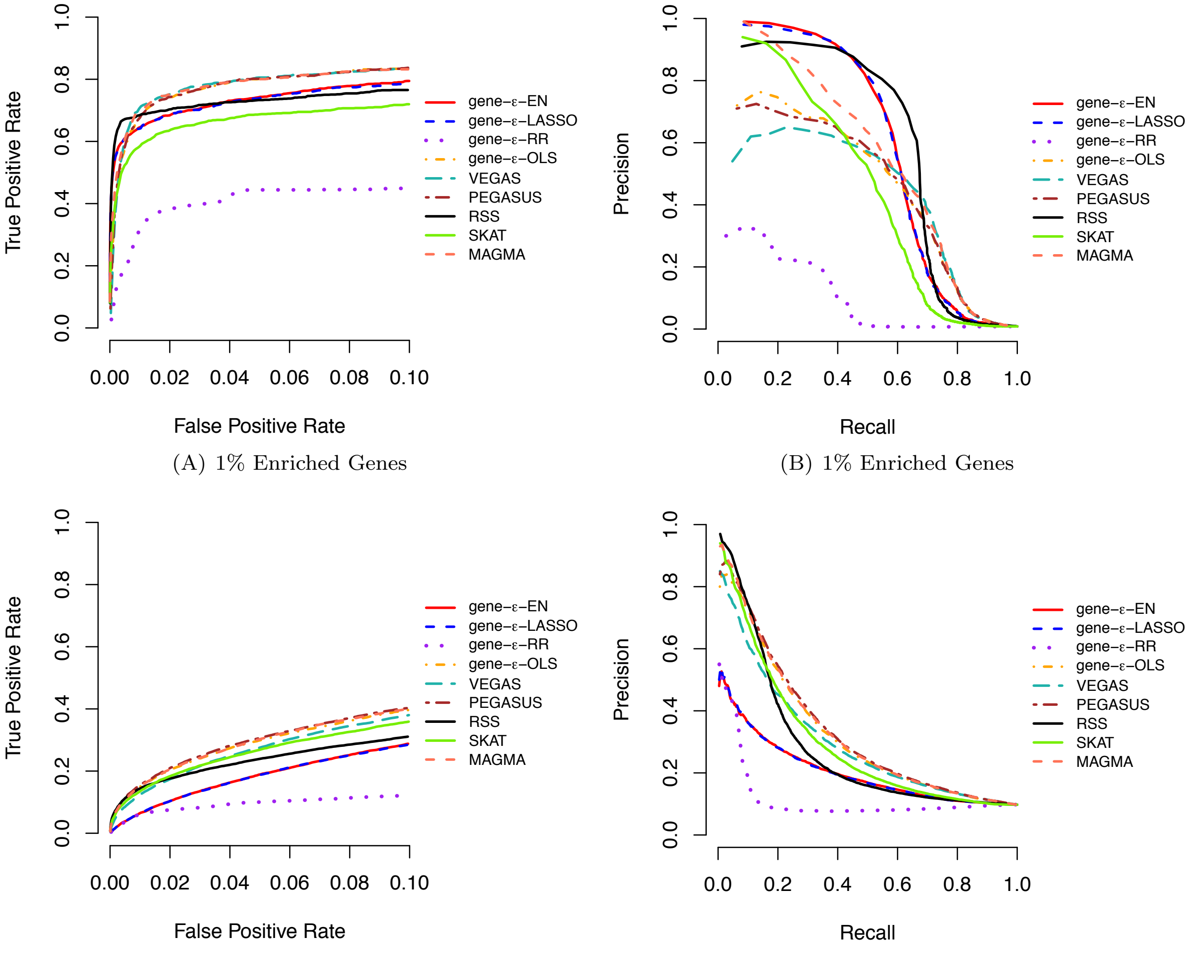

(A) 1% Enriched Genes (B) 1% Enriched Genes

(C) 10% Enriched Genes (D) 10% Enriched Genes

**Figure S2. (A, C) Receiver operating characteristic (ROC) and (B, D) precision-recall curves comparing the performance of gene-$\varepsilon$ and competing approaches in simulations ($N =$ 5,000; $h^2 = 0.2$).** Here, the sample size $N = 5,000$ and the narrow-sense heritability of the simulated quantitative trait is $h^2 = 0.2$. We compute standard GWA SNP-level effect sizes (estimated using ordinary least squares). Results for gene-$\varepsilon$ are shown with LASSO (blue), Elastic Net (EN; red), and Ridge Regression (RR; purple) regularizations. We also show the results of gene-$\varepsilon$ without regularization to illustrate the importance of the regularization step (labeled OLS; orange). We compare gene-$\varepsilon$ with five existing methods: PEGASUS (brown) [6], VEGAS (teal) [7], the Bayesian approach RSS (black) [10], SKAT (green) [11], and MAGMA (peach) [15]. **(A, C)** ROC curves show power versus false positive rate for each approach of sparse (1% enriched genes) and polygenic (10% enriched genes) architectures, respectively. Note that the upper limit of the x-axis has been truncated at 0.1. **(B, D)** Precision-Recall curves for each method applied to the simulations. Note that, in the sparse case (1% enriched genes), the top ranked genes are always true positives, and therefore the minimal recall is not 0. All results are based on 100 replicates (see Section S2).

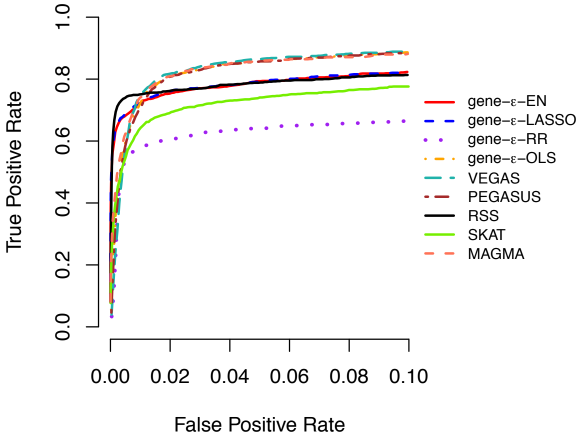

(A) 1% Enriched Genes

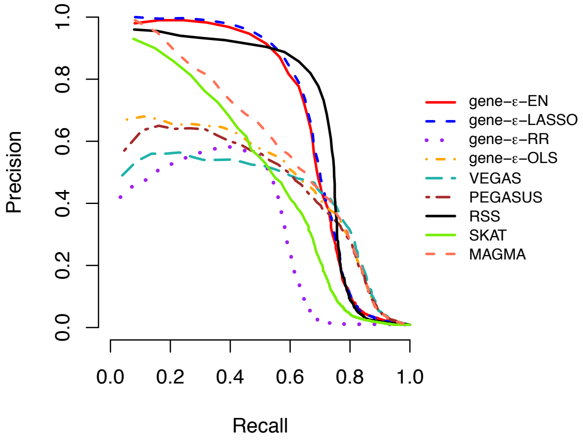

(B) 1% Enriched Genes

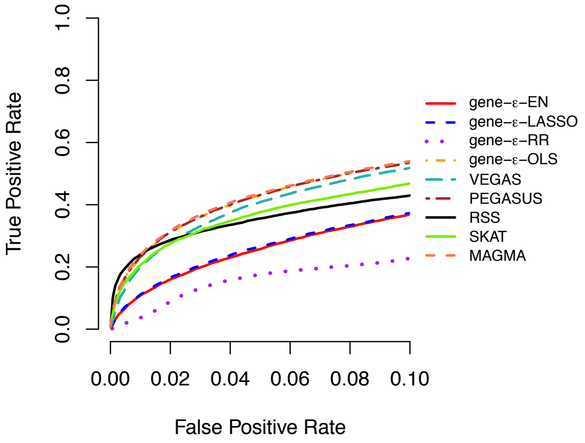

(C) 10% Enriched Genes

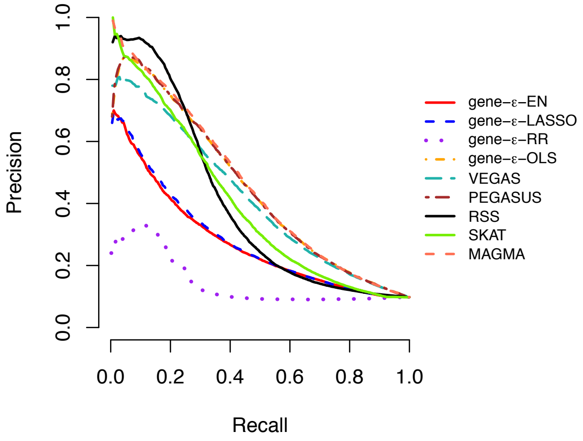

(D) 10% Enriched Genes

**Figure S3. (A, C) Receiver operating characteristic (ROC) and (B, D) precision-recall curves comparing the performance of gene-$\varepsilon$ and competing approaches in simulations ($N =$ 10,000; $h^2 = 0.2$).** Here, the sample size $N = 10,000$ and the narrow-sense heritability of the simulated quantitative trait is $h^2 = 0.2$. We compute standard GWA SNP-level effect sizes (estimated using ordinary least squares). Results for gene-$\varepsilon$ are shown with LASSO (blue), Elastic Net (EN; red), and Ridge Regression (RR; purple) regularizations. We also show the results of gene-$\varepsilon$ without regularization to illustrate the importance of the regularization step (labeled OLS; orange). We compare gene-$\varepsilon$ with five existing methods: PEGASUS (brown) [6], VEGAS (teal) [7], the Bayesian approach RSS (black) [10], SKAT (green) [11], and MAGMA (peach) [15]. **(A, C)** ROC curves show power versus false positive rate for each approach of sparse (1% enriched genes) and polygenic (10% enriched genes) architectures, respectively. Note that the upper limit of the x-axis has been truncated at 0.1. **(B, D)** Precision-Recall curves for each method applied to the simulations. Note that, in the sparse case (1% enriched genes), the top ranked genes are always true positives, and therefore the minimal recall is not 0. All results are based on 100 replicates (see Section S2).

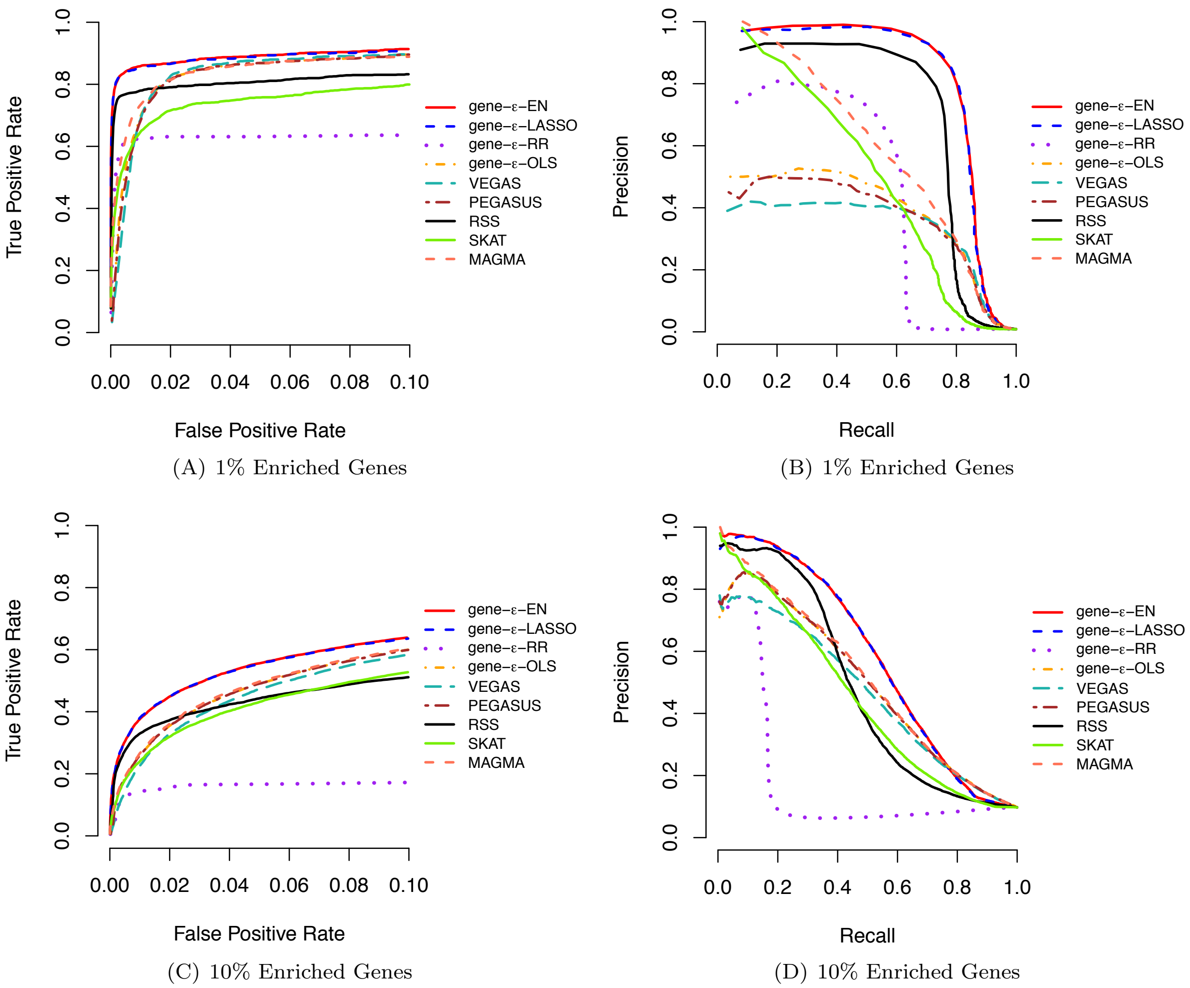

(A) 1% Enriched Genes

(B) 1% Enriched Genes

(C) 10% Enriched Genes

(D) 10% Enriched Genes

**Figure S4. (A, C) Receiver operating characteristic (ROC) and (B, D) precision-recall curves comparing the performance of gene-$\varepsilon$ and competing approaches in simulations ($N$ = 5,000; $h^2 = 0.6$).** Here, the sample size $N = 5,000$ and the narrow-sense heritability of the simulated quantitative trait is $h^2 = 0.6$. We compute standard GWA SNP-level effect sizes (estimated using ordinary least squares). Results for gene-$\varepsilon$ are shown with LASSO (blue), Elastic Net (EN; red), and Ridge Regression (RR; purple) regularizations. We also show the results of gene-$\varepsilon$ without regularization to illustrate the importance of the regularization step (labeled OLS; orange). We compare gene-$\varepsilon$ with five existing methods: PEGASUS (brown) [6], VEGAS (teal) [7], the Bayesian approach RSS (black) [10], SKAT (green) [11], and MAGMA (peach) [15]. **(A, C)** ROC curves show power versus false positive rate for each approach of sparse (1% enriched genes) and polygenic (10% enriched genes) architectures, respectively. Note that the upper limit of the x-axis has been truncated at 0.1. **(B, D)** Precision-Recall curves for each method applied to the simulations. Note that, in the sparse case (1% enriched genes), the top ranked genes are always true positives, and therefore the minimal recall is not 0. All results are based on 100 replicates (see Section S2).

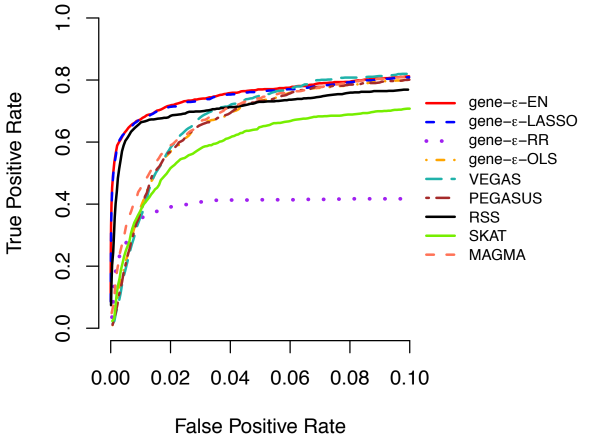

(A) 1% Enriched Genes

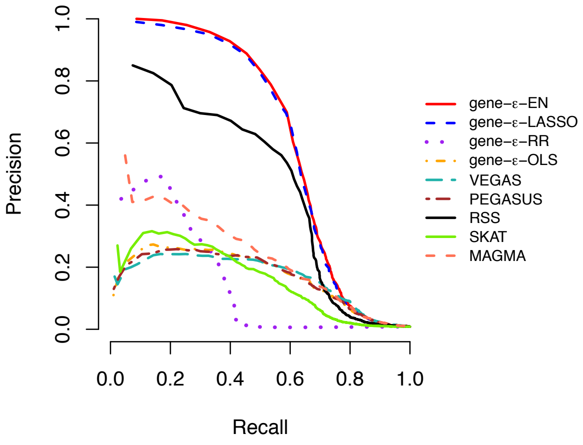

(B) 1% Enriched Genes

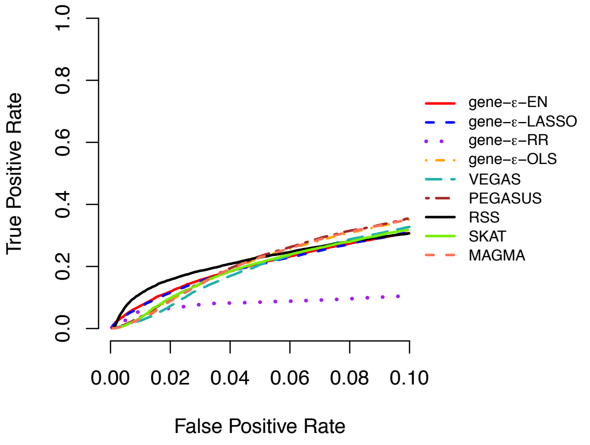

(C) 10% Enriched Genes

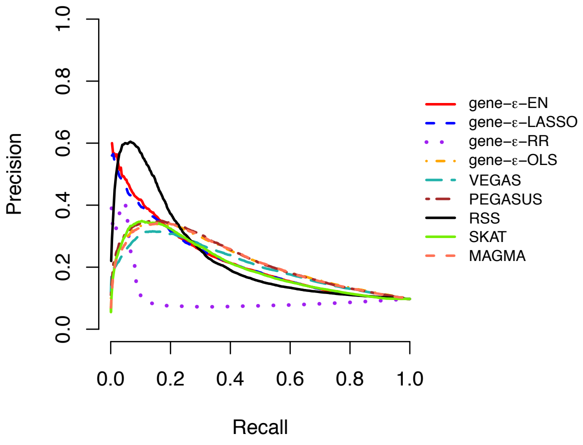

(D) 10% Enriched Genes

**Figure S5. (A, C) Receiver operating characteristic (ROC) and (B, D) precision-recall curves comparing the performance of gene-$\varepsilon$ and competing approaches in simulations with population stratification ($N$ = 5,000; $h^2$ = 0.2).** Here, the sample size $N = 5,000$ and the narrow-sense heritability of the simulated quantitative trait is $h^2 = 0.2$. In this simulation, traits were generated while using the top five principal components (PCs) of the genotype matrix as covariates. GWA summary statistics were computed by fitting a single-SNP univariate linear model (via ordinary least squares) without any control for the additional structure. Results for gene-$\varepsilon$ are shown with LASSO (blue), Elastic Net (EN; red), and Ridge Regression (RR; purple) regularizations. We also show the results of gene-$\varepsilon$ without regularization to illustrate the importance of the regularization step (labeled OLS; orange). We compare gene-$\varepsilon$ with five existing methods: PEGASUS (brown) [6], VEGAS (teal) [7], the Bayesian approach RSS (black) [10], SKAT (green) [11], and MAGMA (peach) [15]. Note that each was method implemented without using any covariates. **(A, C)** ROC curves show power versus false positive rate for each approach of sparse (1% enriched genes) and polygenic (10% enriched genes) architectures, respectively. Note that the upper limit of the x-axis has been truncated at 0.1. **(B, D)** Precision-Recall curves for each method applied to the simulations. Note that, in the sparse case (1% enriched genes), the top ranked genes are always true positives, and therefore the minimal recall is not 0. All results are based on 100 replicates (see Section S2).

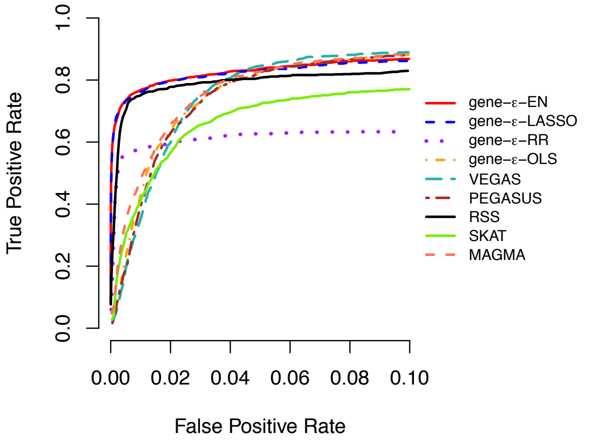

(A) 1% Enriched Genes

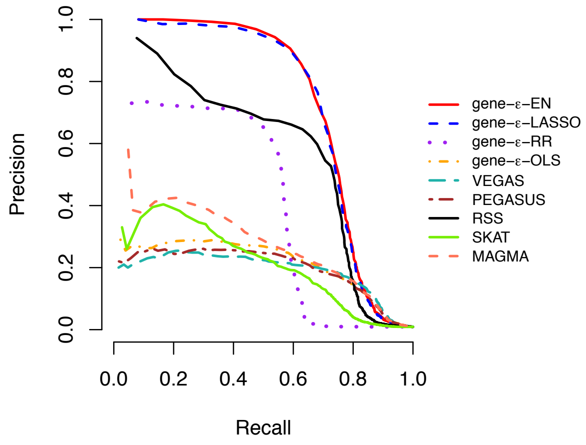

(B) 1% Enriched Genes

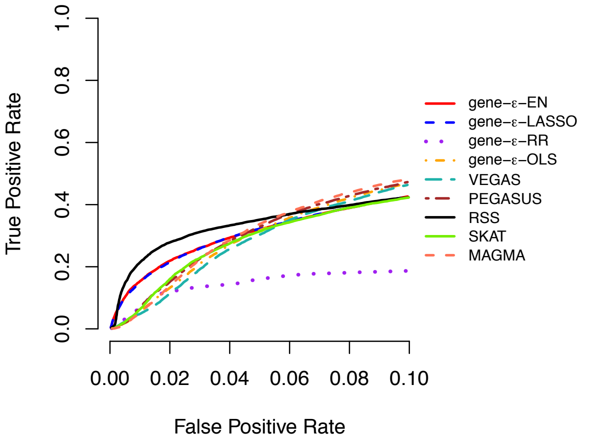

(C) 10% Enriched Genes

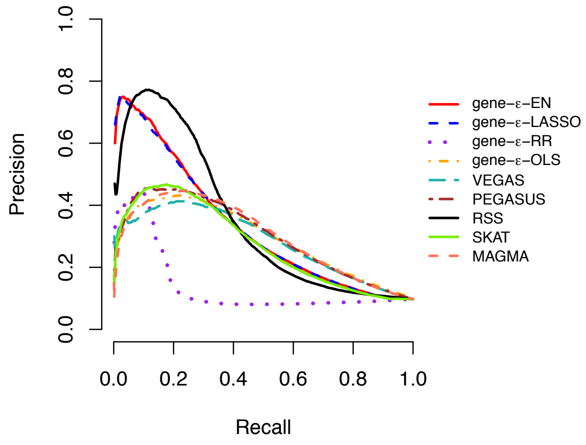

(D) 10% Enriched Genes

**Figure S6. (A, C) Receiver operating characteristic (ROC) and (B, D) precision-recall curves comparing the performance of gene-$\varepsilon$ and competing approaches in simulations with population stratification ($N$ = 10,000; $h^2$ = 0.2).** Here, the sample size $N = 10,000$ and the narrow-sense heritability of the simulated quantitative trait is $h^2 = 0.2$. In this simulation, traits were generated while using the top five principal components (PCs) of the genotype matrix as covariates. GWA summary statistics were computed by fitting a single-SNP univariate linear model (via ordinary least squares) without any control for the additional structure. Results for gene-$\varepsilon$ are shown with LASSO (blue), Elastic Net (EN; red), and Ridge Regression (RR; purple) regularizations. We also show the results of gene-$\varepsilon$ without regularization to illustrate the importance of the regularization step (labeled OLS; orange). We compare gene-$\varepsilon$ with five existing methods: PEGASUS (brown) [6], VEGAS (teal) [7], the Bayesian approach RSS (black) [10], SKAT (green) [11], and MAGMA (peach) [15]. Note that each was method implemented without using any covariates. **(A, C)** ROC curves show power versus false positive rate for each approach of sparse (1% enriched genes) and polygenic (10% enriched genes) architectures, respectively. Note that the upper limit of the x-axis has been truncated at 0.1. **(B, D)** Precision-Recall curves for each method applied to the simulations. Note that, in the sparse case (1% enriched genes), the top ranked genes are always true positives, and therefore the minimal recall is not 0. All results are based on 100 replicates (see Section S2).

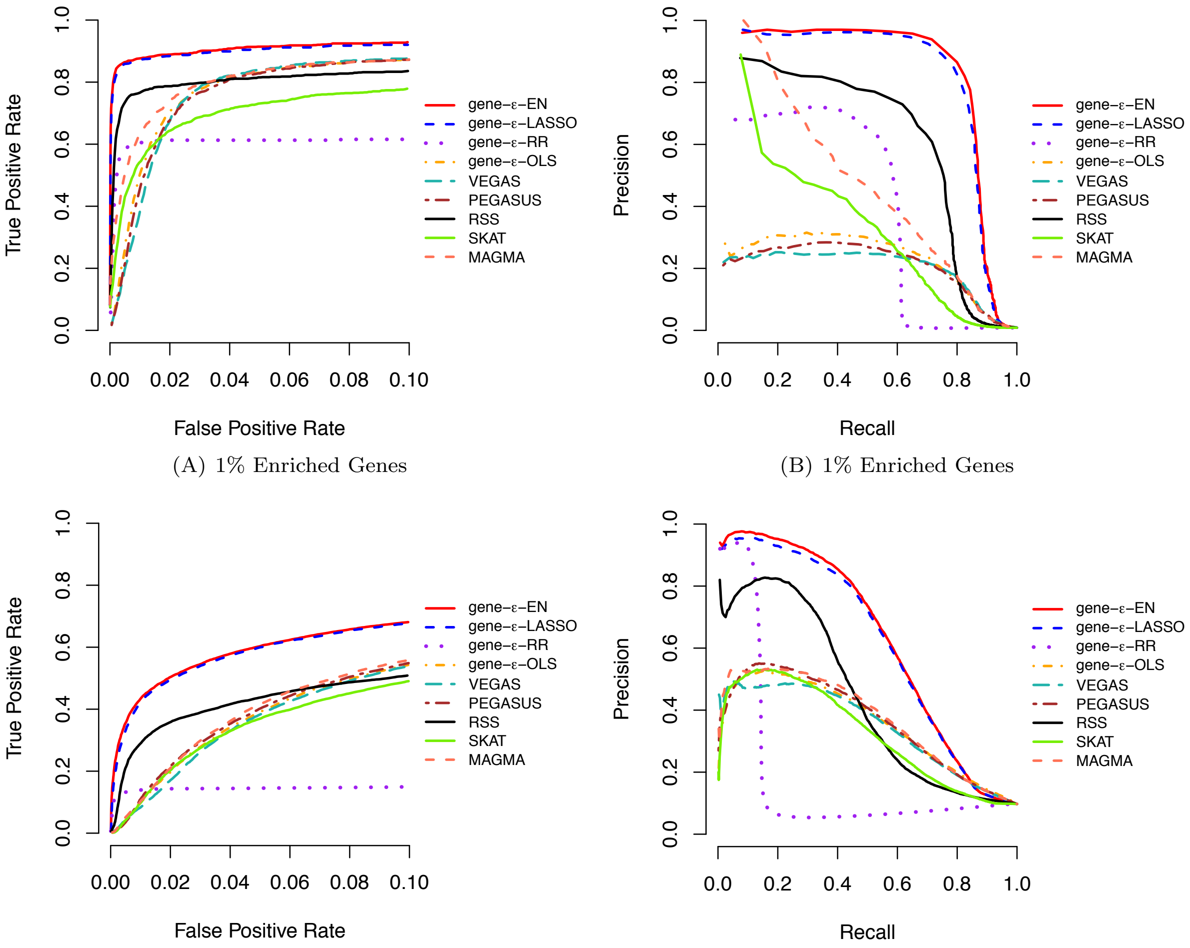

(A) 1% Enriched Genes

(B) 1% Enriched Genes

(C) 10% Enriched Genes

(D) 10% Enriched Genes

**Figure S7. (A, C) Receiver operating characteristic (ROC) and (B, D) precision-recall curves comparing the performance of gene-$\varepsilon$ and competing approaches in simulations with population stratification ($N$ = 5,000; $h^2$ = 0.6).** Here, the sample size $N = 5,000$ and the narrow-sense heritability of the simulated quantitative trait is $h^2 = 0.6$. In this simulation, traits were generated while using the top five principal components (PCs) of the genotype matrix as covariates. GWA summary statistics were computed by fitting a single-SNP univariate linear model (via ordinary least squares) without any control for the additional structure. Results for gene-$\varepsilon$ are shown with LASSO (blue), Elastic Net (EN; red), and Ridge Regression (RR; purple) regularizations. We also show the results of gene-$\varepsilon$ without regularization to illustrate the importance of the regularization step (labeled OLS; orange). We compare gene-$\varepsilon$ with five existing methods: PEGASUS (brown) [6], VEGAS (teal) [7], the Bayesian approach RSS (black) [10], SKAT (green) [11], and MAGMA (peach) [15]. Note that each was method implemented without using any covariates. **(A, C)** ROC curves show power versus false positive rate for each approach of sparse (1% enriched genes) and polygenic (10% enriched genes) architectures, respectively. Note that the upper limit of the x-axis has been truncated at 0.1. **(B, D)** Precision-Recall curves for each method applied to the simulations. Note that, in the sparse case (1% enriched genes), the top ranked genes are always true positives, and therefore the minimal recall is not 0. All results are based on 100 replicates (see Section S2).

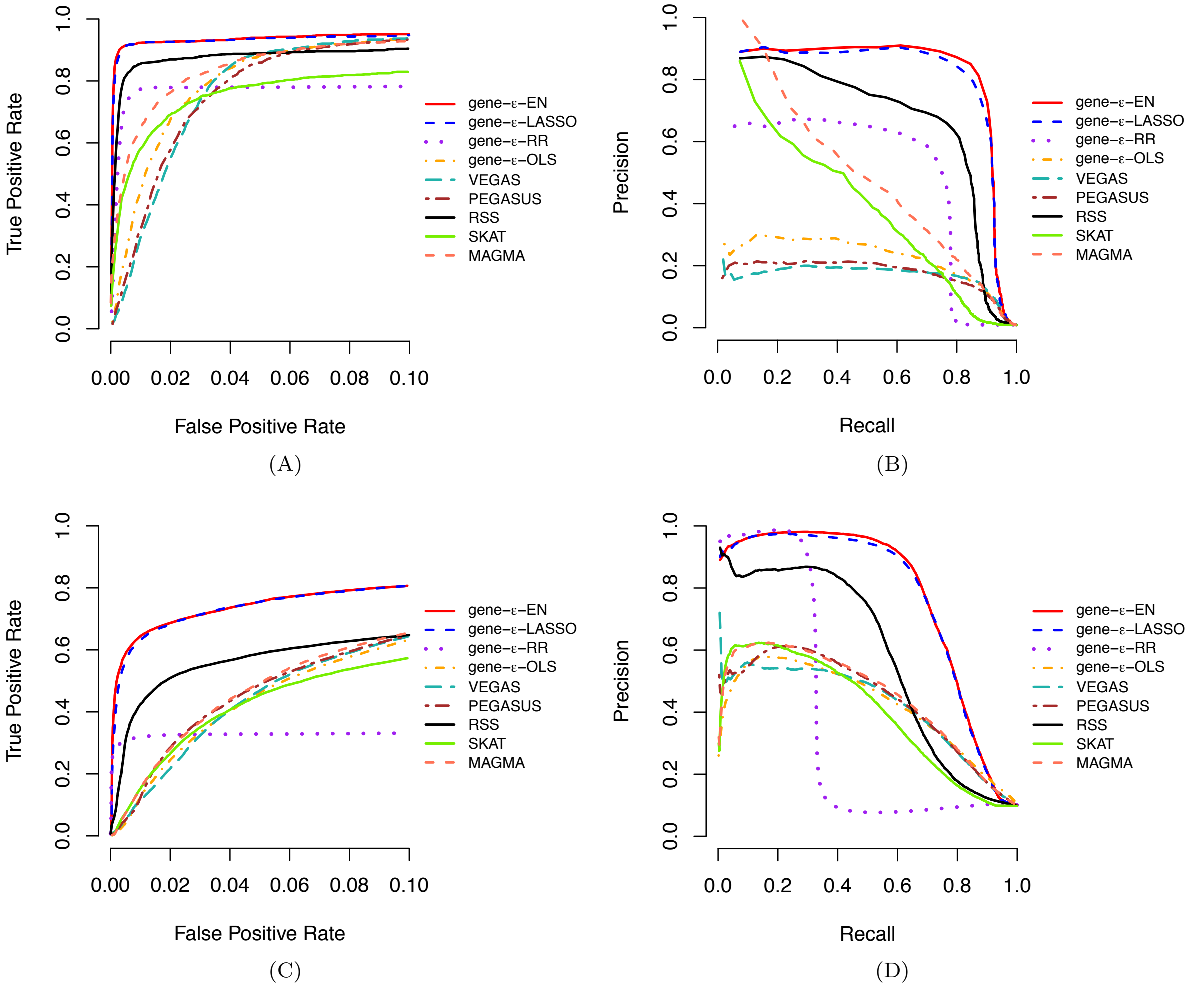

**Figure S8. (A, C) Receiver operating characteristic (ROC) and (B, D) precision-recall curves comparing the performance of gene-$\varepsilon$ and competing approaches in simulations with population stratification ($N$ = 10,000; $h^2$ = 0.6).** Here, the sample size $N = 10,000$ and the narrow-sense heritability of the simulated quantitative trait is $h^2 = 0.6$. In this simulation, traits were generated while using the top five principal components (PCs) of the genotype matrix as covariates. GWA summary statistics were computed by fitting a single-SNP univariate linear model (via ordinary least squares) without any control for the additional structure. Results for gene-$\varepsilon$ are shown with LASSO (blue), Elastic Net (EN; red), and Ridge Regression (RR; purple) regularizations. We also show the results of gene-$\varepsilon$ without regularization to illustrate the importance of the regularization step (labeled OLS; orange). We compare gene-$\varepsilon$ with five existing methods: PEGASUS (brown) [6], VEGAS (teal) [7], the Bayesian approach RSS (black) [10], SKAT (green) [11], and MAGMA (peach) [15]. Note that each was method implemented without using any covariates. **(A, C)** ROC curves show power versus false positive rate for each approach of sparse (1% enriched genes) and polygenic (10% enriched genes) architectures, respectively. Note that the upper limit of the x-axis has been truncated at 0.1. **(B, D)** Precision-Recall curves for each method applied to the simulations. Note that, in the sparse case (1% enriched genes), the top ranked genes are always true positives, and therefore the minimal recall is not 0. All results are based on 100 replicates (see Section S2).

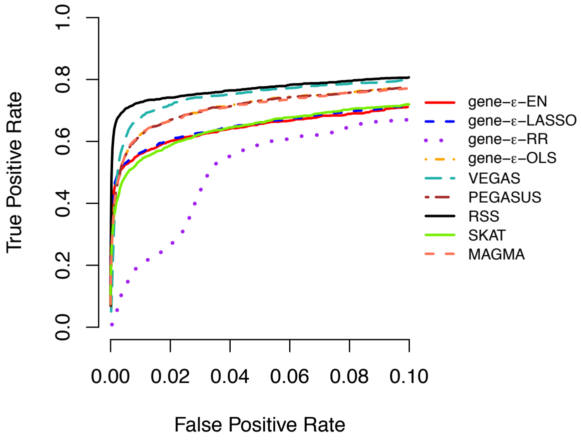

(A) 1% Enriched Genes

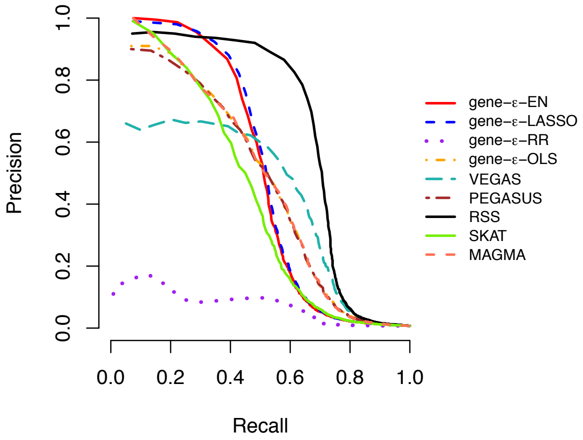

(B) 1% Enriched Genes

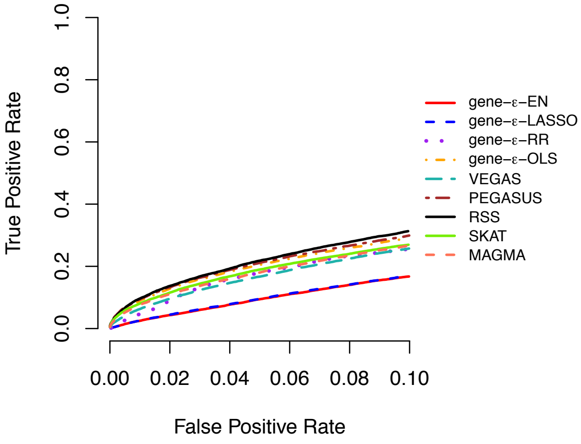

(C) 10% Enriched Genes

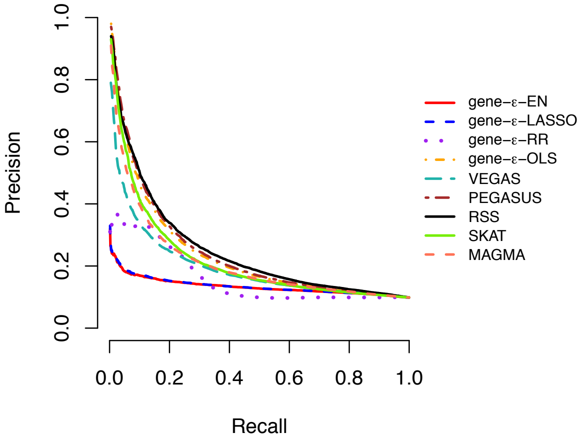

(D) 10% Enriched Genes

**Figure S9. (A, C) Receiver operating characteristic (ROC) and (B, D) precision-recall curves comparing the performance of gene-$\varepsilon$ and competing approaches in simulations with gene boundaries augmented by a 50 kilobase (kb) buffer ($N$ = 5,000; $h^2$ = 0.2).** Here, the sample size $N = 5,000$ and the narrow-sense heritability of the simulated quantitative trait is $h^2 = 0.2$. We compute standard GWA SNP-level effect sizes (estimated using ordinary least squares). Results for gene-$\varepsilon$ are shown with LASSO (blue), Elastic Net (EN; red), and Ridge Regression (RR; purple) regularizations. We also show the results of gene-$\varepsilon$ without regularization to illustrate the importance of the regularization step (labeled OLS; orange). We compare gene-$\varepsilon$ with five existing methods: PEGASUS (brown) [6], VEGAS (teal) [7], the Bayesian approach RSS (black) [10], SKAT (green) [11], and MAGMA (peach) [15]. **(A, C)** ROC curves show power versus false positive rate for each approach of sparse (1% enriched genes) and polygenic (10% enriched genes) architectures, respectively. Note that the upper limit of the x-axis has been truncated at 0.1. **(B, D)** Precision-Recall curves for each method applied to the simulations. Note that, in the sparse case (1% enriched genes), the top ranked genes are always true positives, and therefore the minimal recall is not 0. All results are based on 100 replicates (see Section S2).

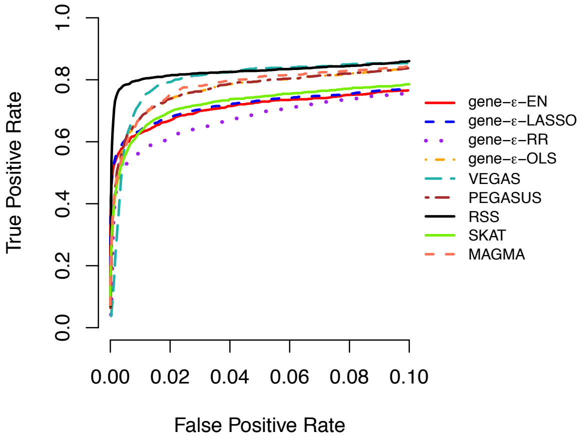

(A) 1% Enriched Genes

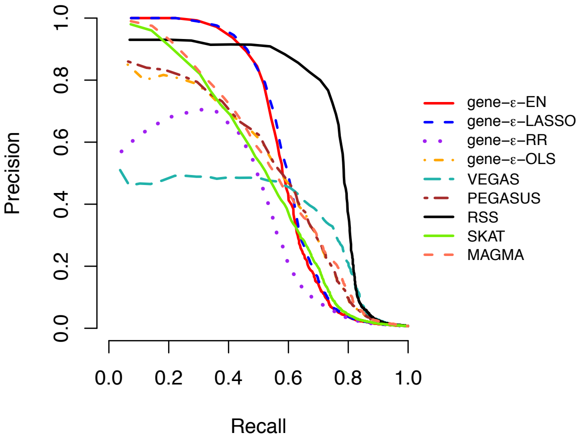

(B) 1% Enriched Genes

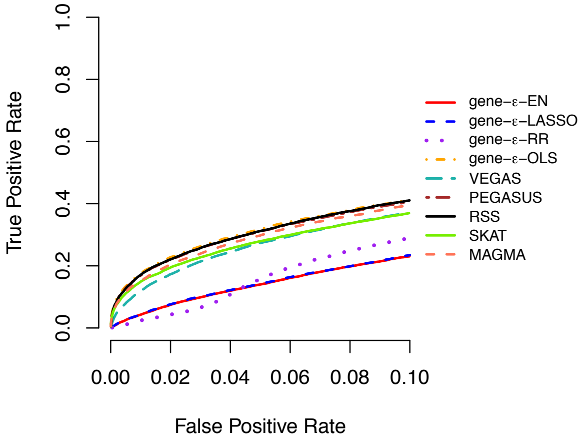

(C) 10% Enriched Genes

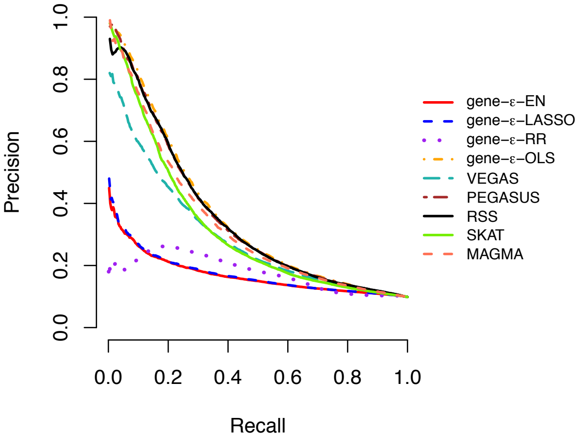

(D) 10% Enriched Genes

**Figure S10. (A, C) Receiver operating characteristic (ROC) and (B, D) precision-recall curves comparing the performance of gene-$\varepsilon$ and competing approaches in simulations with gene boundaries augmented by a 50 kilobase (kb) buffer ($N$ = 10,000; $h^2 = 0.2$).** Here, the sample size $N = 10,000$ and the narrow-sense heritability of the simulated quantitative trait is $h^2 = 0.2$. We compute standard GWA SNP-level effect sizes (estimated using ordinary least squares). Results for gene-$\varepsilon$ are shown with LASSO (blue), Elastic Net (EN; red), and Ridge Regression (RR; purple) regularizations. We also show the results of gene-$\varepsilon$ without regularization to illustrate the importance of the regularization step (labeled OLS; orange). We compare gene-$\varepsilon$ with five existing methods: PEGASUS (brown) [6], VEGAS (teal) [7], the Bayesian approach RSS (black) [10], SKAT (green) [11], and MAGMA (peach) [15]. **(A, C)** ROC curves show power versus false positive rate for each approach of sparse (1% enriched genes) and polygenic (10% enriched genes) architectures, respectively. Note that the upper limit of the x-axis has been truncated at 0.1. **(B, D)** Precision-Recall curves for each method applied to the simulations. Note that, in the sparse case (1% enriched genes), the top ranked genes are always true positives, and therefore the minimal recall is not 0. All results are based on 100 replicates (see Section S2).

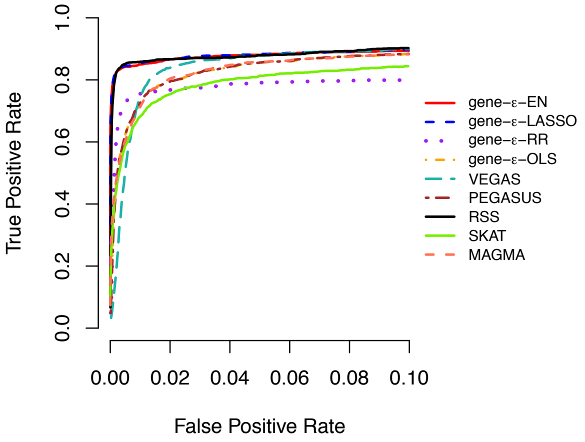

(A) 1% Enriched Genes

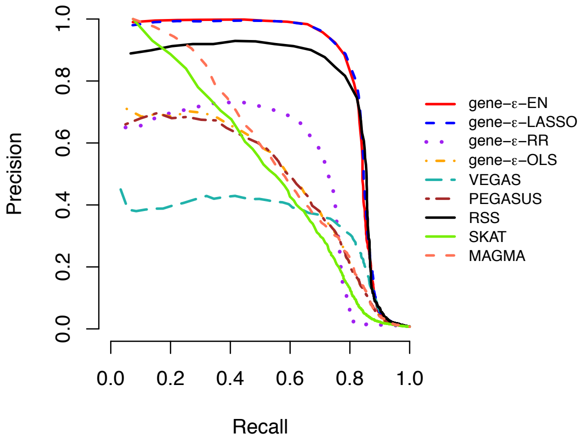

(B) 1% Enriched Genes

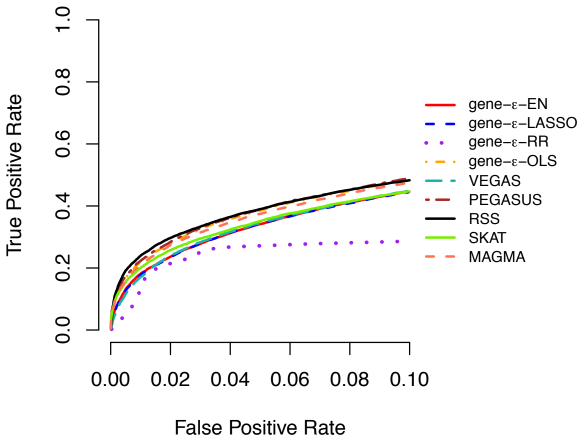

(C) 10% Enriched Genes

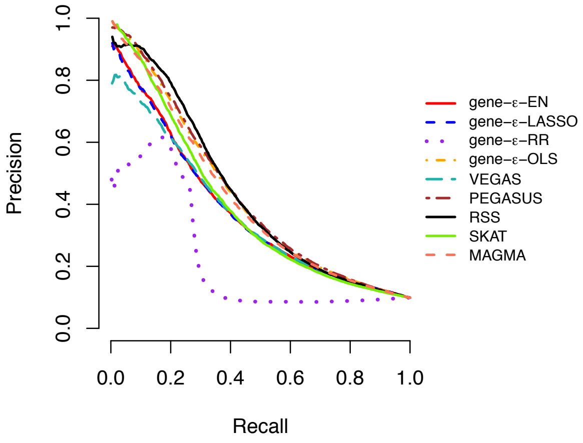

(D) 10% Enriched Genes

**Figure S11. (A, C) Receiver operating characteristic (ROC) and (B, D) precision-recall curves comparing the performance of gene-$\varepsilon$ and competing approaches in simulations with gene boundaries augmented by a 50 kilobase (kb) buffer ($N$ = 5,000; $h^2$ = 0.6).** Here, the sample size $N = 5,000$ and the narrow-sense heritability of the simulated quantitative trait is $h^2 = 0.6$. We compute standard GWA SNP-level effect sizes (estimated using ordinary least squares). Results for gene-$\varepsilon$ are shown with LASSO (blue), Elastic Net (EN; red), and Ridge Regression (RR; purple) regularizations. We also show the results of gene-$\varepsilon$ without regularization to illustrate the importance of the regularization step (labeled OLS; orange). We compare gene-$\varepsilon$ with five existing methods: PEGASUS (brown) [6], VEGAS (teal) [7], the Bayesian approach RSS (black) [10], SKAT (green) [11], and MAGMA (peach) [15]. **(A, C)** ROC curves show power versus false positive rate for each approach of sparse (1% enriched genes) and polygenic (10% enriched genes) architectures, respectively. Note that the upper limit of the x-axis has been truncated at 0.1. **(B, D)** Precision-Recall curves for each method applied to the simulations. Note that, in the sparse case (1% enriched genes), the top ranked genes are always true positives, and therefore the minimal recall is not 0. All results are based on 100 replicates (see Section S2).

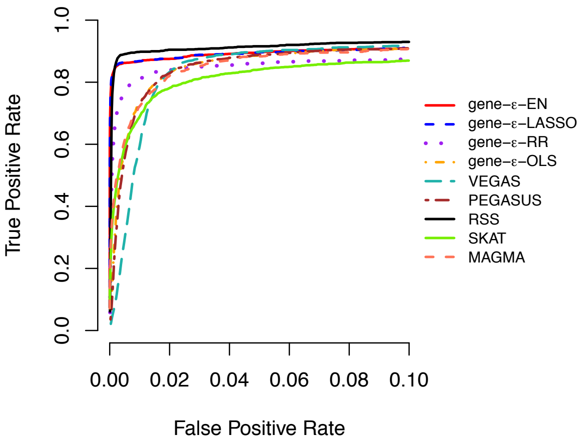

(A) 1% Enriched Genes

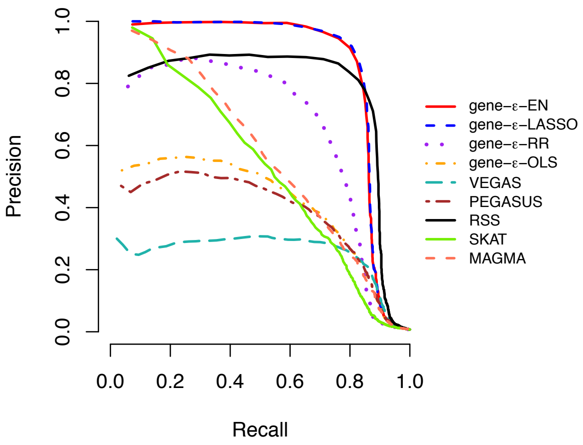

(B) 1% Enriched Genes

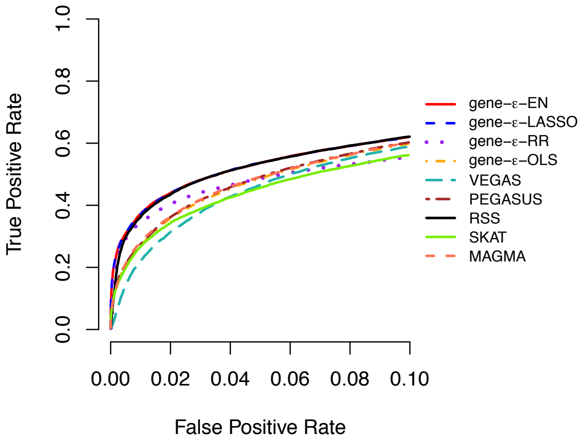

(C) 10% Enriched Genes

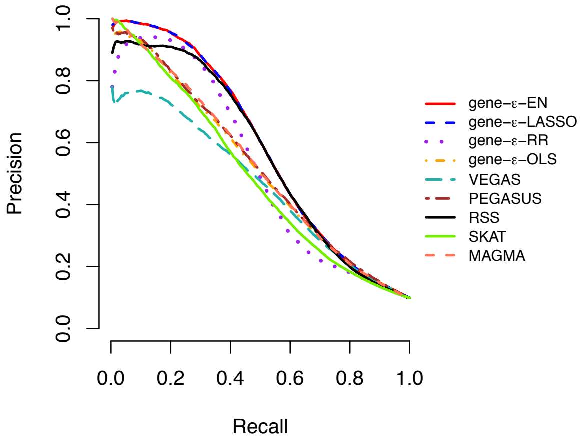

(D) 10% Enriched Genes

**Figure S12. (A, C) Receiver operating characteristic (ROC) and (B, D) precision-recall curves comparing the performance of gene-$\varepsilon$ and competing approaches in simulations with gene boundaries augmented by a 50 kilobase (kb) buffer ($N$ = 10,000; $h^2$ = 0.6).** Here, the sample size $N = 10,000$ and the narrow-sense heritability of the simulated quantitative trait is $h^2 = 0.6$. We compute standard GWA SNP-level effect sizes (estimated using ordinary least squares). Results for gene-$\varepsilon$ are shown with LASSO (blue), Elastic Net (EN; red), and Ridge Regression (RR; purple) regularizations. We also show the results of gene-$\varepsilon$ without regularization to illustrate the importance of the regularization step (labeled OLS; orange). We compare gene-$\varepsilon$ with five existing methods: PEGASUS (brown) [6], VEGAS (teal) [7], the Bayesian approach RSS (black) [10], SKAT (green) [11], and MAGMA (peach) [15]. **(A, C)** ROC curves show power versus false positive rate for each approach of sparse (1% enriched genes) and polygenic (10% enriched genes) architectures, respectively. Note that the upper limit of the x-axis has been truncated at 0.1. **(B, D)** Precision-Recall curves for each method applied to the simulations. Note that, in the sparse case (1% enriched genes), the top ranked genes are always true positives, and therefore the minimal recall is not 0. All results are based on 100 replicates (see Section S2).

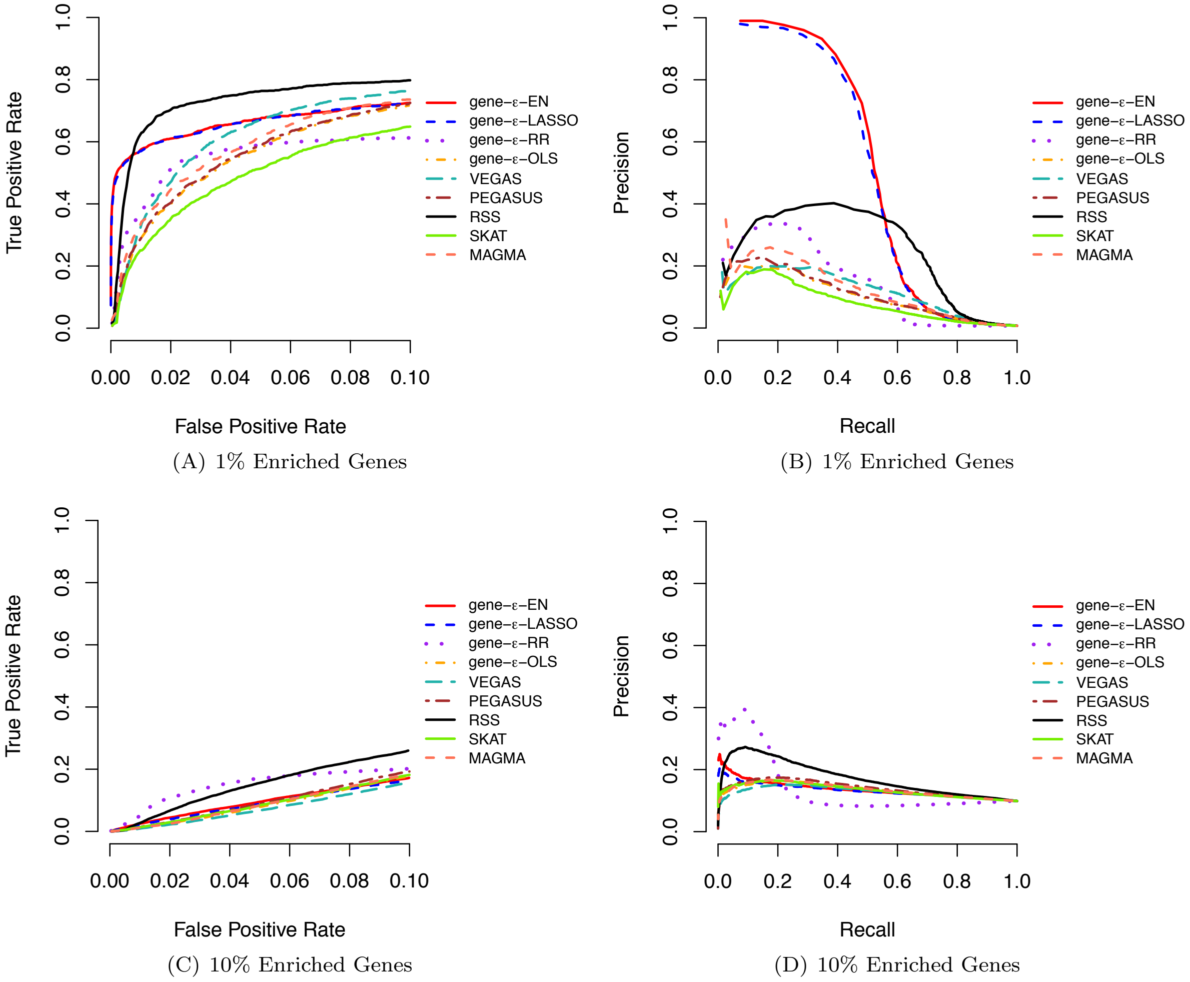

**Figure S13. (A, C) Receiver operating characteristic (ROC) and (B, D) precision-recall curves comparing the performance of gene-$\varepsilon$ and competing approaches in simulations with gene boundaries augmented by a 50 kilobase (kb) buffer and with population stratification ($N$ = 5,000; $h^2$ = 0.2).** Here, the sample size $N = 5,000$ and the narrow-sense heritability of the simulated quantitative trait is $h^2 = 0.2$. In this simulation, traits were generated while using the top five principal components (PCs) of the genotype matrix as covariates. GWA summary statistics were computed by fitting a single-SNP univariate linear model (via ordinary least squares) without any control for the additional structure. Results for gene-$\varepsilon$ are shown with LASSO (blue), Elastic Net (EN; red), and Ridge Regression (RR; purple) regularizations. We also show the results of gene-$\varepsilon$ without regularization to illustrate the importance of the regularization step (labeled OLS; orange). We compare gene-$\varepsilon$ with five existing methods: PEGASUS (brown) [6], VEGAS (teal) [7], the Bayesian approach RSS (black) [10], SKAT (green) [11], and MAGMA (peach) [15]. Note that each was method implemented without using any covariates. **(A, C)** ROC curves show power versus false positive rate for each approach of sparse (1% enriched genes) and polygenic (10% enriched genes) architectures, respectively. Note that the upper limit of the x-axis has been truncated at 0.1. **(B, D)** Precision-Recall curves for each method applied to the simulations. Note that, in the sparse case (1% enriched genes), the top ranked genes are always true positives, and therefore the minimal recall is not 0. All results are based on 100 replicates (see Section S2).

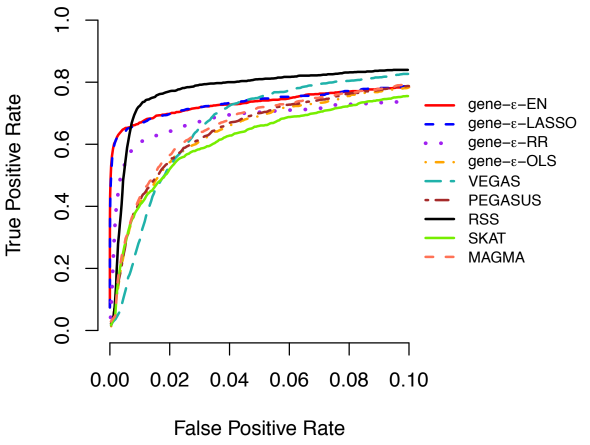

(A) 1% Enriched Genes

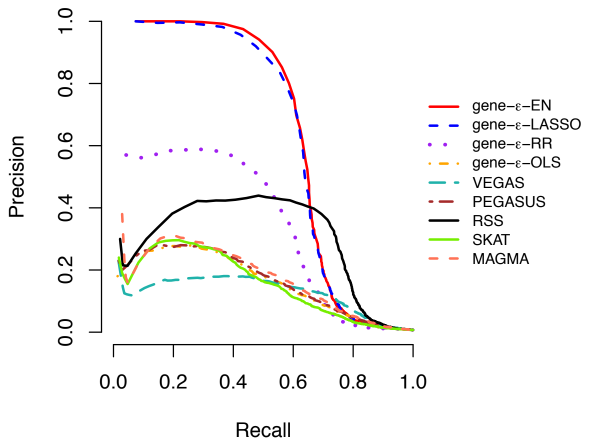

(B) 1% Enriched Genes

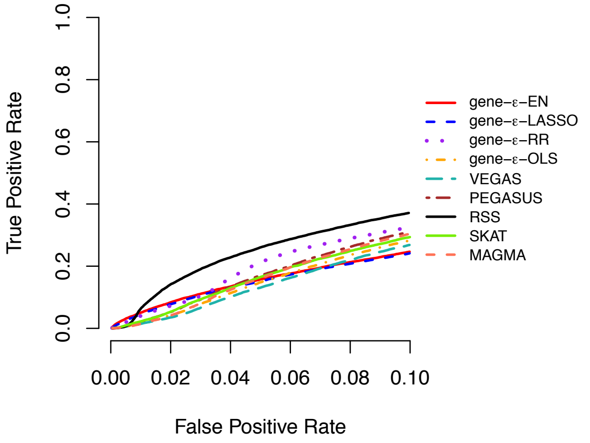

(C) 10% Enriched Genes

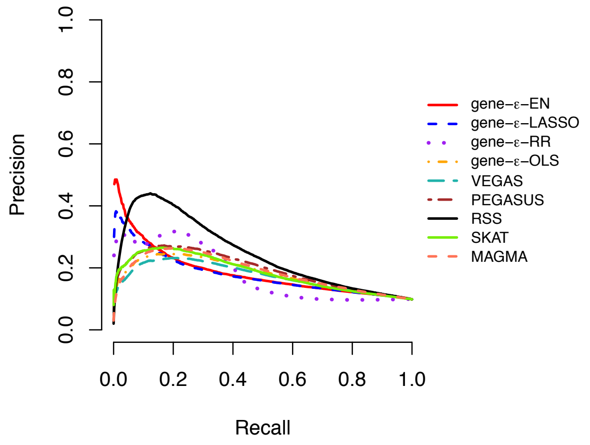

(D) 10% Enriched Genes

**Figure S14. (A, C) Receiver operating characteristic (ROC) and (B, D) precision-recall curves comparing the performance of gene-$\varepsilon$ and competing approaches in simulations with gene boundaries augmented by a 50 kilobase (kb) buffer and with population stratification ($N$ = 10,000; $h^2$ = 0.2).** Here, the sample size $N = 10,000$ and the narrow-sense heritability of the simulated quantitative trait is $h^2 = 0.2$. In this simulation, traits were generated while using the top five principal components (PCs) of the genotype matrix as covariates. GWA summary statistics were computed by fitting a single-SNP univariate linear model (via ordinary least squares) without any control for the additional structure. Results for gene-$\varepsilon$ are shown with LASSO (blue), Elastic Net (EN; red), and Ridge Regression (RR; purple) regularizations. We also show the results of gene-$\varepsilon$ without regularization to illustrate the importance of the regularization step (labeled OLS; orange). We compare gene-$\varepsilon$ with five existing methods: PEGASUS (brown) [6], VEGAS (teal) [7], the Bayesian approach RSS (black) [10], SKAT (green) [11], and MAGMA (peach) [15]. Note that each was method implemented without using any covariates. **(A, C)** ROC curves show power versus false positive rate for each approach of sparse (1% enriched genes) and polygenic (10% enriched genes) architectures, respectively. Note that the upper limit of the x-axis has been truncated at 0.1. **(B, D)** Precision-Recall curves for each method applied to the simulations. Note that, in the sparse case (1% enriched genes), the top ranked genes are always true positives, and therefore the minimal recall is not 0. All results are based on 100 replicates (see Section S2).

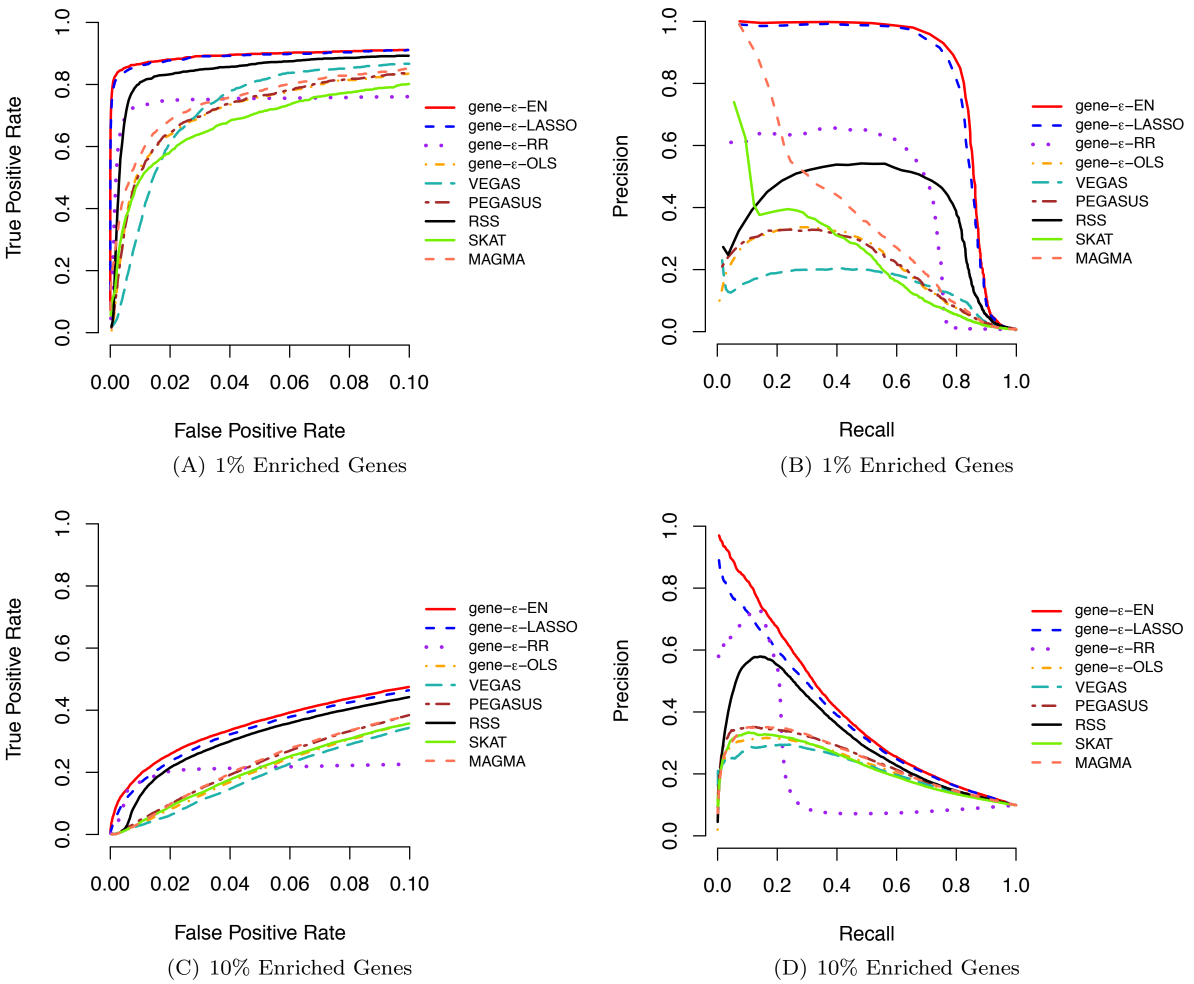

(A) 1% Enriched Genes (B) 1% Enriched Genes

(C) 10% Enriched Genes (D) 10% Enriched Genes

**Figure S15. (A, C) Receiver operating characteristic (ROC) and (B, D) precision-recall curves comparing the performance of gene-$\varepsilon$ and competing approaches in simulations with gene boundaries augmented by a 50 kilobase (kb) buffer and with population stratification ($N$ = 5,000; $h^2$ = 0.6).** Here, the sample size $N = 5,000$ and the narrow-sense heritability of the simulated quantitative trait is $h^2 = 0.6$. In this simulation, traits were generated while using the top five principal components (PCs) of the genotype matrix as covariates. GWA summary statistics were computed by fitting a single-SNP univariate linear model (via ordinary least squares) without any control for the additional structure. Results for gene-$\varepsilon$ are shown with LASSO (blue), Elastic Net (EN; red), and Ridge Regression (RR; purple) regularizations. We also show the results of gene-$\varepsilon$ without regularization to illustrate the importance of the regularization step (labeled OLS; orange). We compare gene-$\varepsilon$ with five existing methods: PEGASUS (brown) [6], VEGAS (teal) [7], the Bayesian approach RSS (black) [10], SKAT (green) [11], and MAGMA (peach) [15]. Note that each was method implemented without using any covariates. **(A, C)** ROC curves show power versus false positive rate for each approach of sparse (1% enriched genes) and polygenic (10% enriched genes) architectures, respectively. Note that the upper limit of the x-axis has been truncated at 0.1. **(B, D)** Precision-Recall curves for each method applied to the simulations. Note that, in the sparse case (1% enriched genes), the top ranked genes are always true positives, and therefore the minimal recall is not 0. All results are based on 100 replicates (see Section S2).

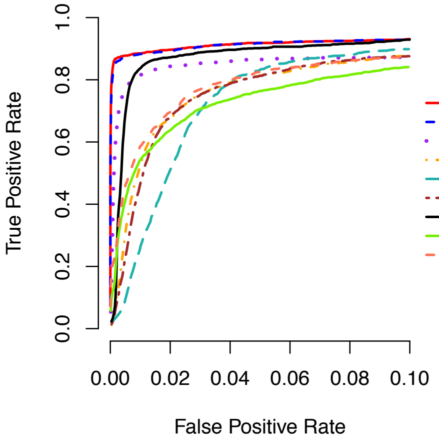

(A) 1% Enriched Genes

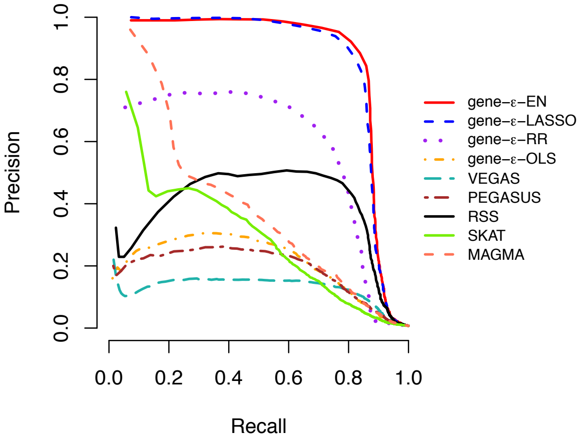

(B) 1% Enriched Genes

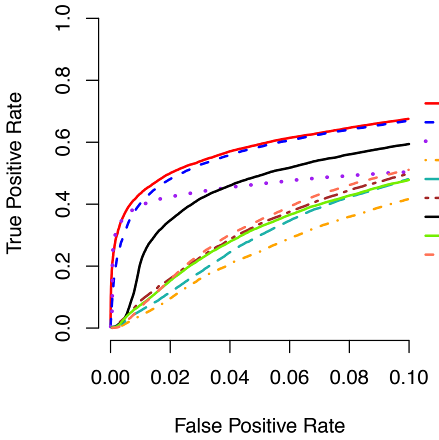

(C) 10% Enriched Genes

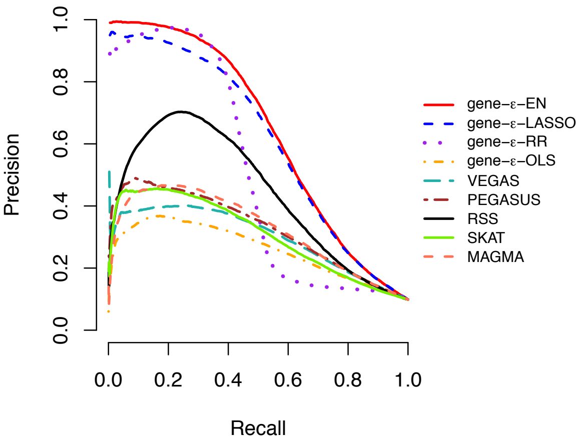

(D) 10% Enriched Genes

**Figure S16. (A, C) Receiver operating characteristic (ROC) and (B, D) precision-recall curves comparing the performance of gene-$\varepsilon$ and competing approaches in simulations with gene boundaries augmented by a 50 kilobase (kb) buffer and with population stratification ($N$ = 10,000; $h^2$ = 0.6).** Here, the sample size $N = 10,000$ and the narrow-sense heritability of the simulated quantitative trait is $h^2 = 0.6$. In this simulation, traits were generated while using the top five principal components (PCs) of the genotype matrix as covariates. GWA summary statistics were computed by fitting a single-SNP univariate linear model (via ordinary least squares) without any control for the additional structure. Results for gene-$\varepsilon$ are shown with LASSO (blue), Elastic Net (EN; red), and Ridge Regression (RR; purple) regularizations. We also show the results of gene-$\varepsilon$ without regularization to illustrate the importance of the regularization step (labeled OLS; orange). We compare gene-$\varepsilon$ with five existing methods: PEGASUS (brown) [6], VEGAS (teal) [7], the Bayesian approach RSS (black) [10], SKAT (green) [11], and MAGMA (peach) [15]. Note that each was method implemented without using any covariates. **(A, C)** ROC curves show power versus false positive rate for each approach of sparse (1% enriched genes) and polygenic (10% enriched genes) architectures, respectively. Note that the upper limit of the x-axis has been truncated at 0.1. **(B, D)** Precision-Recall curves for each method applied to the simulations. Note that, in the sparse case (1% enriched genes), the top ranked genes are always true positives, and therefore the minimal recall is not 0. All results are based on 100 replicates (see Section S2).

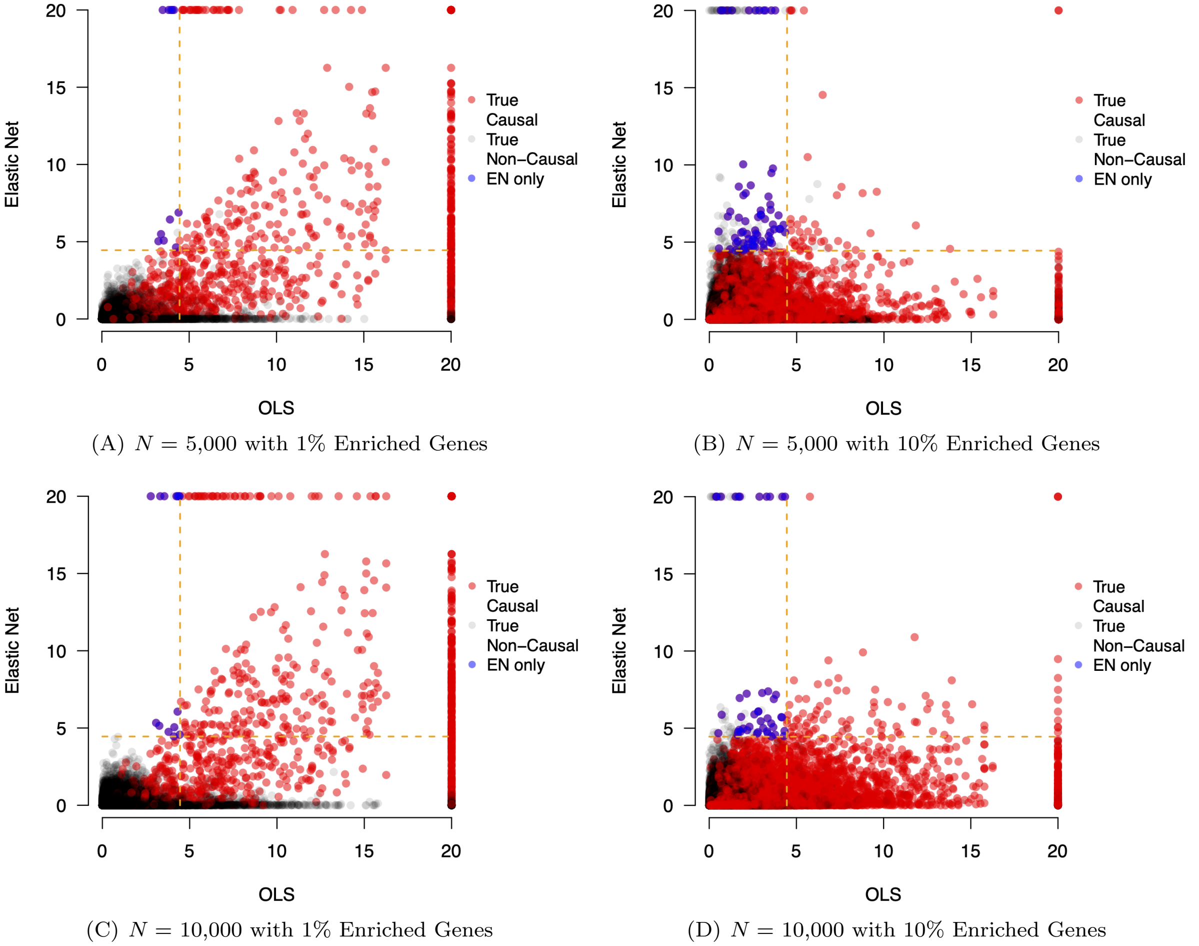

(A) $N$ = 5,000 with 1% Enriched Genes

(B) $N$ = 5,000 with 10% Enriched Genes

(C) $N$ = 10,000 with 1% Enriched Genes

(D) $N$ = 10,000 with 10% Enriched Genes

**Figure S17. Scatter plots assessing how regularization on SNP-level summary statistics affects the ability to identify enriched genes in simulations ($h^2 = 0.2$).** Here, the narrow-sense heritability of the simulated quantitative traits is $h^2 = 0.2$ and sample sizes are set to $N$ = 5,000 in **(A, B)** and $N$ = 10,000 in **(C, D)**. In each case, standard GWA summary statistics were computed by fitting a single-SNP univariate linear model (via ordinary least squares). Results are shown comparing the -$\log_{10}$ transformed gene-level $P$-values derived by gene-$\varepsilon$ with Elastic Net (EN) regularization on the y-axis and without regularization (labeled as OLS) on the x-axis. The horizontal and vertical dashed lines are marked at the Bonferonni-corrected threshold $P = 3.55 \times 10^{-5}$ corrected for the 1,408 genes on chromosome 1 from the UK Biobank genotype data. True positive causal genes used to generate the synthetic phenotypes are colored in red, while non-causal genes are given in grey. Genes in the top right quadrant are selected by both approaches. Genes in the top left and bottom right quadrants are uniquely identified by gene-$\varepsilon$-EN and gene-$\varepsilon$-OLS, respectively. To illustrate the importance of regularization on SNP-level summary statistics, we highlight the true positive genes only identified by gene-$\varepsilon$-EN in blue. Each plot combines results from 100 simulated replicates (see Section S2).

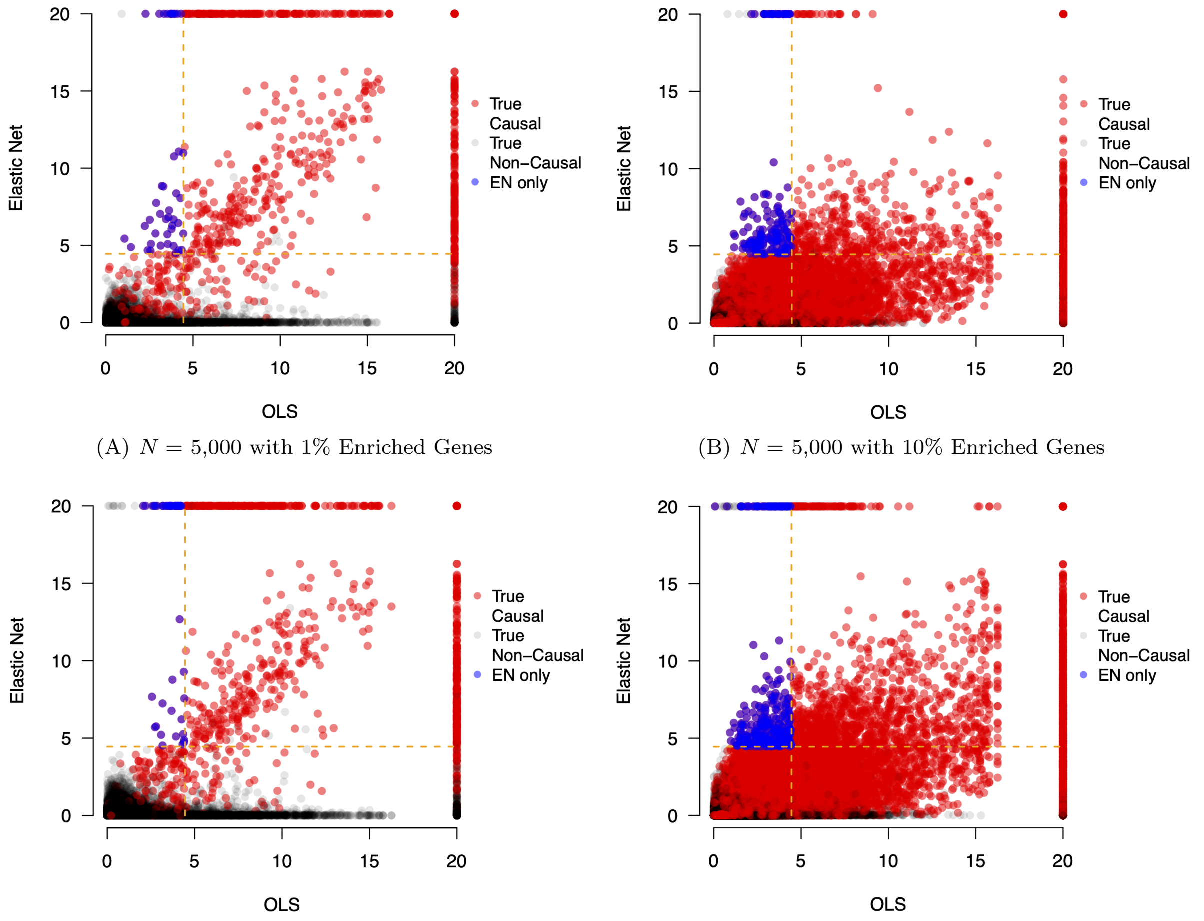

(A) $N$ = 5,000 with 1% Enriched Genes

(B) $N$ = 5,000 with 10% Enriched Genes

(C) $N$ = 10,000 with 1% Enriched Genes

(D) $N$ = 10,000 with 10% Enriched Genes

**Figure S18. Scatter plots assessing how regularization on SNP-level summary statistics affects the ability to identify enriched genes in simulations ($h^2 = 0.6$).** Here, the narrow-sense heritability of the simulated quantitative traits is $h^2 = 0.6$ and sample sizes are set to $N$ = 5,000 in **(A, B)** and $N$ = 10,000 in **(C, D)**. In each case, standard GWA summary statistics were computed by fitting a single-SNP univariate linear model (via ordinary least squares). Results are shown comparing the -$\log_{10}$ transformed gene-level $P$-values derived by gene-$\varepsilon$ with Elastic Net (EN) regularization on the y-axis and without regularization (labeled as OLS) on the x-axis. The horizontal and vertical dashed lines are marked at the Bonferonni-corrected threshold $P = 3.55 \times 10^{-5}$ corrected for the 1,408 genes on chromosome 1 from the UK Biobank genotype data. True positive causal genes used to generate the synthetic phenotypes are colored in red, while non-causal genes are given in grey. Genes in the top right quadrant are selected by both approaches. Genes in the top left and bottom right quadrants are uniquely identified by gene-$\varepsilon$-EN and gene-$\varepsilon$-OLS, respectively. To illustrate the importance of regularization on SNP-level summary statistics, we highlight the true positive genes only identified by gene-$\varepsilon$-EN in blue. Each plot combines results from 100 simulated replicates (see Section S2).

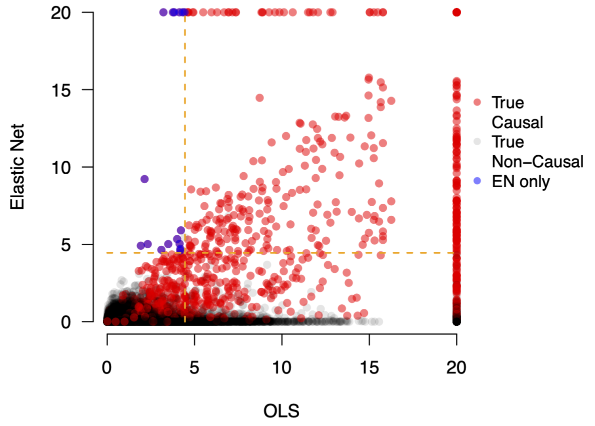

(A) $N$ = 5,000 with 1% Enriched Genes

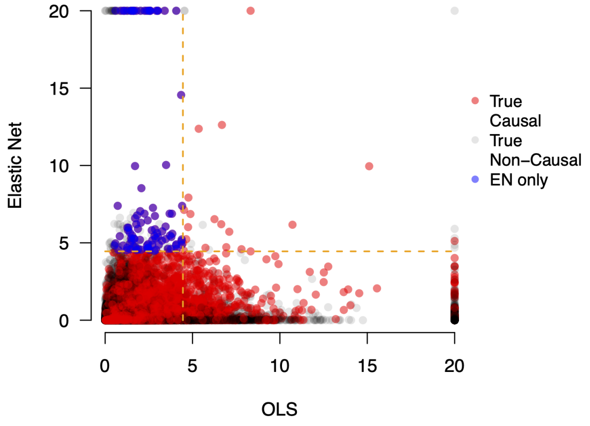

(B) $N$ = 5,000 with 10% Enriched Genes

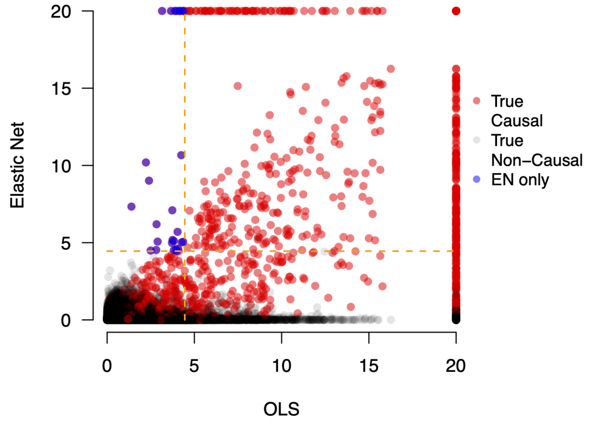

(C) $N$ = 10,000 with 1% Enriched Genes

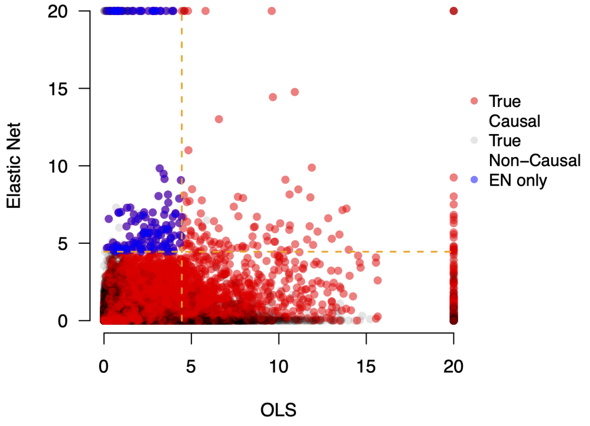

(D) $N$ = 10,000 with 10% Enriched Genes

**Figure S19. Scatter plots assessing how regularization on SNP-level summary statistics affects the ability to identify enriched genes in simulations with population stratification ($h^2$ = 0.2).** Here, the narrow-sense heritability of the simulated quantitative traits is $h^2 = 0.2$ and sample sizes are set to $N$ = 5,000 in **(A, B)** and $N$ = 10,000 in **(C, D)**. In this simulation, traits were generated while using the top five principal components (PCs) of the genotype matrix as covariates. GWA summary statistics were computed by fitting a single-SNP univariate linear model (via ordinary least squares) without any control for the additional structure. Results are shown comparing the -$\log_{10}$ transformed gene-level $P$-values derived by gene-$\varepsilon$ with Elastic Net (EN) regularization on the y-axis and without regularization (labeled as OLS) on the x-axis. The horizontal and vertical dashed lines are marked at the Bonferonni-corrected threshold $P = 3.55 \times 10^{-5}$ corrected for the 1,408 genes on chromosome 1 from the UK Biobank genotype data. True positive causal genes used to generate the synthetic phenotypes are colored in red, while non-causal genes are given in grey. Genes in the top right quadrant are selected by both approaches. Genes in the top left and bottom right quadrants are uniquely identified by gene-$\varepsilon$-EN and gene-$\varepsilon$-OLS, respectively. To illustrate the importance of regularization on SNP-level summary statistics, we highlight the true positive genes only identified by gene-$\varepsilon$-EN in blue. Each plot combines results from 100 simulated replicates (see Section S2).

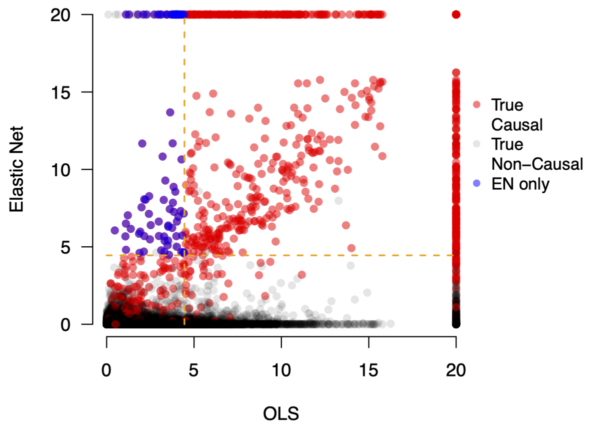

(A) $N$ = 5,000 with 1% Enriched Genes

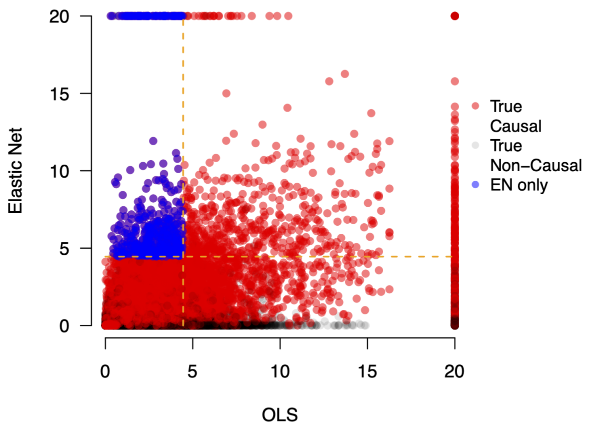

(B) $N$ = 5,000 with 10% Enriched Genes

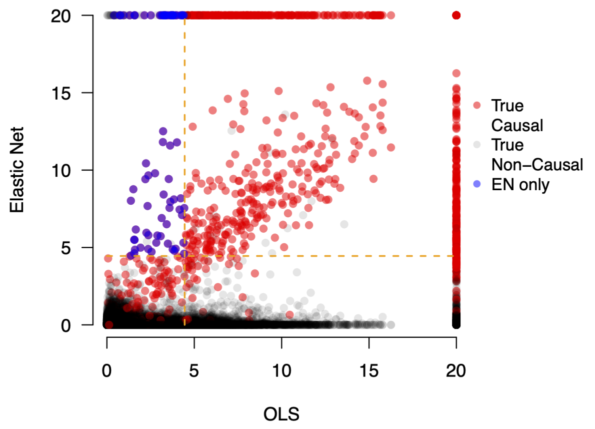

(C) $N$ = 10,000 with 1% Enriched Genes

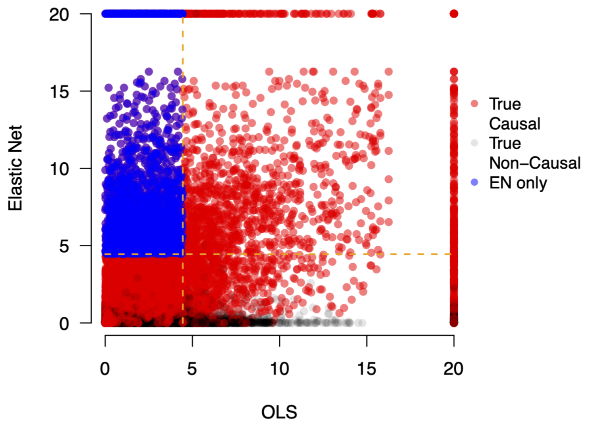

(D) $N$ = 10,000 with 10% Enriched Genes

**Figure S20. Scatter plots assessing how regularization on SNP-level summary statistics affects the ability to identify enriched genes in simulations with population stratification ($h^2 = 0.6$).** Here, the narrow-sense heritability of the simulated quantitative traits is $h^2 = 0.6$ and sample sizes are set to $N$ = 5,000 in **(A, B)** and $N$ = 10,000 in **(C, D)**. In this simulation, traits were generated while using the top five principal components (PCs) of the genotype matrix as covariates. GWA summary statistics were computed by fitting a single-SNP univariate linear model (via ordinary least squares) without any control for the additional structure. Results are shown comparing the -$\log_{10}$ transformed gene-level $P$-values derived by gene-$\varepsilon$ with Elastic Net (EN) regularization on the y-axis and without regularization (labeled as OLS) on the x-axis. The horizontal and vertical dashed lines are marked at the Bonferonni-corrected threshold $P = 3.55 \times 10^{-5}$ corrected for the 1,408 genes on chromosome 1 from the UK Biobank genotype data. True positive causal genes used to generate the synthetic phenotypes are colored in red, while non-causal genes are given in grey. Genes in the top right quadrant are selected by both approaches. Genes in the top left and bottom right quadrants are uniquely identified by gene-$\varepsilon$-EN and gene-$\varepsilon$-OLS, respectively. To illustrate the importance of regularization on SNP-level summary statistics, we highlight the true positive genes only identified by gene-$\varepsilon$-EN in blue. Each plot combines results from 100 simulated replicates (see Section S2).

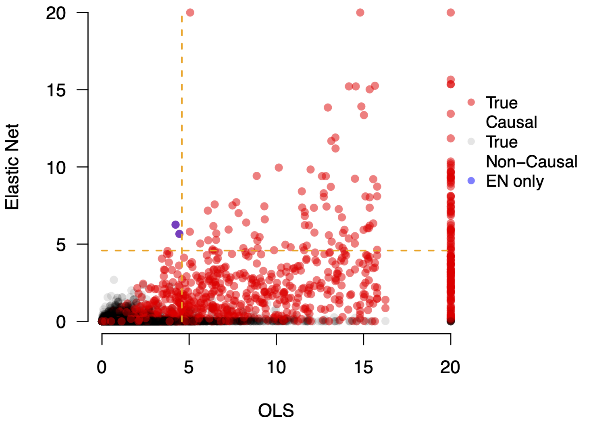

(A) $N$ = 5,000 with 1% Enriched Genes

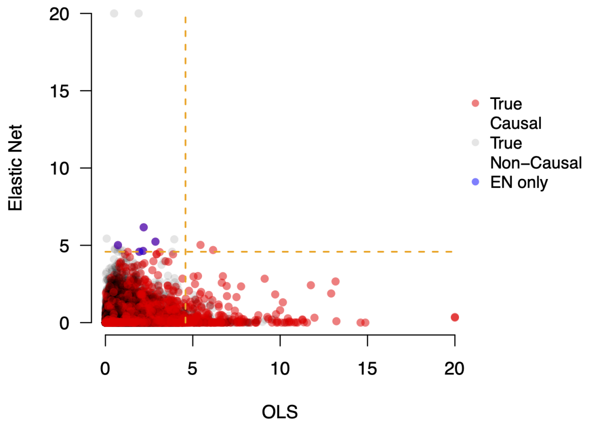

(B) $N$ = 5,000 with 10% Enriched Genes

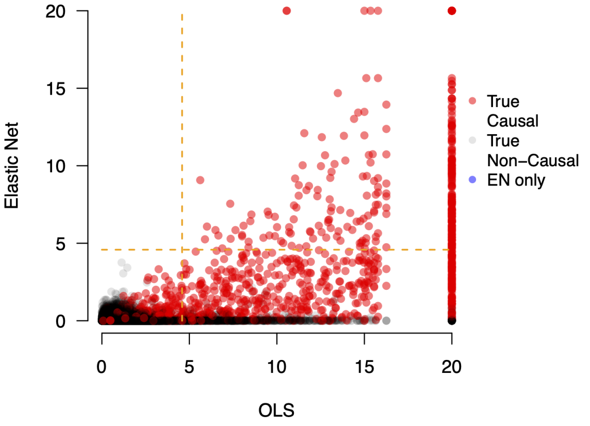

(C) $N$ = 10,000 with 1% Enriched Genes

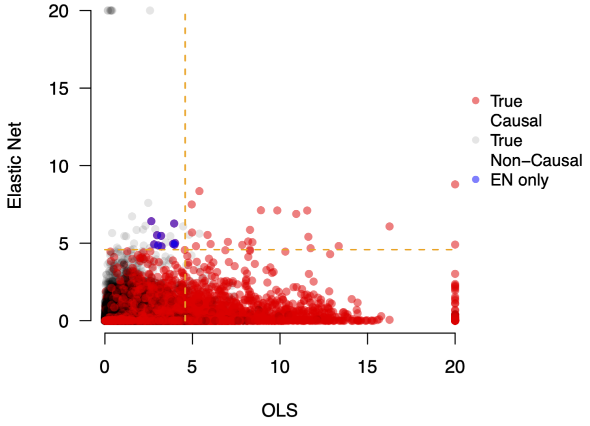

(D) $N$ = 10,000 with 10% Enriched Genes

**Figure S21. Scatter plots assessing how regularization on SNP-level summary statistics affects the ability to identify enriched genes in simulations with gene boundaries augmented by a 50 kilobase (kb) buffer ($h^2 = 0.2$).** Here, the narrow-sense heritability of the simulated quantitative traits is $h^2 = 0.2$ and sample sizes are set to $N$ = 5,000 in **(A, B)** and $N$ = 10,000 in **(C, D)**. In each case, standard GWA summary statistics were computed by fitting a single-SNP univariate linear model (via ordinary least squares). Results are shown comparing the -$\log_{10}$ transformed gene-level $P$-values derived by gene-$\varepsilon$ with Elastic Net (EN) regularization on the y-axis and without regularization (labeled as OLS) on the x-axis. The horizontal and vertical dashed lines are marked at the Bonferonni-corrected threshold $P = 2.61 \times 10^{-5}$ corrected for the 1,916 genes on chromosome 1 from the UK Biobank genotype data. True positive causal genes used to generate the synthetic phenotypes are colored in red, while non-causal genes are given in grey. Genes in the top right quadrant are selected by both approaches. Genes in the top left and bottom right quadrants are uniquely identified by gene-$\varepsilon$-EN and gene-$\varepsilon$-OLS, respectively. To illustrate the importance of regularization on SNP-level summary statistics, we highlight the true positive genes only identified by gene-$\varepsilon$-EN in blue. Each plot combines results from 100 simulated replicates (see Section S2).

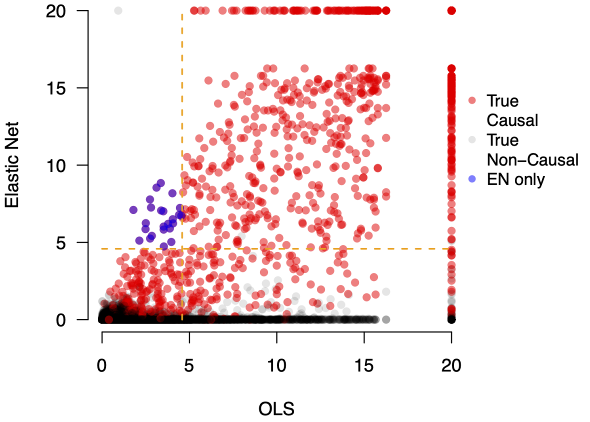

(A) $N$ = 5,000 with 1% Enriched Genes

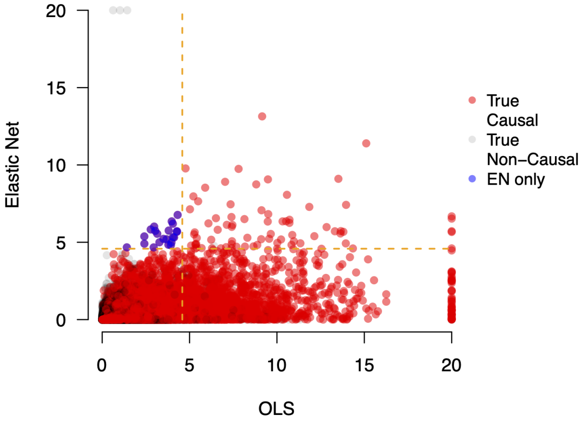

(B) $N$ = 5,000 with 10% Enriched Genes

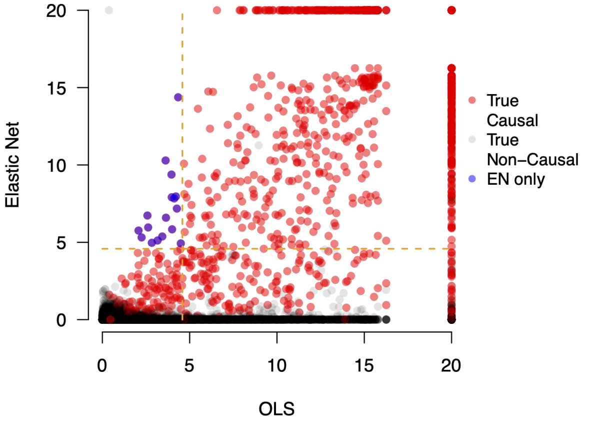

(C) $N$ = 10,000 with 1% Enriched Genes

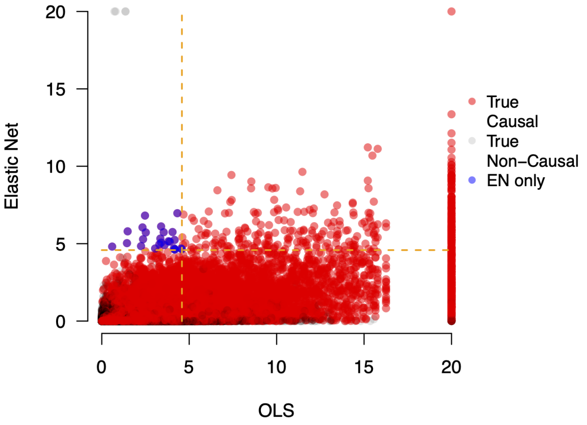

(D) $N$ = 10,000 with 10% Enriched Genes

**Figure S22. Scatter plots assessing how regularization on SNP-level summary statistics affects the ability to identify enriched genes in simulations with gene boundaries augmented by a 50 kilobase (kb) buffer ($h^2$ = 0.6).** Here, the narrow-sense heritability of the simulated quantitative traits is $h^2 = 0.6$ and sample sizes are set to $N$ = 5,000 in **(A, B)** and $N$ = 10,000 in **(C, D)**. In each case, standard GWA summary statistics were computed by fitting a single-SNP univariate linear model (via ordinary least squares). Results are shown comparing the -$\log_{10}$ transformed gene-level $P$-values derived by gene-$\varepsilon$ with Elastic Net (EN) regularization on the y-axis and without regularization (labeled as OLS) on the x-axis. The horizontal and vertical dashed lines are marked at the Bonferonni-corrected threshold $P = 2.61 \times 10^{-5}$ corrected for the 1,916 genes on chromosome 1 from the UK Biobank genotype data. True positive causal genes used to generate the synthetic phenotypes are colored in red, while non-causal genes are given in grey. Genes in the top right quadrant are selected by both approaches. Genes in the top left and bottom right quadrants are uniquely identified by gene-$\varepsilon$-EN and gene-$\varepsilon$-OLS, respectively. To illustrate the importance of regularization on SNP-level summary statistics, we highlight the true positive genes only identified by gene-$\varepsilon$-EN in blue. Each plot combines results from 100 simulated replicates (see Section S2).

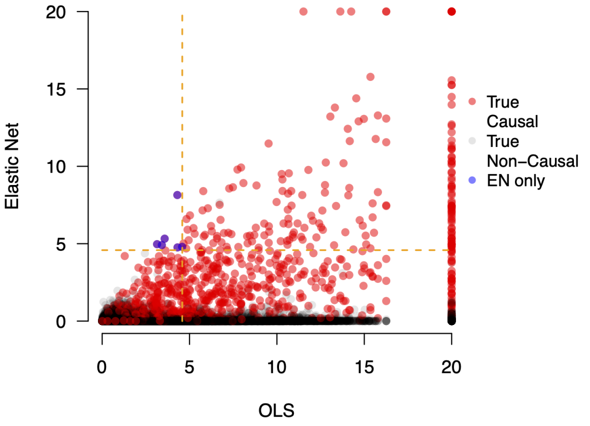

(A) $N = 5{,}000$ with 1% Enriched Genes

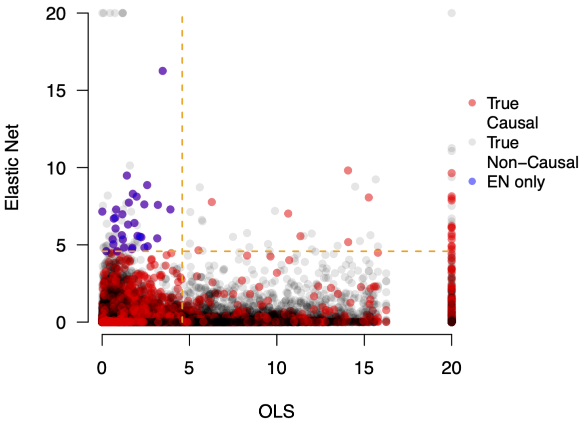

(B) $N = 5{,}000$ with 10% Enriched Genes

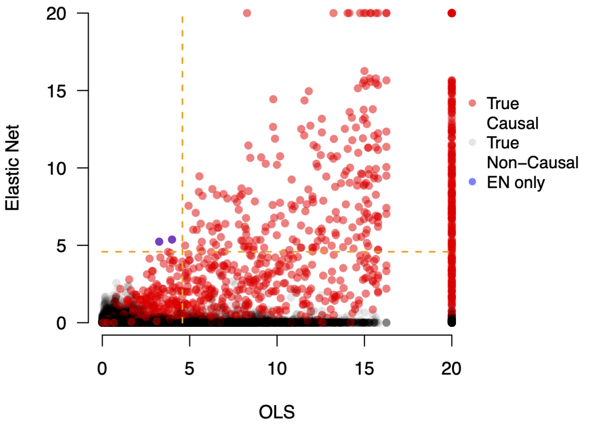

(C) $N = 10{,}000$ with 1% Enriched Genes

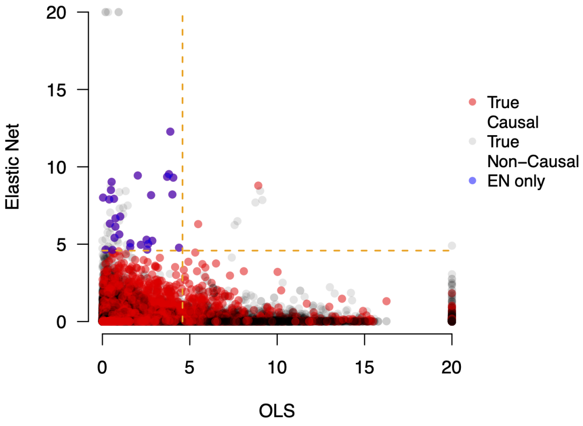

(D) $N = 10{,}000$ with 10% Enriched Genes

**Figure S23. Scatter plots assessing how regularization on SNP-level summary statistics affects the ability to identify enriched genes in simulations with gene boundaries augmented by a 50 kilobase (kb) buffer and with population stratification ($h^2 = 0.2$).** Here, the narrow-sense heritability of the simulated quantitative traits is $h^2 = 0.2$ and sample sizes are set to $N = 5{,}000$ in **(A, B)** and $N = 10{,}000$ in **(C, D)**. In this simulation, traits were generated while using the top five principal components (PCs) of the genotype matrix as covariates. GWA summary statistics were computed by fitting a single-SNP univariate linear model (via ordinary least squares) without any control for the additional structure. Results are shown comparing the -$\log_{10}$ transformed gene-level $P$-values derived by gene-$\varepsilon$ with Elastic Net (EN) regularization on the y-axis and without regularization (labeled as OLS) on the x-axis. The horizontal and vertical dashed lines are marked at the Bonferonni-corrected threshold $P = 2.61 \times 10^{-5}$ corrected for the 1,916 genes on chromosome 1 from the UK Biobank genotype data. True positive causal genes used to generate the synthetic phenotypes are colored in red, while non-causal genes are given in grey. Genes in the top right quadrant are selected by both approaches. Genes in the top left and bottom right quadrants are uniquely identified by gene-$\varepsilon$-EN and gene-$\varepsilon$-OLS, respectively. To illustrate the importance of regularization on SNP-level summary statistics, we highlight the true positive genes only identified by gene-$\varepsilon$-EN in blue. Each plot combines results from 100 simulated replicates (see Section S2).

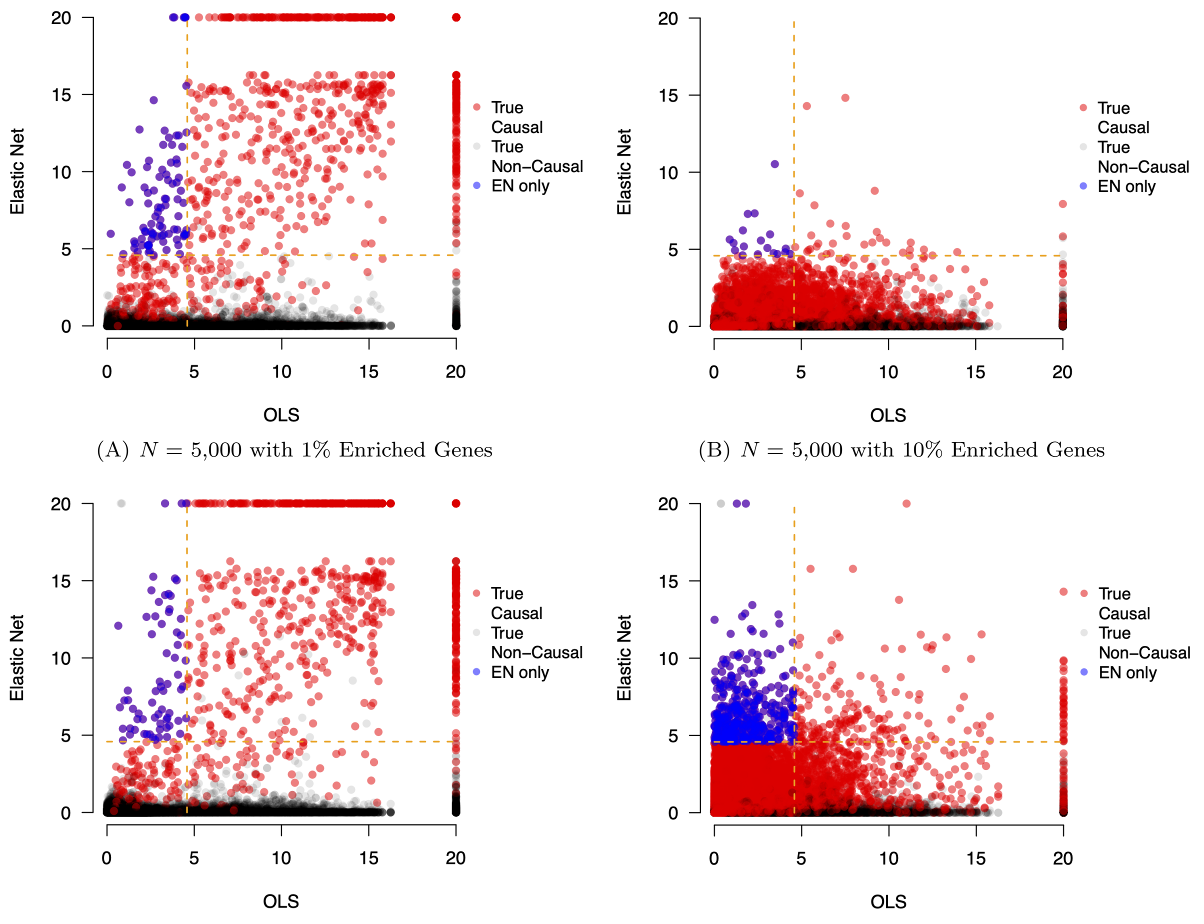

(A) $N$ = 5,000 with 1% Enriched Genes

(B) $N$ = 5,000 with 10% Enriched Genes

(C) $N$ = 10,000 with 1% Enriched Genes

(D) $N$ = 10,000 with 10% Enriched Genes

**Figure S24. Scatter plots assessing how regularization on SNP-level summary statistics affects the ability to identify enriched genes in simulations with gene boundaries augmented by a 50 kilobase (kb) buffer and with population stratification ($h^2 = 0.6$).** Here, the narrow-sense heritability of the simulated quantitative traits is $h^2 = 0.6$ and sample sizes are set to $N =$ 5,000 in **(A, B)** and $N =$ 10,000 in **(C, D)**. In this simulation, traits were generated while using the top five principal components (PCs) of the genotype matrix as covariates. GWA summary statistics were computed by fitting a single-SNP univariate linear model (via ordinary least squares) without any control for the additional structure. Results are shown comparing the -$\log_{10}$ transformed gene-level $P$-values derived by gene-$\varepsilon$ with Elastic Net (EN) regularization on the y-axis and without regularization (labeled as OLS) on the x-axis. The horizontal and vertical dashed lines are marked at the Bonferonni-corrected threshold $P = 2.61 \times 10^{-5}$ corrected for the 1,916 genes on chromosome 1 from the UK Biobank genotype data. True positive causal genes used to generate the synthetic phenotypes are colored in red, while non-causal genes are given in grey. Genes in the top right quadrant are selected by both approaches. Genes in the top left and bottom right quadrants are uniquely identified by gene-$\varepsilon$-EN and gene-$\varepsilon$-OLS, respectively. To illustrate the importance of regularization on SNP-level summary statistics, we highlight the true positive genes only identified by gene-$\varepsilon$-EN in blue. Each plot combines results from 100 simulated replicates (see Section S2).

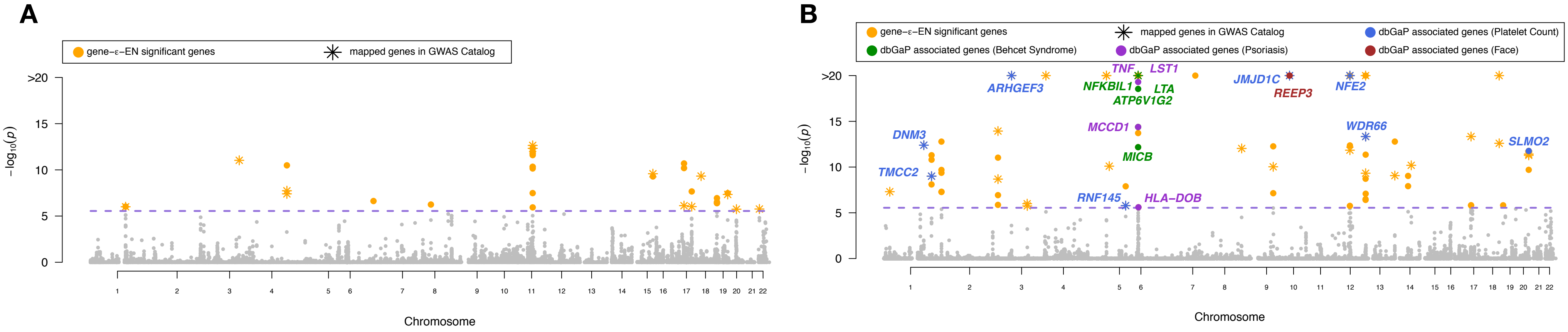

**C**

| | $p$ value | $q$ value | Odds.ratio | Combined score | # of sig. genes in dbGaP |
|---|---|---|---|---|---|
| Body Height | 5.20e-04 | 1.79e-01 | 7.42 | 56.12 | 5 |
| Macular Degeneration | 1.14e-02 | 1.97e-01 | 14.53 | 98.43 | 3 |
| Prion Diseases | 1.22e-02 | 1.00 | 81.63 | 359.78 | 1 |
| Neuroblastoma | 1.35e-02 | 1.00 | 11.32 | 48.73 | 2 |
| Smoking | 2.08e-02 | 1.00 | 47.62 | 184.41 | 1 |
| Stroke | 9.07e-02 | 1.00 | 3.95 | 9.49 | 2 |
| Prostatic Neoplasms | 1.23e-01 | 1.00 | 7.62 | 15.95 | 1 |
| Creatinine | 1.49e-01 | 1.00 | 6.21 | 11.82 | 1 |
| Respiratory Function Tests | 2.33e-01 | 1.00 | 3.78 | 5.51 | 1 |
| – | – | – | – | – | – |

**D**

| | $p$ value | $q$ value | Odds.ratio | Combined score | # of sig. genes in dbGaP |
|---|---|---|---|---|---|
| Platelet Count | 2.16e-11 | 7.45e-09 | 40.04 | 983.32 | 8 |
| Behcet Syndrome | 2.59e-06 | 4.47e-04 | 22.90 | 294.64 | 5 |
| Psoriasis | 1.74e-05 | 2.00e-03 | 25.74 | 282.13 | 4 |
| Face | 8.06e-05 | 6.96e-03 | 135.14 | 1273.71 | 2 |
| Diabetes Mellitus, Type 1 | 2.71e-03 | 1.87e-01 | 10.81 | 63.89 | 3 |
| Dengue Hemorrhagic Fever | 1.11e-02 | 6.36e-01 | 90.09 | 405.81 | 1 |
| Hearing Loss | 1.11e-02 | 5.45e-01 | 90.09 | 405.81 | 1 |
| Erythrocytes | 1.59e-02 | 6.85e-01 | 10.40 | 43.05 | 2 |
| Body Mass Index | 2.29e-02 | 8.78e-01 | 3.09 | 11.68 | 5 |
| Arteries | 4.30e-02 | 1.00 | 6.07 | 19.11 | 2 |

**Figure S25. Gene-level association results from applying gene-$\varepsilon$ to body height (panels A and C) and mean platelet volume (MPV; panels B and D), assayed in European-ancestry individuals in the UK Biobank with UCSC RefSeq gene boundaries augmented by a 50 kilobase (kb) buffer.** Body height has been estimated to have a narrow-sense heritability $h^2$ in the range of 0.45 to 0.80 [19–28]; while, MPV has been estimated to have $h^2$ between 0.50 and 0.70 [22, 23, 29]. Manhattan plots of gene-$\varepsilon$ gene-level association $P$-values using Elastic Net regularized effect sizes for **(A)** body height and **(B)** MPV. The purple dashed line indicates a log-transformed Bonferroni-corrected significance threshold ($P = 2.83 \times 10^{-6}$ correcting for 17,680 autosomal genes analyzed). We color code all significant genes identified by gene-$\varepsilon$ in orange, and annotate genes overlapping with the database of Genotypes and Phenotypes (dbGaP). In **(C)** and **(D)**, we conduct gene set enrichment analysis using Enrichr [30, 31] to identify dbGaP categories enriched for significant gene-level associations reported by gene-$\varepsilon$. We highlight categories with $Q$-values (i.e., false discovery rates) less than 0.05 and annotate corresponding genes in the Manhattan plots in **(A)** and **(B)**, respectively. For height, the most enriched dbGAP category is "Body Height", with 5 of the genes identified by gene-$\varepsilon$ appearing in this category. For MPV, the four significant dbGAP categories are "Platelet Count", "Behcet Syndrome", "Psoriasis", and "Face" — all of which have been connected to trait [32–36].

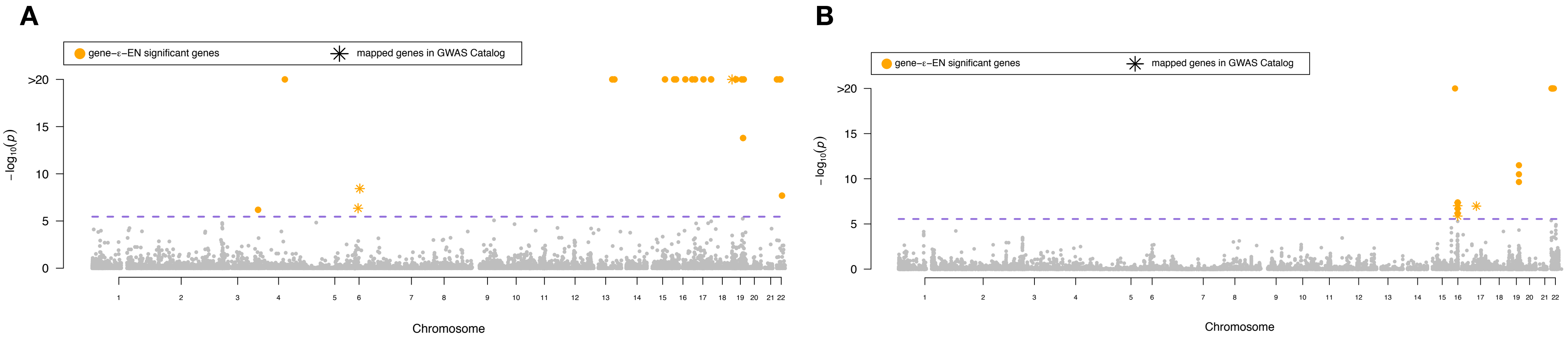

**C**

| | *p* value | *q* value | Odds.ratio | Combined score | # of sig. genes in dbGaP |
|---|---|---|---|---|---|
| Adiposity | 4.79e-03 | 1.00 | 208.33 | 1112.68 | 1 |
| Metabolic Syndrome X | 2.73e-02 | 1.00 | 36.23 | 130.53 | 1 |
| Body Mass Index | 9.59e-02 | 1.00 | 3.81 | 8.94 | 2 |
| — | — | — | — | — | — |
| — | — | — | — | — | — |
| — | — | — | — | — | — |
| — | — | — | — | — | — |
| — | — | — | — | — | — |
| — | — | — | — | — | — |
| — | — | — | — | — | — |

**D**

| | *p* value | *q* value | Odds.ratio | Combined score | # of sig. genes in dbGaP |
|---|---|---|---|---|---|
| gamma-Glutamyltransferase | 1.85e-02 | 1.00 | 53.48 | 213.24 | 1 |
| Alkaline Phosphatase | 1.94e-02 | 1.00 | 51.15 | 201.72 | 1 |
| Arteries | 7.30e-02 | 1.00 | 13.22 | 34.59 | 1 |
| Body Mass Index | 3.13e-01 | 1.00 | 2.69 | 3.13 | 1 |
| — | — | — | — | — | — |
| — | — | — | — | — | — |
| — | — | — | — | — | — |
| — | — | — | — | — | — |
| — | — | — | — | — | — |
| — | — | — | — | — | — |

**Figure S26. Gene-level association results from applying gene-$\varepsilon$ to body mass index (BMI), assayed in European-ancestry individuals in the UK Biobank.** BMI has been estimated to have a narrow-sense heritability $h^2$ ranging from 0.25 to 0.60 [20, 22, 23, 25, 26, 28, 37–40]. Manhattan plots of gene-$\varepsilon$ gene-level association $P$-values using Elastic Net regularized effect sizes when gene boundaries are defined by **(A)** using UCSC annotations directly, and **(B)** augmenting the gene boundaries by adding SNPs within a ±50kb buffer. The purple dashed line indicates a log-transformed Bonferroni-corrected significance threshold ($P = 3.49 \times 10^{-6}$ and $P = 2.83 \times 10^{-6}$ correcting for the 14,322 and 17,680 autosomal genes analyzed, respectively). We color code all significant genes identified by gene-$\varepsilon$ in orange, and annotate genes previously associated with BMI in the database of Genotypes and Phenotypes (dbGaP). In **(C)** and **(D)**, we conduct gene set enrichment analysis using Enrichr [30, 31] to identify dbGaP categories enriched for significant gene-level associations reported by gene-$\varepsilon$ in **(A)** and **(B)**, respectively. While many of the scored categories are biologically related to BMI (e.g., "Body Mass Index", "Adiposity", and "Arteries") [41–44], none of them had $Q$-values (i.e., false discovery rates) less than 0.05.

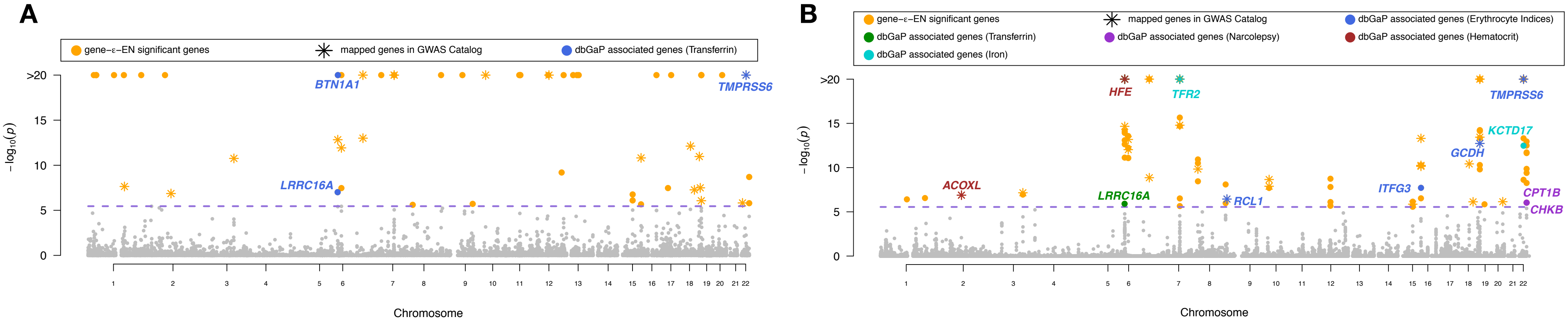

**C**

| | $p$ value | $q$ value | Odds.ratio | Combined score | # of sig. genes in dbGaP |
|---|---|---|---|---|---|
| Transferrin | 3.56e-06 | 1.23e-03 | 95.69 | 1200.47 | 3 |
| Hematocrit | 7.02e-04 | 1.21e-01 | 17.26 | 125.30 | 3 |
| Iron-Regulatory Proteins | 8.53e-03 | 9.80e-01 | 16.96 | 557.27 | 1 |
| Erythrocyte Indices | 9.30e-03 | 8.02e-01 | 13.76 | 64.36 | 2 |
| Iron | 9.67e-03 | 6.67e-01 | 6.84 | 31.71 | 3 |
| Platelet Count | 1.04e-02 | 5.97e-01 | 13.00 | 59.35 | 2 |
| Esophageal Neoplasms | 2.54e-02 | 1.00 | 38.99 | 143.25 | 1 |
| Leukemia, Lymphocytic, Chronic, B-Cell | 2.81e-02 | 1.00 | 35.09 | 125.28 | 1 |
| Glomerulonephritis, IGA | 3.09e-02 | 1.00 | 31.90 | 110.89 | 1 |
| Hemoglobin A, Glycosylated | 5.16e-02 | 1.00 | 5.48 | 16.25 | 2 |

**D**

| | $p$ value | $q$ value | Odds.ratio | Combined score | # of sig. genes in dbGaP |
|---|---|---|---|---|---|
| Erythrocyte Indices | 3.48e-06 | 1.20e-03 | 21.55 | 270.84 | 5 |
| Transferrin | 1.46e-05 | 2.53e-03 | 59.94 | 667.23 | 3 |
| Narcolepsy | 6.12e-05 | 7.04e-03 | 146.52 | 1421.34 | 2 |
| Hematocrit | 1.72e-04 | 1.48e-02 | 14.41 | 124.95 | 4 |
| Iron | 6.93e-04 | 4.78e-02 | 7.14 | 51.91 | 5 |
| Ferritins | 1.36e-02 | 7.81e-01 | 73.26 | 314.91 | 1 |
| carbohydrate-deficient transferrin | 1.36e-02 | 6.70e-01 | 73.26 | 314.91 | 1 |
| Iron-Regulatory Proteins | 1.36e-02 | 5.86e-01 | 73.26 | 314.91 | 1 |
| Blood Pressure | 1.75e-02 | 6.70e-01 | 2.90 | 11.75 | 6 |
| Prostate-Specific Antigen | 1.81e-02 | 6.24e-01 | 54.95 | 220.50 | 1 |

**Figure S27. Gene-level association results from applying gene-$\varepsilon$ to mean corpuscular volume (MCV), assayed in European-ancestry individuals in the UK Biobank.** MCV has been estimated to have a narrow-sense heritability $h^2$ in the range of 0.20 to 0.60 [22, 23, 45, 46]. Manhattan plots of gene-$\varepsilon$ gene-level association $P$-values using Elastic Net regularized effect sizes when gene boundaries are defined by **(A)** using UCSC annotations directly, and **(B)** augmenting the gene boundaries by adding SNPs within a ±50kb buffer. The purple dashed line indicates a log-transformed Bonferroni-corrected significance threshold ($P = 3.49 \times 10^{-6}$ and $P = 2.83 \times 10^{-6}$ correcting for the 14,322 and 17,680 autosomal genes analyzed, respectively). We color code all significant genes identified by gene-$\varepsilon$ in orange, and annotate genes previously associated with MCV in the database of Genotypes and Phenotypes (dbGaP). In **(C)** and **(D)**, we conduct gene set enrichment analysis using Enrichr [30,31] to identify dbGaP categories enriched for significant gene-level associations reported by gene-$\varepsilon$. We highlight categories with $Q$-values (i.e., false discovery rates) less than 0.05 and annotate corresponding genes in the Manhattan plots in **(A)** and **(B)**, respectively. The dbGAP categories significantly enriched for gene-level associations with MCV included "Transferrin", "Erythrocyte Indices", "Hematocrit", "Narcolepsy", and "Iron" — all of which have been connected to trait [34, 47–53].

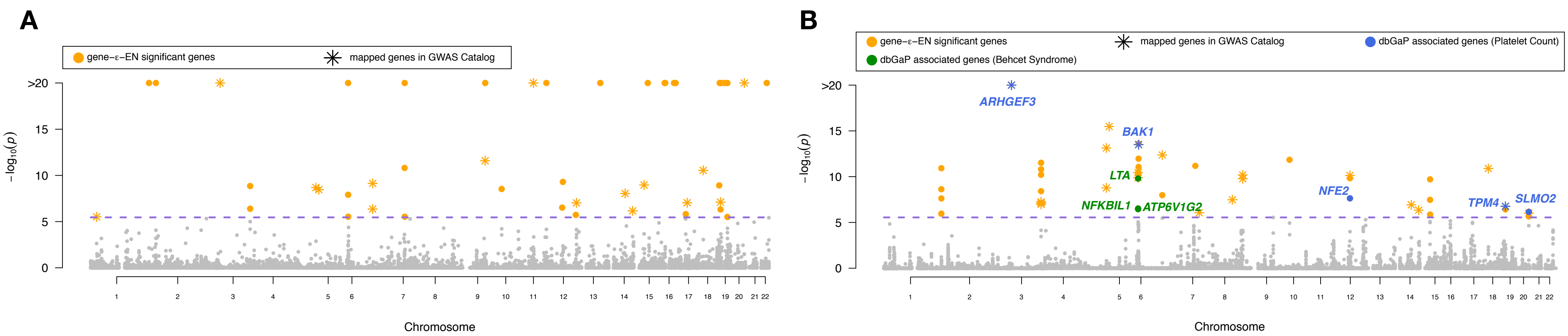

**C**

| | $p$ value | $q$ value | Odds.ratio | Combined score | # of sig. genes in dbGaP |
|---|---|---|---|---|---|
| Platelet Count | 2.95e-04 | 1.02e-01 | 23.15 | 188.14 | 3 |
| Smoking | 3.67e-04 | 6.32e-02 | 69.44 | 549.39 | 2 |
| Hearing Loss | 7.18e-03 | 8.26e-01 | 138.89 | 685.56 | 1 |
| Esophageal Neoplasms | 2.14e-02 | 1.00 | 46.30 | 177.98 | 1 |
| Macular Degeneration | 3.26e-02 | 1.00 | 7.06 | 24.17 | 2 |
| Cardiomegaly | 3.54e-02 | 1.00 | 27.78 | 92.80 | 1 |
| Pulse | 8.07e-02 | 1.00 | 11.90 | 29.96 | 1 |
| Body Mass Index | 8.72e-02 | 1.00 | 2.86 | 6.98 | 3 |
| Arthritis, Rheumatoid | 1.32e-01 | 1.00 | 7.06 | 14.28 | 1 |
| Coronary Disease | 1.79e-01 | 1.00 | 5.08 | 8.74 | 1 |

**D**

| | $p$ value | $q$ value | Odds.ratio | Combined score | # of sig. genes in dbGaP |
|---|---|---|---|---|---|
| Platelet Count | 2.07e-07 | 7.13e-05 | 37.79 | 581.69 | 5 |
| Smoking | 3.82e-04 | 6.59e-02 | 68.03 | 535.37 | 2 |
| Behcet Syndrome | 4.08e-04 | 4.69e-02 | 20.75 | 161.97 | 3 |
| Psoriasis | 4.76e-03 | 4.10e-01 | 19.44 | 103.95 | 2 |
| Erythrocyte Indices | 6.94e-03 | 4.79e-01 | 16.01 | 79.55 | 2 |
| Hearing Loss | 7.33e-03 | 4.22e-01 | 136.05 | 668.77 | 1 |
| Testicular Neoplasms | 1.22e-02 | 6.01e-01 | 81.63 | 359.76 | 1 |
| Cardiomegaly | 3.61e-02 | 1.00 | 27.21 | 90.35 | 1 |
| Heart Rate | 5.54e-02 | 1.00 | 5.27 | 15.24 | 2 |
| Aorta | 8.01e-02 | 1.00 | 12.00 | 30.31 | 1 |

**Figure S28. Gene-level association results from applying gene-$\varepsilon$ to platelet count (PLC), assayed in European-ancestry individuals in the UK Biobank.** PLC has been estimated to have a narrow-sense heritability $h^2$ ranging from 0.55 to 0.80 [22, 23, 29]. Manhattan plots of gene-$\varepsilon$ gene-level association $P$-values using Elastic Net regularized effect sizes when gene boundaries are defined by **(A)** using UCSC annotations directly, and **(B)** augmenting the gene boundaries by adding SNPs within a ±50kb buffer. The purple dashed line indicates a log-transformed Bonferroni-corrected significance threshold ($P = 3.49 \times 10^{-6}$ and $P = 2.83 \times 10^{-6}$ correcting for the 14,322 and 17,680 autosomal genes analyzed, respectively). We color code all significant genes identified by gene-$\varepsilon$ in orange, and annotate genes previously associated with PLC in the database of Genotypes and Phenotypes (dbGaP). In **(C)** and **(D)**, we conduct gene set enrichment analysis using Enrichr [30, 31] to identify dbGaP categories enriched for significant gene-level associations reported by gene-$\varepsilon$. We highlight categories with $Q$-values (i.e., false discovery rates) less than 0.05 and annotate corresponding genes in the Manhattan plots in **(A)** and **(B)**, respectively. The most significant dbGAP category is "Platelet Count" for both SNP-to-gene annotation schemes. The other significant dbGAP category was "Smoking" which has been previously connected to PLC [36, 54, 55].

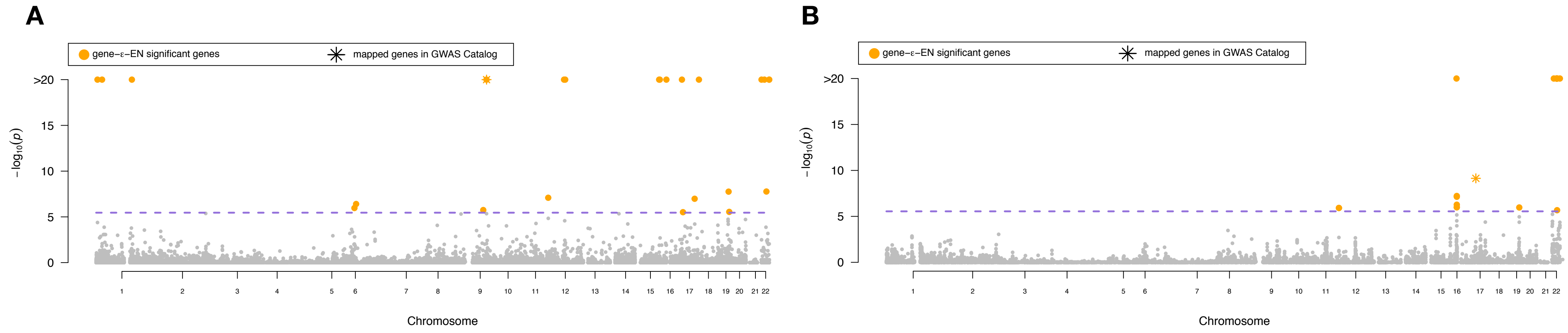

**C**

| | $p$ value | $q$ value | Odds.ratio | Combined score | # of sig. genes in dbGaP |
|---|---|---|---|---|---|
| Adiposity | 4.99e-03 | 1.00 | 200.00 | 1060.03 | 1 |
| Glomerulonephritis, IGA | 1.37e-02 | 1.00 | 72.73 | 312.20 | 1 |
| Metabolic Syndrome X | 2.84e-02 | 1.00 | 34.78 | 123.91 | 1 |
| Inflammatory Bowel Diseases | 4.29e-02 | 1.00 | 22.86 | 71.99 | 1 |
| Schizophrenia | 9.77e-02 | 1.00 | 9.76 | 22.69 | 1 |
| Body Mass Index | 1.03e-01 | 1.00 | 3.66 | 8.33 | 2 |
| Macular Degeneration | 1.38e-01 | 1.00 | 6.78 | 13.45 | 1 |
| Stroke | 3.05e-01 | 1.00 | 2.77 | 3.29 | 1 |
| — | — | — | — | — | — |
| — | — | — | — | — | — |

**D**

| | $p$ value | $q$ value | Odds.ratio | Combined score | # of sig. genes in dbGaP |
|---|---|---|---|---|---|
| Dermatitis, Atopic | 3.79e-03 | 1.00 | 263.16 | 1466.8 | 1 |
| gamma-Glutamyltransferase | 2.07e-02 | 1.00 | 47.85 | 185.52 | 1 |
| Alkaline Phosphatase | 2.16e-02 | 1.00 | 45.77 | 175.44 | 1 |
| Inflammatory Bowel Diseases | 3.27e-02 | 1.00 | 30.08 | 102.83 | 1 |
| Body Mass Index | 3.43e-01 | 1.00 | 2.41 | 2.58 | 1 |
| — | — | — | — | — | — |
| — | — | — | — | — | — |
| — | — | — | — | — | — |
| — | — | — | — | — | — |
| — | — | — | — | — | — |

**Figure S29. Gene-level association results from applying gene-$\varepsilon$ to waist-hip ratio (WHR), assayed in European-ancestry individuals in the UK Biobank.** WHR has been estimated to have a narrow-sense heritability $h^2$ ranging from 0.10 to 0.25 [20, 22, 24, 37, 40, 56]. Manhattan plots of gene-$\varepsilon$ gene-level association $P$-values using Elastic Net regularized effect sizes when gene boundaries are defined by **(A)** using UCSC annotations directly, and **(B)** augmenting the gene boundaries by adding SNPs within a $\pm$50kb buffer. The purple dashed line indicates a log-transformed Bonferroni-corrected significance threshold ($P = 3.49 \times 10^{-6}$ and $P = 2.83 \times 10^{-6}$ correcting for the 14,322 and 17,680 autosomal genes analyzed, respectively). We color code all significant genes identified by gene-$\varepsilon$ in orange, and annotate genes previously associated with WHR in the database of Genotypes and Phenotypes (dbGaP). In **(C)** and **(D)**, we conduct gene set enrichment analysis using Enrichr [30, 31] to identify dbGaP categories enriched for significant gene-level associations reported by gene-$\varepsilon$ in **(A)** and **(B)**, respectively. While many of the scored categories are biologically related to WHR (e.g., "Body Mass Index", "Adiposity", and "Inflammatory Bowel Diseases") [57, 58], none of them had $Q$-values (i.e., false discovery rates) less than 0.05.

### Supplementary Tables

| | | *gene-ε Approaches* | | | |
|---|---|---|---|---|---|
| **Causal Genes** | **Metric** | **OLS** | **RR** | **EN** | **LASSO** |
| 1% | Power | 0.572 (0.149) | 0.358 (0.131) | 0.295 (0.111) | 0.296 (0.113) |
| | FDR | 0.415 (0.164) | 0.652 (0.314) | **0.007 (0.038)** | 0.017 (0.074) |
| 10% | Power | **0.072 (0.102)** | 0.044 (0.026) | 0.013 (0.020) | 0.011 (0.011) |
| | FDR | 0.187 (0.184) | 0.468 (0.310) | 0.470 (0.392) | 0.403 (0.401) |

| | | *Other Methods* | | | | |
|---|---|---|---|---|---|---|
| **Causal Genes** | **Metric** | **PEGASUS** | **VEGAS** | **RSS** | **SKAT** | **MAGMA** |
| 1% | Power | 0.569 (0.151) | 0.617 (0.152) | 0.487 (0.134) | 0.473 (0.141) | **0.623 (0.161)** |
| | FDR | 0.408 (0.170) | 0.432 (0.159) | 0.074 (0.103) | 0.374 (0.186) | 0.456 (0.160) |
| 10% | Power | 0.047 (0.014) | 0.028 (0.013) | 0.014 (0.007) | 0.036 (0.012) | 0.045 (0.015) |
| | FDR | 0.126 (0.121) | 0.184 (0.183) | **0.059 (0.194)** | 0.128 (0.140) | 0.136 (0.144) |

**Table S1. Empirical power and false discovery rates (FDR) for detecting enriched genes (genes containing at least one causal SNP) after correcting for multiple hypothesis testing in simulations ($N$ = 5,000; $h^2 = 0.2$).** We computed standard GWA SNP-level effect sizes (estimated using ordinary least squares) as input to each method listed. We show the power of gene-$\varepsilon$ to identify enriched genes under the Bonferonni-corrected threshold $P = 3.55 \times 10^{-5}$, corrected for 1,408 genes simulated using chromosome 1 from the UK Biobank genotype data (see Section S2). Results for gene-$\varepsilon$ are shown with LASSO, Elastic Net (EN), and Ridge Regression (RR) regularizations. We also show the power of gene-$\varepsilon$ without regularization to illustrate the importance of this step (OLS). Additionally, we compare the performance gene-$\varepsilon$ with five existing methods: PEGASUS [6], VEGAS [7], RSS [10], SKAT [11], and MAGMA [15]. The last is a Bayesian method and is evaluated based on the "median probability criterion" (i.e., posterior enrichment probability of a gene is greater than 0.5). All results are based on 100 replicates and standard deviations of the estimates across runs are given in the parentheses. Approaches with the greatest power are bolded in purple, while methods with the lowest FDR is bolded in blue.

| | | *gene-ε Approaches* | | | |
|---|---|---|---|---|---|
| **Causal Genes** | **Metric** | **OLS** | **RR** | **EN** | **LASSO** |
| 1% | Power | 0.686 (0.130) | 0.497 (0.120) | 0.404 (0.125) | 0.406 (0.130) |
| | FDR | 0.512 (0.146) | 0.356 (0.207) | 0.012 (0.055) | **0.005 (0.036)** |
| 10% | Power | 0.131 (0.031) | **0.155 (0.045)** | 0.018 (0.013) | 0.021 (0.016) |
| | FDR | 0.169 (0.082) | 0.678 (0.150) | 0.255 (0.281) | 0.271 (0.306) |

| | | *Other Methods* | | | | |
|---|---|---|---|---|---|---|
| **Causal Genes** | **Metric** | **PEGASUS** | **VEGAS** | **RSS** | **SKAT** | **MAGMA** |
| 1% | Power | 0.678 (0.119) | **0.742 (0.123)** | 0.609 (0.119) | 0.569 (0.130) | 0.737 (0.123) |
| | FDR | 0.527 (0.143) | 0.536 (0.131) | 0.088 (0.102) | 0.487 (0.153) | 0.571 (0.126) |
| 10% | Power | 0.139 (0.027) | 0.106 (0.026) | 0.060 (0.017) | 0.105 (0.021) | 0.152 (0.027) |
| | FDR | 0.169 (0.080) | 0.232 (0.098) | **0.065 (0.089)** | 0.172 (0.088) | 0.191 (0.075) |

**Table S2. Empirical power and false discovery rates (FDR) for detecting enriched genes (genes containing at least one causal SNP) after correcting for multiple hypothesis testing in simulations ($N = 10{,}000$; $h^2 = 0.2$).** We computed standard GWA SNP-level effect sizes (estimated using ordinary least squares) as input to each method listed. We show the power of gene-$\varepsilon$ to identify enriched genes under the Bonferonni-corrected threshold $P = 3.55 \times 10^{-5}$, corrected for 1,408 genes simulated using chromosome 1 from the UK Biobank genotype data (see Section S2). Results for gene-$\varepsilon$ are shown with LASSO, Elastic Net (EN), and Ridge Regression (RR) regularizations. We also show the power of gene-$\varepsilon$ without regularization to illustrate the importance of this step (OLS). Additionally, we compare the performance gene-$\varepsilon$ with five existing methods: PEGASUS [6], VEGAS [7], RSS [10], SKAT [11], and MAGMA [15]. The last is a Bayesian method and is evaluated based on the "median probability criterion" (i.e., posterior enrichment probability of a gene is greater than 0.5). All results are based on 100 replicates and standard deviations of the estimates across runs are given in the parentheses. Approaches with the greatest power are bolded in purple, while methods with the lowest FDR is bolded in blue.

| | | *gene-ε Approaches* | | | |
|---|---|---|---|---|---|
| **Causal Genes** | **Metric** | **OLS** | **RR** | **EN** | **LASSO** |
| 1% | Power | 0.730 (0.132) | 0.555 (0.123) | 0.692 (0.137) | 0.695 (0.126) |
| | FDR | 0.610 (0.106) | 0.237 (0.142) | **0.015 (0.039)** | 0.019 (0.045) |
| 10% | Power | 0.207 (0.035) | 0.118 (0.025) | 0.054 (0.026) | 0.057 (0.031) |
| | FDR | 0.220 (0.083) | 0.209 (0.213) | **0.017 (0.047)** | 0.031 (0.072) |

| | | *Other Methods* | | | | |
|---|---|---|---|---|---|---|
| **Causal Genes** | **Metric** | **PEGASUS** | **VEGAS** | **RSS** | **SKAT** | **MAGMA** |
| 1% | Power | 0.733 (0.130) | **0.780 (0.127)** | 0.663 (0.126) | 0.638 (0.149) | 0.778 (0.131) |
| | FDR | 0.619 (0.105) | 0.632 (0.103) | 0.101 (0.096) | 0.585 (0.118) | 0.664 (0.097) |
| 10% | Power | 0.222 (0.032) | 0.197 (0.030) | 0.117 (0.019) | 0.182 (0.028) | **0.260 (0.035)** |
| | FDR | 0.232 (0.079) | 0.274 (0.083) | 0.076 (0.068) | 0.208 (0.084) | 0.256 (0.081) |

**Table S3. Empirical power and false discovery rates (FDR) for detecting enriched genes (genes containing at least one causal SNP) after correcting for multiple hypothesis testing in simulations ($N = 5{,}000$; $h^2 = 0.6$).** We computed standard GWA SNP-level effect sizes (estimated using ordinary least squares) as input to each method listed. We show the power of gene-$\varepsilon$ to identify enriched genes under the Bonferonni-corrected threshold $P = 3.55 \times 10^{-5}$, corrected for 1,408 genes simulated using chromosome 1 from the UK Biobank genotype data (see Section S2). Results for gene-$\varepsilon$ are shown with LASSO, Elastic Net (EN), and Ridge Regression (RR) regularizations. We also show the power of gene-$\varepsilon$ without regularization to illustrate the importance of this step (OLS). Additionally, we compare the performance gene-$\varepsilon$ with five existing methods: PEGASUS [6], VEGAS [7], RSS [10], SKAT [11], and MAGMA [15]. The last is a Bayesian method and is evaluated based on the "median probability criterion" (i.e., posterior enrichment probability of a gene is greater than 0.5). All results are based on 100 replicates and standard deviations of the estimates across runs are given in the parentheses. Approaches with the greatest power are bolded in purple, while methods with the lowest FDR is bolded in blue.

| Causal Genes | Metric | *gene-ε Approaches* OLS | RR | EN | LASSO |
|---|---|---|---|---|---|
| 1% | Power | 0.798 (0.108) | 0.702 (0.112) | 0.764 (0.106) | 0.765 (0.112) |
| | FDR | 0.703 (0.086) | 0.318 (0.120) | 0.033 (0.062) | **0.033 (0.057)** |
| 10% | Power | 0.355 (0.068) | 0.206 (0.036) | 0.155 (0.045) | 0.164 (0.046) |
| | FDR | 0.265 (0.073) | 0.031 (0.028) | **0.010 (0.022)** | 0.017 (0.027) |

| Causal Genes | Metric | *Other Methods* PEGASUS | VEGAS | RSS | SKAT | MAGMA |
|---|---|---|---|---|---|---|
| 1% | Power | 0.817 (0.104) | **0.876 (0.092)** | 0.773 (0.109) | 0.739 (0.124) | 0.867 (0.093) |
| | FDR | 0.715 (0.089) | 0.717 (0.084) | 0.120 (0.104) | 0.680 (0.092) | 0.746 (0.074) |
| 10% | Power | 0.412 (0.043) | 0.405 (0.041) | 0.250 (0.029) | 0.330 (0.039) | **0.472 (0.045)** |
| | FDR | 0.308 (0.056) | 0.333 (0.055) | 0.068 (0.038) | 0.279 (0.060) | 0.343 (0.053) |

**Table S4. Empirical power and false discovery rates (FDR) for detecting enriched genes (genes containing at least one causal SNP) after correcting for multiple hypothesis testing in simulations ($N$ = 10,000; $h^2 = 0.6$).** We computed standard GWA SNP-level effect sizes (estimated using ordinary least squares) as input to each method listed. We show the power of gene-$\varepsilon$ to identify enriched genes under the Bonferonni-corrected threshold $P = 3.55 \times 10^{-5}$, corrected for 1,408 genes simulated using chromosome 1 from the UK Biobank genotype data (see Section S2). Results for gene-$\varepsilon$ are shown with LASSO, Elastic Net (EN), and Ridge Regression (RR) regularizations. We also show the power of gene-$\varepsilon$ without regularization to illustrate the importance of this step (OLS). Additionally, we compare the performance gene-$\varepsilon$ with five existing methods: PEGASUS [6], VEGAS [7], RSS [10], SKAT [11], and MAGMA [15]. The last is a Bayesian method and is evaluated based on the "median probability criterion" (i.e., posterior enrichment probability of a gene is greater than 0.5). All results are based on 100 replicates and standard deviations of the estimates across runs are given in the parentheses. Approaches with the greatest power are bolded in purple, while methods with the lowest FDR is bolded in blue.

| | | *gene-ε Approaches* | | | |
|---|---|---|---|---|---|
| **Causal Genes** | **Metric** | **OLS** | **RR** | **EN** | **LASSO** |
| 1% | Power | 0.524 (0.136) | 0.340 (0.134) | 0.324 (0.119) | 0.339 (0.127) |
| | FDR | 0.773 (0.102) | 0.467 (0.325) | **0.004 (0.032)** | 0.014 (0.058) |
| 10% | Power | 0.044 (0.024) | 0.053 (0.017) | 0.013 (0.019) | 0.011 (0.015) |
| | FDR | 0.715 (0.136) | 0.560 (0.179) | **0.434 (0.388)** | 0.463 (0.395) |

| | | *Other Methods* | | | | |
|---|---|---|---|---|---|---|
| **Causal Genes** | **Metric** | **PEGASUS** | **VEGAS** | **RSS** | **SKAT** | **MAGMA** |
| 1% | Power | 0.558 (0.126) | 0.611 (0.134) | 0.490 (0.137) | 0.467 (0.132) | **0.615 (0.140)** |
| | FDR | 0.780 (0.096) | 0.778 (0.090) | 0.334 (0.147) | 0.783 (0.092) | 0.789 (0.091) |
| 10% | Power | **0.064 (0.023)** | 0.044 (0.019) | 0.014 (0.010) | 0.051 (0.017) | 0.063 (0.021) |
| | FDR | 0.661 (0.129) | 0.749 (0.114) | 0.563 (0.257) | 0.679 (0.119) | 0.683 (0.135) |

**Table S5. Empirical power and false discovery rates (FDR) for detecting enriched genes (genes containing at least one causal SNP) after correcting for multiple hypothesis testing in simulations with population stratification ($N$ = 5,000; $h^2 = 0.2$).** In this simulation, traits were generated while using the top five principal components (PCs) of the genotype matrix as covariates. GWA summary statistics were computed by fitting a single-SNP univariate linear model (via ordinary least squares) without any control for the additional structure. We show the power of gene-$\varepsilon$ to identify enriched genes under the Bonferonni-corrected threshold $P = 3.55 \times 10^{-5}$, corrected for 1,408 genes simulated using chromosome 1 from the UK Biobank genotype data (see Section S2). Results for gene-$\varepsilon$ are shown with LASSO, Elastic Net (EN), and Ridge Regression (RR) regularizations. We also show the power of gene-$\varepsilon$ without regularization to illustrate the importance of this step (OLS). Additionally, we compare the performance gene-$\varepsilon$ with five existing methods: PEGASUS [6], VEGAS [7], RSS [10], SKAT [11], and MAGMA [15]. The last is a Bayesian method and is evaluated based on the "median probability criterion" (i.e., posterior enrichment probability of a gene is greater than 0.5). All results are based on 100 replicates and standard deviations of the estimates across runs are given in the parentheses. Approaches with the greatest power are bolded in purple, while methods with the lowest FDR is bolded in blue.

| | | *gene-ε Approaches* | | | |
|---|---|---|---|---|---|
| **Causal Genes** | **Metric** | **OLS** | **RR** | **EN** | **LASSO** |
| 1% | Power | 0.658 (0.134) | 0.506 (0.123) | 0.471 (0.130) | 0.487 (0.136) |
| | FDR | 0.763 (0.093) | 0.233 (0.189) | **0.005 (0.031)** | 0.020 (0.060) |
| 10% | Power | 0.086 (0.050) | 0.120 (0.038) | 0.024 (0.021) | 0.021 (0.024) |
| | FDR | 0.618 (0.144) | 0.617 (0.188) | **0.236 (0.269)** | 0.253 (0.311) |

| | | *Other Methods* | | | | |
|---|---|---|---|---|---|---|
| **Causal Genes** | **Metric** | **PEGASUS** | **VEGAS** | **RSS** | **SKAT** | **MAGMA** |
| 1% | Power | 0.694 (0.116) | 0.756 (0.121) | 0.611 (0.119) | 0.583 (0.120) | **0.760 (0.117)** |
| | FDR | 0.793 (0.072) | 0.792 (0.068) | 0.325 (0.116) | 0.775 (0.084) | 0.807 (0.064) |
| 10% | Power | **0.154 (0.029)** | 0.126 (0.029) | 0.061 (0.017) | 0.121 (0.024) | 0.176 (0.031) |
| | FDR | 0.536 (0.117) | 0.605 (0.104) | 0.273 (0.109) | 0.534 (0.122) | 0.549 (0.102) |

**Table S6. Empirical power and false discovery rates (FDR) for detecting enriched genes (genes containing at least one causal SNP) after correcting for multiple hypothesis testing in simulations with population stratification ($N$ = 10,000; $h^2 = 0.2$).** In this simulation, traits were generated while using the top five principal components (PCs) of the genotype matrix as covariates. GWA summary statistics were computed by fitting a single-SNP univariate linear model (via ordinary least squares) without any control for the additional structure. We show the power of gene-$\varepsilon$ to identify enriched genes under the Bonferonni-corrected threshold $P = 3.55 \times 10^{-5}$, corrected for 1,408 genes simulated using chromosome 1 from the UK Biobank genotype data (see Section S2). Results for gene-$\varepsilon$ are shown with LASSO, Elastic Net (EN), and Ridge Regression (RR) regularizations. We also show the power of gene-$\varepsilon$ without regularization to illustrate the importance of this step (OLS). Additionally, we compare the performance gene-$\varepsilon$ with five existing methods: PEGASUS [6], VEGAS [7], RSS [10], SKAT [11], and MAGMA [15]. The last is a Bayesian method and is evaluated based on the "median probability criterion" (i.e., posterior enrichment probability of a gene is greater than 0.5). All results are based on 100 replicates and standard deviations of the estimates across runs are given in the parentheses. Approaches with the greatest power are bolded in purple, while methods with the lowest FDR is bolded in blue.

| | | *gene-ε Approaches* | | | |
|---|---|---|---|---|---|
| **Causal Genes** | **Metric** | **OLS** | **RR** | **EN** | **LASSO** |
| 1% | Power | 0.727 (0.126) | 0.553 (0.130) | 0.740 (0.128) | 0.734 (0.130) |
| | FDR | 0.760 (0.086) | 0.274 (0.126) | **0.031 (0.056)** | 0.050 (0.076) |
| 10% | Power | 0.155 (0.059) | 0.106 (0.022) | 0.083 (0.036) | 0.080 (0.036) |
| | FDR | 0.483 (0.134) | 0.042 (0.085) | **0.020 (0.047)** | 0.032 (0.054) |

| | | *Other Methods* | | | | |
|---|---|---|---|---|---|---|
| **Causal Genes** | **Metric** | **PEGASUS** | **VEGAS** | **RSS** | **SKAT** | **MAGMA** |
| 1% | Power | 0.734 (0.126) | 0.786 (0.122) | 0.662 (0.129) | 0.645 (0.134) | **0.796 (0.119)** |
| | FDR | 0.785 (0.075) | 0.787 (0.070) | 0.267 (0.123) | 0.776 (0.071) | 0.801 (0.064) |
| 10% | Power | 0.237 (0.036) | 0.210 (0.032) | 0.115 (0.019) | 0.189 (0.030) | **0.275 (0.038)** |
| | FDR | 0.462 (0.111) | 0.505 (0.102) | 0.187 (0.078) | 0.464 (0.104) | 0.467 (0.102) |

**Table S7. Empirical power and false discovery rates (FDR) for detecting enriched genes (genes containing at least one causal SNP) after correcting for multiple hypothesis testing in simulations with population stratification ($N$ = 5,000; $h^2 = 0.6$).** In this simulation, traits were generated while using the top five principal components (PCs) of the genotype matrix as covariates. GWA summary statistics were computed by fitting a single-SNP univariate linear model (via ordinary least squares) without any control for the additional structure. We show the power of gene-$\varepsilon$ to identify enriched genes under the Bonferonni-corrected threshold $P = 3.55 \times 10^{-5}$, corrected for 1,408 genes simulated using chromosome 1 from the UK Biobank genotype data (see Section S2). Results for gene-$\varepsilon$ are shown with LASSO, Elastic Net (EN), and Ridge Regression (RR) regularizations. We also show the power of gene-$\varepsilon$ without regularization to illustrate the importance of this step (OLS). Additionally, we compare the performance gene-$\varepsilon$ with five existing methods: PEGASUS [6], VEGAS [7], RSS [10], SKAT [11], and MAGMA [15]. The last is a Bayesian method and is evaluated based on the "median probability criterion" (i.e., posterior enrichment probability of a gene is greater than 0.5). All results are based on 100 replicates and standard deviations of the estimates across runs are given in the parentheses. Approaches with the greatest power are bolded in purple, while methods with the lowest FDR is bolded in blue.

| | | *gene-ε Approaches* | | | |
|---|---|---|---|---|---|
| **Causal Genes** | **Metric** | **OLS** | **RR** | **EN** | **LASSO** |
| 1% | Power | 0.798 (0.100) | 0.712 (0.111) | 0.793 (0.108) | 0.774 (0.118) |
| | FDR | 0.789 (0.077) | 0.345 (0.119) | **0.077 (0.083)** | 0.101 (0.100) |
| 10% | Power | 0.168 (0.125) | 0.211 (0.031) | 0.261 (0.062) | 0.261 (0.061) |
| | FDR | 0.490 (0.144) | **0.011 (0.019)** | 0.021 (0.026) | 0.027 (0.029) |

| | | *Other Methods* | | | | |
|---|---|---|---|---|---|---|
| **Causal Genes** | **Metric** | **PEGASUS** | **VEGAS** | **RSS** | **SKAT** | **MAGMA** |
| 1% | Power | 0.809 (0.104) | 0.869 (0.090) | 0.773 (0.113) | 0.744 (0.123) | **0.870 (0.089)** |
| | FDR | 0.833 (0.046) | 0.832 (0.047) | 0.301 (0.098) | 0.806 (0.059) | 0.848 (0.040) |
| 10% | Power | 0.426 (0.044) | 0.420 (0.042) | 0.251 (0.030) | 0.342 (0.038) | **0.488 (0.048)** |
| | FDR | 0.449 (0.068) | 0.470 (0.067) | 0.134 (0.044) | 0.429 (0.075) | 0.474 (0.064) |

**Table S8. Empirical power and false discovery rates (FDR) for detecting enriched genes (genes containing at least one causal SNP) after correcting for multiple hypothesis testing in simulations with population stratification ($N$ = 10,000; $h^2 = 0.6$).** In this simulation, traits were generated while using the top five principal components (PCs) of the genotype matrix as covariates. GWA summary statistics were computed by fitting a single-SNP univariate linear model (via ordinary least squares) without any control for the additional structure. We show the power of gene-$\varepsilon$ to identify enriched genes under the Bonferonni-corrected threshold $P = 3.55 \times 10^{-5}$, corrected for 1,408 genes simulated using chromosome 1 from the UK Biobank genotype data (see Section S2). Results for gene-$\varepsilon$ are shown with LASSO, Elastic Net (EN), and Ridge Regression (RR) regularizations. We also show the power of gene-$\varepsilon$ without regularization to illustrate the importance of this step (OLS). Additionally, we compare the performance gene-$\varepsilon$ with five existing methods: PEGASUS [6], VEGAS [7], RSS [10], SKAT [11], and MAGMA [15]. The last is a Bayesian method and is evaluated based on the "median probability criterion" (i.e., posterior enrichment probability of a gene is greater than 0.5). All results are based on 100 replicates and standard deviations of the estimates across runs are given in the parentheses. Approaches with the greatest power are bolded in purple, while methods with the lowest FDR is bolded in blue.

| | | *gene-ε Approaches* | | | |
|---|---|---|---|---|---|
| **Causal Genes** | **Metric** | **OLS** | **RR** | **EN** | **LASSO** |
| 1% | Power | 0.487 (0.118) | 0.482 (0.167) | 0.126 (0.063) | 0.136 (0.063) |
| | FDR | 0.407 (0.178) | 0.770 (0.259) | **0.000 (0.000)** | 0.006 (0.054) |
| 10% | Power | 0.019 (0.029) | **0.070 (0.055)** | 0.005 (0.005) | 0.006 (0.005) |
| | FDR | 0.077 (0.185) | 0.622 (0.208) | 0.581 (0.352) | 0.447 (0.334) |

| | | *Other Methods* | | | | |
|---|---|---|---|---|---|---|
| **Causal Genes** | **Metric** | **PEGASUS** | **VEGAS** | **RSS** | **SKAT** | **MAGMA** |
| 1% | Power | 0.457 (0.110) | **0.563 (0.104)** | 0.540 (0.110) | 0.388 (0.106) | 0.514 (0.124) |
| | FDR | 0.366 (0.192) | 0.393 (0.177) | 0.090 (0.124) | 0.329 (0.212) | 0.430 (0.183) |
| 10% | Power | 0.014 (0.007) | 0.008 (0.004) | 0.006 (0.002) | 0.012 (0.006) | 0.010 (0.006) |
| | FDR | 0.060 (0.158) | 0.178 (0.290) | **0.016 (0.088)** | 0.073 (0.177) | 0.121 (0.237) |

**Table S9. Empirical power and false discovery rates (FDR) for detecting enriched genes (genes containing at least one causal SNP) after correcting for multiple hypothesis testing in simulations with gene boundaries augmented by a 50 kilobase (kb) buffer ($N$ = 5,000; $h^2 = 0.2$).** We computed standard GWA SNP-level effect sizes (estimated using ordinary least squares) as input to each method listed. We show the power of gene-$\varepsilon$ to identify enriched genes under the Bonferonni-corrected threshold $P = 2.61 \times 10^{-5}$, corrected for 1,916 genes simulated using chromosome 1 from the UK Biobank genotype data (see Section S2). Results for gene-$\varepsilon$ are shown with LASSO, Elastic Net (EN), and Ridge Regression (RR) regularizations. We also show the power of gene-$\varepsilon$ without regularization to illustrate the importance of this step (OLS). Additionally, we compare the performance gene-$\varepsilon$ with five existing methods: PEGASUS [6], VEGAS [7], RSS [10], SKAT [11], and MAGMA [15]. The last is a Bayesian method and is evaluated based on the "median probability criterion" (i.e., posterior enrichment probability of a gene is greater than 0.5). All results are based on 100 replicates and standard deviations of the estimates across runs are given in the parentheses. Approaches with the greatest power are bolded in purple, while methods with the lowest FDR is bolded in blue.

| | | *gene-ε Approaches* | | | |
|---|---|---|---|---|---|
| **Causal Genes** | **Metric** | **OLS** | **RR** | **EN** | **LASSO** |
| 1% | Power | 0.602 (0.115) | 0.388 (0.119) | 0.220 (0.086) | 0.226 (0.088) |
| | FDR | 0.523 (0.151) | 0.229 (0.230) | **0.000 (0.000)** | **0.000 (0.000)** |
| 10% | Power | 0.106 (0.055) | **0.221 (0.057)** | 0.011 (0.019) | 0.004 (0.004) |
| | FDR | 0.183 (0.131) | 0.746 (0.094) | 0.374 (0.398) | 0.396 (0.494) |

| | | *Other Methods* | | | | |
|---|---|---|---|---|---|---|
| **Causal Genes** | **Metric** | **PEGASUS** | **VEGAS** | **RSS** | **SKAT** | **MAGMA** |
| 1% | Power | 0.612 (0.113) | **0.700 (0.121)** | 0.654 (0.120) | 0.550 (0.127) | 0.652 (0.117) |
| | FDR | 0.533 (0.146) | 0.547 (0.154) | 0.115 (0.137) | 0.508 (0.156) | 0.601 (0.139) |
| 10% | Power | 0.057 (0.013) | 0.024 (0.010) | 0.014 (0.007) | 0.044 (0.013) | 0.049 (0.012) |
| | FDR | 0.112 (0.114) | 0.174 (0.199) | 0.107 (0.200) | **0.097 (0.119)** | 0.124 (0.131) |

**Table S10. Empirical power and false discovery rates (FDR) for detecting enriched genes (genes containing at least one causal SNP) after correcting for multiple hypothesis testing in simulations with gene boundaries augmented by a 50 kilobase (kb) buffer ($N$ = 10,000; $h^2 = 0.2$).** We computed standard GWA SNP-level effect sizes (estimated using ordinary least squares) as input to each method listed. We show the power of gene-$\varepsilon$ to identify enriched genes under the Bonferonni-corrected threshold $P = 2.61 \times 10^{-5}$, corrected for 1,916 genes simulated using chromosome 1 from the UK Biobank genotype data (see Section S2). Results for gene-$\varepsilon$ are shown with LASSO, Elastic Net (EN), and Ridge Regression (RR) regularizations. We also show the power of gene-$\varepsilon$ without regularization to illustrate the importance of this step (OLS). Additionally, we compare the performance gene-$\varepsilon$ with five existing methods: PEGASUS [6], VEGAS [7], RSS [10], SKAT [11], and MAGMA [15]. The last is a Bayesian method and is evaluated based on the "median probability criterion" (i.e., posterior enrichment probability of a gene is greater than 0.5). All results are based on 100 replicates and standard deviations of the estimates across runs are given in the parentheses. Approaches with the greatest power are bolded in purple, while methods with the lowest FDR is bolded in blue.

| | | *gene-ε Approaches* | | | |
|---|---|---|---|---|---|
| **Causal Genes** | **Metric** | **OLS** | **RR** | **EN** | **LASSO** |
| 1% | Power | 0.707 (0.111) | 0.567 (0.107) | 0.624 (0.115) | 0.638 (0.123) |
| | FDR | 0.591 (0.132) | 0.260 (0.161) | **0.004 (0.022)** | 0.006 (0.030) |
| 10% | Power | 0.101 (0.030) | **0.172 (0.060)** | 0.013 (0.012) | 0.010 (0.007) |
| | FDR | 0.119 (0.101) | 0.365 (0.160) | 0.060 (0.192) | **0.056 (0.183)** |

| | | *Other Methods* | | | | |
|---|---|---|---|---|---|---|
| **Causal Genes** | **Metric** | **PEGASUS** | **VEGAS** | **RSS** | **SKAT** | **MAGMA** |
| 1% | Power | 0.719 (0.108) | **0.787 (0.106)** | 0.744 (0.107) | 0.636 (0.114) | 0.753 (0.109) |
| | FDR | 0.605 (0.127) | 0.626 (0.116) | 0.116 (0.126) | 0.575 (0.142) | 0.677 (0.108) |
| 10% | Power | 0.117 (0.019) | 0.065 (0.015) | 0.054 (0.012) | 0.093 (0.016) | 0.118 (0.017) |
| | FDR | 0.132 (0.101) | 0.226 (0.154) | 0.091 (0.128) | 0.116 (0.106) | 0.162 (0.106) |

**Table S11. Empirical power and false discovery rates (FDR) for detecting enriched genes (genes containing at least one causal SNP) after correcting for multiple hypothesis testing in simulations with gene boundaries augmented by a 50 kilobase (kb) buffer ($N$ = 5,000; $h^2 = 0.6$).** We computed standard GWA SNP-level effect sizes (estimated using ordinary least squares) as input to each method listed. We show the power of gene-$\varepsilon$ to identify enriched genes under the Bonferonni-corrected threshold $P = 2.61 \times 10^{-5}$, corrected for 1,916 genes simulated using chromosome 1 from the UK Biobank genotype data (see Section S2). Results for gene-$\varepsilon$ are shown with LASSO, Elastic Net (EN), and Ridge Regression (RR) regularizations. We also show the power of gene-$\varepsilon$ without regularization to illustrate the importance of this step (OLS). Additionally, we compare the performance gene-$\varepsilon$ with five existing methods: PEGASUS [6], VEGAS [7], RSS [10], SKAT [11], and MAGMA [15]. The last is a Bayesian method and is evaluated based on the "median probability criterion" (i.e., posterior enrichment probability of a gene is greater than 0.5). All results are based on 100 replicates and standard deviations of the estimates across runs are given in the parentheses. Approaches with the greatest power are bolded in purple, while methods with the lowest FDR is bolded in blue.

| | | *gene-ε Approaches* | | | |
|---|---|---|---|---|---|
| **Causal Genes** | **Metric** | **OLS** | **RR** | **EN** | **LASSO** |
| 1% | Power | 0.784 (0.103) | 0.617 (0.114) | 0.667 (0.115) | 0.673 (0.118) |
| | FDR | 0.691 (0.096) | 0.219 (0.157) | **0.006 (0.023)** | 0.007 (0.028) |
| 10% | Power | 0.224 (0.048) | 0.100 (0.083) | 0.024 (0.015) | 0.020 (0.012) |
| | FDR | 0.199 (0.088) | 0.053 (0.134) | **0.007 (0.040)** | 0.008 (0.041) |

| | | *Other Methods* | | | | |
|---|---|---|---|---|---|---|
| **Causal Genes** | **Metric** | **PEGASUS** | **VEGAS** | **RSS** | **SKAT** | **MAGMA** |
| 1% | Power | 0.799 (0.099) | **0.843 (0.087)** | 0.820 (0.087) | 0.733 (0.104) | 0.826 (0.088) |
| | FDR | 0.705 (0.095) | 0.726 (0.078) | 0.118 (0.113) | 0.680 (0.098) | 0.766 (0.071) |
| 10% | Power | 0.261 (0.027) | 0.213 (0.026) | 0.159 (0.018) | 0.217 (0.026) | **0.304 (0.028)** |
| | FDR | 0.229 (0.082) | 0.278 (0.094) | 0.089 (0.066) | 0.204 (0.082) | 0.260 (0.081) |

**Table S12. Empirical power and false discovery rates (FDR) for detecting enriched genes (genes containing at least one causal SNP) after correcting for multiple hypothesis testing in simulations with gene boundaries augmented by a 50 kilobase (kb) buffer ($N$ = 10,000; $h^2 = 0.6$).** We computed standard GWA SNP-level effect sizes (estimated using ordinary least squares) as input to each method listed. We show the power of gene-$\varepsilon$ to identify enriched genes under the Bonferonni-corrected threshold $P = 2.61 \times 10^{-5}$, corrected for 1,916 genes simulated using chromosome 1 from the UK Biobank genotype data (see Section S2). Results for gene-$\varepsilon$ are shown with LASSO, Elastic Net (EN), and Ridge Regression (RR) regularizations. We also show the power of gene-$\varepsilon$ without regularization to illustrate the importance of this step (OLS). Additionally, we compare the performance gene-$\varepsilon$ with five existing methods: PEGASUS [6], VEGAS [7], RSS [10], SKAT [11], and MAGMA [15]. The last is a Bayesian method and is evaluated based on the "median probability criterion" (i.e., posterior enrichment probability of a gene is greater than 0.5). All results are based on 100 replicates and standard deviations of the estimates across runs are given in the parentheses. Approaches with the greatest power are bolded in purple, while methods with the lowest FDR is bolded in blue.

| | | *gene-ε Approaches* | | | |
|---|---|---|---|---|---|
| **Causal Genes** | **Metric** | **OLS** | **RR** | **EN** | **LASSO** |
| 1% | Power | 0.431 (0.117) | 0.427 (0.137) | 0.155 (0.073) | 0.158 (0.073) |
| | FDR | 0.838 (0.128) | 0.577 (0.286) | **0.004 (0.035)** | 0.014 (0.081) |
| 10% | Power | 0.026 (0.089) | **0.079 (0.030)** | 0.027 (0.035) | 0.009 (0.021) |
| | FDR | 0.903 (0.091) | **0.571 (0.125)** | 0.624 (0.392) | 0.705 (0.430) |

| | | *Other Methods* | | | | |
|---|---|---|---|---|---|---|
| **Causal Genes** | **Metric** | **PEGASUS** | **VEGAS** | **RSS** | **SKAT** | **MAGMA** |
| 1% | Power | 0.463 (0.116) | **0.556 (0.117)** | 0.520 (0.110) | 0.387 (0.110) | 0.500 (0.110) |
| | FDR | 0.847 (0.115) | 0.828 (0.114) | 0.608 (0.133) | 0.858 (0.115) | 0.846 (0.105) |
| 10% | Power | 0.027 (0.013) | 0.016 (0.011) | 0.006 (0.006) | 0.022 (0.012) | 0.020 (0.011) |
| | FDR | 0.868 (0.094) | 0.921 (0.081) | 0.898 (0.103) | 0.886 (0.091) | 0.896 (0.091) |

**Table S13. Empirical power and false discovery rates (FDR) for detecting enriched genes (genes containing at least one causal SNP) after correcting for multiple hypothesis testing in simulations with gene boundaries augmented by a 50 kilobase (kb) buffer and with population stratification ($N$ = 5,000; $h^2$ = 0.2).** In this simulation, traits were generated while using the top five principal components (PCs) of the genotype matrix as covariates. GWA summary statistics were computed by fitting a single-SNP univariate linear model (via ordinary least squares) without any control for the additional structure. We show the power of gene-$\varepsilon$ to identify enriched genes under the Bonferonni-corrected threshold $P = 2.61 \times 10^{-5}$, corrected for 1,916 genes simulated using chromosome 1 from the UK Biobank genotype data (see Section S2). Results for gene-$\varepsilon$ are shown with LASSO, Elastic Net (EN), and Ridge Regression (RR) regularizations. We also show the power of gene-$\varepsilon$ without regularization to illustrate the importance of this step (OLS). Additionally, we compare the performance gene-$\varepsilon$ with five existing methods: PEGASUS [6], VEGAS [7], RSS [10], SKAT [11], and MAGMA [15]. The last is a Bayesian method and is evaluated based on the "median probability criterion" (i.e., posterior enrichment probability of a gene is greater than 0.5). All results are based on 100 replicates and standard deviations of the estimates across runs are given in the parentheses. Approaches with the greatest power are bolded in purple, while methods with the lowest FDR is bolded in blue.

| | | *gene-ε Approaches* | | | |
|---|---|---|---|---|---|
| **Causal Genes** | **Metric** | **OLS** | **RR** | **EN** | **LASSO** |
| 1% | Power | 0.572 (0.131) | 0.426 (0.121) | 0.294 (0.101) | 0.298 (0.099) |
| | FDR | 0.833 (0.085) | 0.311 (0.255) | **0.000 (0.000)** | **0.000 (0.000)** |
| 10% | Power | 0.036 (0.017) | **0.198 (0.085)** | 0.018 (0.035) | 0.016 (0.052) |
| | FDR | 0.816 (0.131) | 0.645 (0.202) | **0.466 (0.469)** | 0.799 (0.391) |

| | | *Other Methods* | | | | |
|---|---|---|---|---|---|---|
| **Causal Genes** | **Metric** | **PEGASUS** | **VEGAS** | **RSS** | **SKAT** | **MAGMA** |
| 1% | Power | 0.619 (0.125) | **0.699 (0.114)** | 0.649 (0.118) | 0.554 (0.128) | 0.662 (0.123) |
| | FDR | 0.849 (0.078) | 0.837 (0.077) | 0.590 (0.105) | 0.835 (0.093) | 0.857 (0.070) |
| 10% | Power | 0.076 (0.017) | 0.041 (0.015) | 0.019 (0.010) | 0.059 (0.016) | 0.071 (0.017) |
| | FDR | 0.739 (0.135) | 0.836 (0.092) | 0.775 (0.127) | 0.747 (0.141) | 0.765 (0.115) |

**Table S14. Empirical power and false discovery rates (FDR) for detecting enriched genes (genes containing at least one causal SNP) after correcting for multiple hypothesis testing in simulations with gene boundaries augmented by a 50 kilobase (kb) buffer and with population stratification ($N$ = 10,000; $h^2 = 0.2$).** In this simulation, traits were generated while using the top five principal components (PCs) of the genotype matrix as covariates. GWA summary statistics were computed by fitting a single-SNP univariate linear model (via ordinary least squares) without any control for the additional structure. We show the power of gene-$\varepsilon$ to identify enriched genes under the Bonferonni-corrected threshold $P = 2.61 \times 10^{-5}$, corrected for 1,916 genes simulated using chromosome 1 from the UK Biobank genotype data (see Section S2). Results for gene-$\varepsilon$ are shown with LASSO, Elastic Net (EN), and Ridge Regression (RR) regularizations. We also show the power of gene-$\varepsilon$ without regularization to illustrate the importance of this step (OLS). Additionally, we compare the performance gene-$\varepsilon$ with five existing methods: PEGASUS [6], VEGAS [7], RSS [10], SKAT [11], and MAGMA [15]. The last is a Bayesian method and is evaluated based on the "median probability criterion" (i.e., posterior enrichment probability of a gene is greater than 0.5). All results are based on 100 replicates and standard deviations of the estimates across runs are given in the parentheses. Approaches with the greatest power are bolded in purple, while methods with the lowest FDR is bolded in blue.

| | | *gene-ε Approaches* | | | |
|---|---|---|---|---|---|
| **Causal Genes** | **Metric** | **OLS** | **RR** | **EN** | **LASSO** |
| 1% | Power | 0.687 (0.119) | 0.636 (0.116) | 0.696 (0.122) | 0.700 (0.114) |
| | FDR | 0.820 (0.097) | 0.324 (0.162) | **0.011 (0.043)** | 0.019 (0.052) |
| 10% | Power | 0.069 (0.033) | **0.155 (0.030)** | 0.008 (0.004) | 0.006 (0.003) |
| | FDR | 0.702 (0.171) | 0.269 (0.112) | **0.042 (0.134)** | 0.207 (0.361) |

| | | *Other Methods* | | | | |
|---|---|---|---|---|---|---|
| **Causal Genes** | **Metric** | **PEGASUS** | **VEGAS** | **RSS** | **SKAT** | **MAGMA** |
| 1% | Power | 0.712 (0.108) | **0.781 (0.101)** | 0.745 (0.113) | 0.649 (0.113) | 0.751 (0.111) |
| | FDR | 0.847 (0.076) | 0.837 (0.074) | 0.513 (0.112) | 0.840 (0.083) | 0.849 (0.069) |
| 10% | Power | 0.133 (0.019) | 0.077 (0.017) | 0.053 (0.013) | 0.109 (0.017) | 0.137 (0.023) |
| | FDR | 0.614 (0.179) | 0.719 (0.149) | 0.530 (0.129) | 0.634 (0.169) | 0.614 (0.161) |

**Table S15. Empirical power and false discovery rates (FDR) for detecting enriched genes (genes containing at least one causal SNP) after correcting for multiple hypothesis testing in simulations with gene boundaries augmented by a 50 kilobase (kb) buffer and with population stratification ($N$ = 5,000; $h^2$ = 0.6).** In this simulation, traits were generated while using the top five principal components (PCs) of the genotype matrix as covariates. GWA summary statistics were computed by fitting a single-SNP univariate linear model (via ordinary least squares) without any control for the additional structure. We show the power of gene-$\varepsilon$ to identify enriched genes under the Bonferonni-corrected threshold $P = 2.61 \times 10^{-5}$, corrected for 1,916 genes simulated using chromosome 1 from the UK Biobank genotype data (see Section S2). Results for gene-$\varepsilon$ are shown with LASSO, Elastic Net (EN), and Ridge Regression (RR) regularizations. We also show the power of gene-$\varepsilon$ without regularization to illustrate the importance of this step (OLS). Additionally, we compare the performance gene-$\varepsilon$ with five existing methods: PEGASUS [6], VEGAS [7], RSS [10], SKAT [11], and MAGMA [15]. The last is a Bayesian method and is evaluated based on the "median probability criterion" (i.e., posterior enrichment probability of a gene is greater than 0.5). All results are based on 100 replicates and standard deviations of the estimates across runs are given in the parentheses. Approaches with the greatest power are bolded in purple, while methods with the lowest FDR is bolded in blue.

| | | *gene-ε Approaches* | | | |
|---|---|---|---|---|---|
| **Causal Genes** | **Metric** | **OLS** | **RR** | **EN** | **LASSO** |
| 1% | Power | 0.762 (0.103) | 0.682 (0.118) | 0.752 (0.119) | 0.764 (0.111) |
| | FDR | 0.828 (0.072) | 0.326 (0.147) | **0.015 (0.041)** | 0.035 (0.060) |
| 10% | Power | 0.055 (0.059) | 0.151 (0.045) | 0.037 (0.020) | 0.033 (0.022) |
| | FDR | 0.729 (0.159) | 0.029 (0.036) | **0.007 (0.032)** | 0.066 (0.178) |

| | | *Other Methods* | | | | |
|---|---|---|---|---|---|---|
| **Causal Genes** | **Metric** | **PEGASUS** | **VEGAS** | **RSS** | **SKAT** | **MAGMA** |
| 1% | Power | 0.802 (0.104) | **0.842 (0.089)** | 0.811 (0.096) | 0.733 (0.111) | 0.825 (0.092) |
| | FDR | 0.865 (0.054) | 0.862 (0.050) | 0.542 (0.100) | 0.854 (0.061) | 0.881 (0.042) |
| 10% | Power | 0.284 (0.029) | 0.228 (0.029) | 0.158 (0.019) | 0.235 (0.026) | **0.324 (0.032)** |
| | FDR | 0.543 (0.100) | 0.589 (0.097) | 0.336 (0.075) | 0.534 (0.113) | 0.546 (0.088) |

**Table S16. Empirical power and false discovery rates (FDR) for detecting enriched genes (genes containing at least one causal SNP) after correcting for multiple hypothesis testing in simulations with gene boundaries augmented by a 50 kilobase (kb) buffer and with population stratification ($N$ = 10,000; $h^2 = 0.6$).** In this simulation, traits were generated while using the top five principal components (PCs) of the genotype matrix as covariates. GWA summary statistics were computed by fitting a single-SNP univariate linear model (via ordinary least squares) without any control for the additional structure. We show the power of gene-$\varepsilon$ to identify enriched genes under the Bonferonni-corrected threshold $P = 2.61 \times 10^{-5}$, corrected for 1,916 genes simulated using chromosome 1 from the UK Biobank genotype data (see Section S2). Results for gene-$\varepsilon$ are shown with LASSO, Elastic Net (EN), and Ridge Regression (RR) regularizations. We also show the power of gene-$\varepsilon$ without regularization to illustrate the importance of this step (OLS). Additionally, we compare the performance gene-$\varepsilon$ with five existing methods: PEGASUS [6], VEGAS [7], RSS [10], SKAT [11], and MAGMA [15]. The last is a Bayesian method and is evaluated based on the "median probability criterion" (i.e., posterior enrichment probability of a gene is greater than 0.5). All results are based on 100 replicates and standard deviations of the estimates across runs are given in the parentheses. Approaches with the greatest power are bolded in purple, while methods with the lowest FDR is bolded in blue.

| | | *Significance Level* | | | |
|---|---|---|---|---|---|
| **Sample Size** | **gene-$\varepsilon$ Approach** | $\alpha = \mathbf{0.05}$ | $\alpha = \mathbf{0.01}$ | $\alpha = \mathbf{0.001}$ | $\alpha = \mathbf{2.61 \times 10^{-5}}$ |
| $N$ = 5,000 | OLS | 0.0481 (0.0103) | 0.0091 (0.0038) | 0.0008 (0.0010) | 0.0000 (0.0001) |
| | Ridge Regression | 0.0082 (0.0024) | 0.0065 (0.0020) | 0.0056 (0.0018) | 0.0000 (0.0003) |
| | Elastic Net | 0.0035 (0.0094) | 0.0013 (0.0045) | 0.0004 (0.0016) | 0.0000 (0.0001) |
| | LASSO | 0.0043 (0.0093) | 0.0015 (0.0043) | 0.0004 (0.0013) | 0.0000 (0.0001) |
| $N$ = 10,000 | OLS | 0.0486 (0.0109) | 0.0095 (0.0034) | 0.0008 (0.0008) | 0.0000 (0.0000) |
| | Ridge Regression | 0.0067 (0.0029) | 0.0050 (0.0031) | 0.0044 (0.0033) | 0.0000 (0.0003) |
| | Elastic Net | 0.0009 (0.0028) | 0.0004 (0.0009) | 0.0000 (0.0002) | 0.0000 (0.0001) |
| | LASSO | 0.0007 (0.0026) | 0.0002 (0.0009) | 0.0000 (0.0002) | 0.0000 (0.0001) |

**Table S17. Empirical type I error estimates using different gene-$\varepsilon$ approaches.** Here, quantitative traits are simulated with just noise randomly drawn from standard normal distributions. This represents the scenario in which all SNPs are non-causal and satisfy the conventional null hypothesis $H_0 : \beta_j = 0$. GWA summary statistics were computed by fitting a single-SNP univariate linear model (via ordinary least squares). Each table entry lists the mean type I error rate estimates for the four gene-$\varepsilon$ modeling approaches — which is computed as the proportion of $P$-values under some significance level $\alpha$. Empirical size for the analyses used significance levels of $\alpha$ = 0.05, 0.01, 0.001, and $2.61 \times 10^{-5}$ (the Bonferonni-corrected threshold), respectively. Sample sizes of the individual-level data (used to derive the summary statistics), were set to $N$ = 5,000 and 10,000 observations. These results are based on 100 simulated datasets and the standard errors across the replicated are included in the parentheses. Overall, gene-$\varepsilon$ controls the type I error rate for reasonably sized datasets, and can be slightly conservative when the sample size is small and the GWA summary statistics are less precise/more inflated.

| gene-$\varepsilon$ Approach | Trait | # Mix. Comp. | % Associated SNPs | % Causal SNPs | $\varepsilon$-genic Threshold ($\sigma_\varepsilon^2 = \sigma_2^2$) |
|---|---|---|---|---|---|
| Elastic Net | Height | 8 | 10.88% | 1.39% | $3.46 \times 10^{-5}$ |
| | BMI | 6 | 12.61% | 6.23% | $5.18 \times 10^{-5}$ |
| | MCV | 8 | 13.38% | 0.32% | $6.15 \times 10^{-5}$ |
| | MPV | 9 | 11.49% | 0.21% | $7.05 \times 10^{-5}$ |
| | PLC | 8 | 13.20% | 0.45% | $6.56 \times 10^{-5}$ |
| | WHR | 6 | 13.33% | 6.28% | $5.01 \times 10^{-5}$ |
| OLS | Height | 4 | 48.00% | 7.90% | $4.16 \times 10^{-5}$ |
| | BMI | 3 | 48.74% | 23.28% | $4.39 \times 10^{-5}$ |
| | MCV | 9 | 35.87% | 1.67% | $6.04 \times 10^{-5}$ |
| | MPV | 9 | 35.94% | 2.21% | $6.70 \times 10^{-5}$ |
| | PLC | 7 | 40.42% | 2.45% | $5.96 \times 10^{-5}$ |
| | WHR | 2 | 99.99% | 44.51% | $1.55 \times 10^{-5}$ |

**Table S18. Characterization of the genetic architectures of six traits assayed in European-ancestry individuals in the UK Biobank.** Here, we report the way difference regularization makes when gene-$\varepsilon$ characterizes $\varepsilon$-genic effects in complex traits. Results are shown for Elastic Net (which is highlighted in the main text). We also show results when no shrinkage is applied to illustrate the importance of this step (denoted by OLS). In the former case, we regress the GWA SNP-level effect size estimates onto chromosome-specific LD matrices to derive a regularized set of summary statistics $\widetilde{\boldsymbol{\beta}}$. gene-$\varepsilon$ assumes a reformulated null distribution of SNP-level effects $\widetilde{\beta}_j \sim \mathcal{N}(0, \sigma_\varepsilon^2)$, where $\sigma_\varepsilon^2$ is the SNP-level null threshold and represents the maximum proportion of phenotypic variance explained (PVE) by a spurious or non-associated SNP. We used an EM-algorithm with 100 iterations to fit $K$-mixture Gaussian models over the regularized effect sizes to estimate $\sigma_\varepsilon^2$. Here, each mixture component had distinctively smaller variances ($\sigma_1^2 > \cdots > \sigma_K^2$; with the $K$-th component fixed at $\sigma_K^2 = 0$), and the number of total mixture components $K$ was chosen based on a grid of values where the best model yielded the highest Bayesian Information Criterion (BIC). We assume associated SNPs appear in the first component, non-associated SNPs appear in the last component, and null SNPs with spurious effects fell in between (i.e., $\sigma_\varepsilon^2 = \sigma_2^2$). Thus, a SNP is considered to have some level of association with a trait if $\mathbb{E}[\beta_j^2] > \sigma_K^2 = 0$; while a SNP is considered "causal" if $\mathbb{E}[\beta_j^2] > \sigma_2^2$. Column 3 gives the $K$ used for each trait. Column 4 and 5 detail the percentage of associated and causal SNPs, respectively. The last column gives the mean threshold for $\varepsilon$-genic effects across the chromosomes.

**Table S19. Significant genes for body height in the UK Biobank analysis using gene-$\varepsilon$-EN.** Here, we analyze 17,680 genes from $N$ = 349,468 individuals of European-ancestry. This file gives the gene-$\varepsilon$ gene-level association $P$-values using Elastic Net regularized effect sizes when gene boundaries are defined by (page 1) using UCSC annotations directly, and (page 2) augmenting the gene boundaries by adding SNPs within a ±50kb buffer. Significance was determined by using a Bonferroni-corrected $P$-value threshold (in our analyses, $P = 0.05/14322$ autosomal genes $= 3.49\times10^{-6}$ and $P = 0.05/17680$ autosomal genes $= 2.83\times10^{-6}$, respectively). The columns of tables on both pages provide: (1) chromosome position; (2) gene name; (3) gene-$\varepsilon$-EN gene $P$-value; (4) gene-specific heritability estimates; (5) whether or not an association between gene and trait is listed in the GWAS catalog (marked as "yes" or "no"); (6-7) the starting and ending position of the gene's genomic position; (8) number of SNPs within a gene that were included in analysis; (9) the most significant SNP according to GWA summary statistics; (10) the $P$-value of the most significant SNP; and, on the first page, (11) the corresponding gene-level posterior enrichment probability as found by RSS for comparison. Note that an "NA" in column (11) occurs wherever the MCMC for RSS failed to converge. Highlighted rows represent enriched genes whose top SNP is not marginally significant according to a genome-wide Bonferroni-corrected threshold ($P = 4.67\times10^{-8}$ correcting for 1,070,306 SNPs analyzed). (XLSX)

**Table S20. Significant genes for body mass index (BMI) in the UK Biobank analysis using gene-$\varepsilon$-EN.** Here, we analyze 17,680 genes from $N$ = 349,468 individuals of European-ancestry. This file gives the gene-$\varepsilon$ gene-level association $P$-values using Elastic Net regularized effect sizes when gene boundaries are defined by (page 1) using UCSC annotations directly, and (page 2) augmenting the gene boundaries by adding SNPs within a ±50kb buffer. Significance was determined by using a Bonferroni-corrected $P$-value threshold (in our analyses, $P = 0.05/14322$ autosomal genes $= 3.49\times10^{-6}$ and $P = 0.05/17680$ autosomal genes $= 2.83\times10^{-6}$, respectively). The columns of tables on both pages provide: (1) chromosome position; (2) gene name; (3) gene-$\varepsilon$-EN gene $P$-value; (4) gene-specific heritability estimates; (5) whether or not an association between gene and trait is listed in the GWAS catalog (marked as "yes" or "no"); (6-7) the starting and ending position of the gene's genomic position; (8) number of SNPs within a gene that were included in analysis; (9) the most significant SNP according to GWA summary statistics; (10) the $P$-value of the most significant SNP; and, on the first page, (11) the corresponding gene-level posterior enrichment probability as found by RSS for comparison. Note that an "NA" in column (11) occurs wherever the MCMC for RSS failed to converge. Highlighted rows represent enriched genes whose top SNP is not marginally significant according to a genome-wide Bonferroni-corrected threshold ($P = 4.67\times10^{-8}$ correcting for 1,070,306 SNPs analyzed). (XLSX)

**Table S21. Significant genes for mean corpuscular volume (MCV) in the UK Biobank analysis using gene-$\varepsilon$-EN.** Here, we analyze 17,680 genes from $N = 349{,}468$ individuals of European-ancestry. This file gives the gene-$\varepsilon$ gene-level association $P$-values using Elastic Net regularized effect sizes when gene boundaries are defined by (page 1) using UCSC annotations directly, and (page 2) augmenting the gene boundaries by adding SNPs within a $\pm$50kb buffer. Significance was determined by using a Bonferroni-corrected $P$-value threshold (in our analyses, $P = 0.05/14322$ autosomal genes $= 3.49 \times 10^{-6}$ and $P = 0.05/17680$ autosomal genes $= 2.83 \times 10^{-6}$, respectively). The columns of tables on both pages provide: (1) chromosome position; (2) gene name; (3) gene-$\varepsilon$-EN gene $P$-value; (4) gene-specific heritability estimates; (5) whether or not an association between gene and trait is listed in the GWAS catalog (marked as "yes" or "no"); (6-7) the starting and ending position of the gene's genomic position; (8) number of SNPs within a gene that were included in analysis; (9) the most significant SNP according to GWA summary statistics; (10) the $P$-value of the most significant SNP; and, on the first page, (11) the corresponding gene-level posterior enrichment probability as found by RSS for comparison. Note that an "NA" in column (11) occurs wherever the MCMC for RSS failed to converge. Highlighted rows represent enriched genes whose top SNP is not marginally significant according to a genome-wide Bonferroni-corrected threshold ($P = 4.67 \times 10^{-8}$ correcting for 1,070,306 SNPs analyzed). (XLSX)

**Table S22. Significant genes for mean platelet volume (MPV) in the UK Biobank analysis using gene-$\varepsilon$-EN.** Here, we analyze 17,680 genes from $N = 349{,}468$ individuals of European-ancestry. This file gives the gene-$\varepsilon$ gene-level association $P$-values using Elastic Net regularized effect sizes when gene boundaries are defined by (page 1) using UCSC annotations directly, and (page 2) augmenting the gene boundaries by adding SNPs within a $\pm$50kb buffer. Significance was determined by using a Bonferroni-corrected $P$-value threshold (in our analyses, $P = 0.05/14322$ autosomal genes $= 3.49 \times 10^{-6}$ and $P = 0.05/17680$ autosomal genes $= 2.83 \times 10^{-6}$, respectively). The columns of tables on both pages provide: (1) chromosome position; (2) gene name; (3) gene-$\varepsilon$-EN gene $P$-value; (4) gene-specific heritability estimates; (5) whether or not an association between gene and trait is listed in the GWAS catalog (marked as "yes" or "no"); (6-7) the starting and ending position of the gene's genomic position; (8) number of SNPs within a gene that were included in analysis; (9) the most significant SNP according to GWA summary statistics; (10) the $P$-value of the most significant SNP; and, on the first page, (11) the corresponding gene-level posterior enrichment probability as found by RSS for comparison. Note that an "NA" in column (11) occurs wherever the MCMC for RSS failed to converge. Highlighted rows represent enriched genes whose top SNP is not marginally significant according to a genome-wide Bonferroni-corrected threshold ($P = 4.67 \times 10^{-8}$ correcting for 1,070,306 SNPs analyzed). (XLSX)

**Table S23. Significant genes for platelet count (PLC) in the UK Biobank analysis using gene-$\varepsilon$-EN.** Here, we analyze 17,680 genes from $N$ = 349,468 individuals of European-ancestry. This file gives the gene-$\varepsilon$ gene-level association $P$-values using Elastic Net regularized effect sizes when gene boundaries are defined by (page 1) using UCSC annotations directly, and (page 2) augmenting the gene boundaries by adding SNPs within a ±50kb buffer. Significance was determined by using a Bonferroni-corrected $P$-value threshold (in our analyses, $P = 0.05/14322$ autosomal genes $= 3.49\times10^{-6}$ and $P = 0.05/17680$ autosomal genes $= 2.83\times10^{-6}$, respectively). The columns of tables on both pages provide: (1) chromosome position; (2) gene name; (3) gene-$\varepsilon$-EN gene $P$-value; (4) gene-specific heritability estimates; (5) whether or not an association between gene and trait is listed in the GWAS catalog (marked as "yes" or "no"); (6-7) the starting and ending position of the gene's genomic position; (8) number of SNPs within a gene that were included in analysis; (9) the most significant SNP according to GWA summary statistics; (10) the $P$-value of the most significant SNP; and, on the first page, (11) the corresponding gene-level posterior enrichment probability as found by RSS for comparison. Note that an "NA" in column (11) occurs wherever the MCMC for RSS failed to converge. Highlighted rows represent enriched genes whose top SNP is not marginally significant according to a genome-wide Bonferroni-corrected threshold ($P = 4.67 \times 10^{-8}$ correcting for 1,070,306 SNPs analyzed). (XLSX)

**Table S24. Significant genes for waist-hip ratio (WHR) in the UK Biobank analysis using gene-$\varepsilon$-EN.** Here, we analyze 17,680 genes from $N$ = 349,468 individuals of European-ancestry. This file gives the gene-$\varepsilon$ gene-level association $P$-values using Elastic Net regularized effect sizes when gene boundaries are defined by (page 1) using UCSC annotations directly, and (page 2) augmenting the gene boundaries by adding SNPs within a ±50kb buffer. Significance was determined by using a Bonferroni-corrected $P$-value threshold (in our analyses, $P = 0.05/14322$ autosomal genes $= 3.49 \times 10^{-6}$ and $P = 0.05/17680$ autosomal genes $= 2.83\times10^{-6}$, respectively). The columns of tables on both pages provide: (1) chromosome position; (2) gene name; (3) gene-$\varepsilon$-EN gene $P$-value; (4) gene-specific heritability estimates; (5) whether or not an association between gene and trait is listed in the GWAS catalog (marked as "yes" or "no"); (6-7) the starting and ending position of the gene's genomic position; (8) number of SNPs within a gene that were included in analysis; (9) the most significant SNP according to GWA summary statistics; (10) the $P$-value of the most significant SNP; and, on the first page, (11) the corresponding gene-level posterior enrichment probability as found by RSS for comparison. Note that an "NA" in column (11) occurs wherever the MCMC for RSS failed to converge. Highlighted rows represent enriched genes whose top SNP is not marginally significant according to a genome-wide Bonferroni-corrected threshold ($P = 4.67 \times 10^{-8}$ correcting for 1,070,306 SNPs analyzed). (XLSX)

### S4 Additional Detailed Results for Traits in the UK Biobank

In this section, we present additional detailed findings and results from applying gene-$\varepsilon$ to the six quantitative traits — height, body mass index (BMI), mean red blood cell volume (MCV), mean platelet volume (MPV), platelet count (PLC), waist-hip ratio (WHR) — assayed in self-identified European-ancestry individuals in the UK Biobank [1]. For these extra set of analyses, we obtained the genotype data release (without imputed genotypes) and implemented the same quality control procedure that was used in the main text (Section S1). This resulted in a final dataset of $N$ = 349,468 individuals and $J$ = 410,172 genome-wide SNPs. Once again, we used the NCBI's Reference Sequence (RefSeq) database in the UCSC Genome Browser [3] to annotate SNPs with the appropriate genes in one of two ways. In the first setting, we use the UCSC gene boundary definitions directly; while in the second setting, we augment the gene boundaries by adding SNPs within a ±50 kilobase (kb) buffer to account for possible regulatory elements. Genes with only 1 SNP in their boundary were excluded from the respective analysis. For these data, a total of 13,029 autosomal genes were analyzed when using the UCSC boundaries as defined; while, a total of 17,680 autosomal genes were analyzed when including the 50kb buffer. Lastly, we regressed the top ten principal components of the genotype data onto each trait to control for population structure, and then we derived OLS SNP-level effect sizes using the traditional GWA framework. Here, our goal is to compare how the four different implementations of gene-$\varepsilon$ (i.e., OLS with no regularization, Ridge Regression, Elastic Net, and LASSO) analyze these summary statistics.

As shown in the main text, we begin with assessing how the various regularization solutions result in different characterizations of genetic architectures (Table S25). In general, we find the same general themes we saw in our simulation study. Less aggressive shrinkage approaches (e.g., OLS and Ridge) are subject to misclassifications of associated, spurious, and non-associated SNPs. As result, these methods struggle to avoid identifying false positive SNP-level associations, across all six traits. For example, gene-$\varepsilon$-OLS assumes that approximately 54% and 50% of the SNPs analyzed are associated with BMI and WHR, respectively. This once again highlights the need for computational frameworks that are able to appropriately correct for inflation in summary statistics.

Lastly, we applied each version of gene-$\varepsilon$ to the (regularized) GWA summary statistics and generated genome-wide gene-level association $P$-values. Recall that we are motivated to identify enriched genes, which we define as a gene containing at least one associated SNP and achieving a gene-level association $P$-value below a Bonferroni-corrected significance threshold. In our analyses, this significance threshold is $P = 0.05/13029$ autosomal genes $= 3.84 \times 10^{-6}$ when the UCSC gene boundaries are used directly, and $P = 0.05/17680$ autosomal genes $= 2.83 \times 10^{-6}$ when the ±50kb buffer is applied, respectively. As a validation step, we used the gene set enrichment analysis tool Enrichr [30] to identify dbGaP categories with an overrepresentation of significant genes reported by the four different implementations of gene-$\varepsilon$. A comparison of gene-level associations and gene set enrichments between the each gene-$\varepsilon$ approaches are also listed (Tables S26 and S27). Note that, similar to the main text, we use the findings of gene-$\varepsilon$-EN as the reference.

| gene-$\varepsilon$ Approach | Trait | # Mix. Comp. | % Associated SNPs | % Causal SNPs | $\varepsilon$-genic Threshold ($\sigma_2^2$) |
|---|---|---|---|---|---|
| Elastic Net | Height | 7 | 11.65% | 1.27% | $1.90 \times 10^{-5}$ |
| | BMI | 6 | 16.61% | 7.60% | $2.73 \times 10^{-5}$ |
| | MCV | 8 | 11.60% | 0.21% | $3.48 \times 10^{-5}$ |
| | MPV | 8 | 5.47% | 0.20% | $3.37 \times 10^{-5}$ |
| | PLC | 7 | 10.33% | 0.39% | $3.68 \times 10^{-5}$ |
| | WHR | 8 | 14.38% | 7.20% | $3.67 \times 10^{-5}$ |
| LASSO | Height | 8 | 10.66% | 0.67% | $2.29 \times 10^{-5}$ |
| | BMI | 6 | 16.57% | 7.35% | $2.71 \times 10^{-5}$ |
| | MCV | 7 | 9.94% | 0.23% | $3.56 \times 10^{-5}$ |
| | MPV | 9 | 4.51% | 0.20% | $3.32 \times 10^{-5}$ |
| | PLC | 8 | 8.57% | 0.26% | $3.80 \times 10^{-4}$ |
| | WHR | 6 | 15.27% | 7.05% | $3.29 \times 10^{-4}$ |
| Ridge Regression | Height | 5 | 47.73% | 13.93% | $5.67 \times 10^{-7}$ |
| | BMI | 5 | 38.53% | 17.98% | $1.49 \times 10^{-7}$ |
| | MCV | 5 | 45.28% | 9.79% | $8.76 \times 10^{-6}$ |
| | MPV | 9 | 34.42% | 2.69% | $1.13 \times 10^{-7}$ |
| | PLC | 5 | 46.14% | 8.59% | $8.71 \times 10^{-6}$ |
| | WHR | 5 | 42.09% | 19.12% | $1.31 \times 10^{-7}$ |
| OLS | Height | 5 | 49.08% | 6.36% | $3.08 \times 10^{-5}$ |
| | BMI | 3 | 54.54% | 25.25% | $3.31 \times 10^{-5}$ |
| | MCV | 9 | 34.39% | 1.96% | $5.55 \times 10^{-5}$ |
| | MPV | 9 | 36.07% | 2.16% | $6.74 \times 10^{-5}$ |
| | PLC | 9 | 36.95% | 1.84% | $6.28 \times 10^{-5}$ |
| | WHR | 3 | 50.03% | 23.81% | $3.49 \times 10^{-5}$ |

**Table S25. Characterization of the genetic architectures of six traits assayed in European-ancestry individuals in the UK Biobank (using un-imputed genotypes).** Here, we report the way different regularizations in gene-$\varepsilon$ characterize $\varepsilon$-genic effects in complex traits. Results are shown for Elastic Net (which is highlighted in the main text), as well as for LASSO and Ridge Regression. We also show results when no shrinkage is applied to illustrate the importance of this step (denoted by OLS). In the three former cases, we regress the GWA SNP-level effect size estimates onto chromosome-specific LD matrices to derive a regularized set of summary statistics $\widetilde{\boldsymbol{\beta}}$. gene-$\varepsilon$ assumes a reformulated null distribution of SNP-level effects $\widetilde{\beta}_j \sim \mathcal{N}(0, \sigma_\varepsilon^2)$, where $\sigma_\varepsilon^2$ is the SNP-level null threshold and represents the maximum proportion of phenotypic variance explained (PVE) by a spurious or non-associated SNP. We used an EM-algorithm with 100 iterations to fit $K$-mixture Gaussian models over the regularized effect sizes to estimate $\sigma_\varepsilon^2$. Here, each mixture component had distinctively smaller variances ($\sigma_1^2 > \cdots > \sigma_K^2$; with the $K$-th component fixed at $\sigma_K^2 = 0$), and the number of total mixture components $K$ was chosen based on a grid of values where the best model yielded the highest Bayesian Information Criterion (BIC). We assume associated SNPs appear in the first component, non-associated SNPs appear in the last component, and null SNPs with spurious effects fell in between (i.e., $\sigma_\varepsilon^2 = \sigma_2^2$). Thus, a SNP is considered to have some level of association with a trait if $\mathbb{E}[\beta_j^2] > \sigma_K^2 = 0$; while a SNP is considered "causal" if $\mathbb{E}[\beta_j^2] > \sigma_2^2$. Column 3 gives the $K$ used for each trait. Column 4 and 5 detail the percentage of associated and causal SNPs, respectively. The last column gives the mean threshold for $\varepsilon$-genic effects across the chromosomes.

| | Trait | OLS | Ridge Regression | LASSO | Elastic Net |
|---|---|---|---|---|---|
| # Sig. Genes | Height | 501 | 8 | 65 | 67 |
| | BMI | 640 | 8 | 42 | 40 |
| | MCV | 318 | 10 | 62 | 78 |
| | MPV | 326 | 29 | 62 | 66 |
| | PLC | 289 | 15 | 54 | 52 |
| | WHR | 677 | 6 | 49 | 22 |
| % Sig. Gene Overlap w/ Elastic Net | Height | 7.78% | 37.50% | 69.23% | — |
| | BMI | 2.19% | 12.50% | 64.29% | — |
| | MCV | 12.89% | 60.00% | 70.97% | — |
| | MPV | 13.80% | 51.70% | 83.87% | — |
| | PLC | 11.42% | 46.67% | 75.93% | — |
| | WHR | 1.18% | 0.00% | 20.41% | — |
| # Enriched dbGaP Categories | Height | 1 | 1 | 0 | 1 |
| | BMI | 33 | 16 | 0 | 0 |
| | MCV | 1 | 3 | 1 | 1 |
| | MPV | 6 | 1 | 2 | 2 |
| | PLC | 2 | 3 | 1 | 2 |
| | WHR | 23 | 3 | 0 | 0 |
| % Enriched dbGaP Overlap w/ Elastic Net | Height | 100.00% (Body Height) | 100.00% (Body Height) | 0.00% | — |
| | BMI | 0.00% | 0.00% | 0.00% | — |
| | MCV | 100.00% (Erythrocyte Indices) | 33.33% (Erythrocyte Indices) | 100.00% (Erythrocyte Indices) | — |
| | MPV | 16.67% (Platelet Count) | 100.00% (Platelet Count) | 100.00% (Platelet Count; Face) | — |
| | PLC | 50.00% (Platelet Count) | 33.33% (Platelet Count) | 100.00% (Platelet Count) | — |
| | WHR | 0.00% | 0.00% | 0.00% | — |

**Table S26. Comparison of the different gene-$\varepsilon$ approaches on the six quantitative traits assayed in European-ancestry individuals from the UK Biobank un-imputed genotyped data.** Traits include: height; body mass index (BMI); mean corpuscular volume (MCV); mean platelet volume (MPV); platelet count (PLC); and waist-hip ratio (WHR). Here, we list the number of significant genes found when using gene-$\varepsilon$ with various regularization strategies, as well as the number of dbGAP categories enriched for significant genes identified by gene-$\varepsilon$. We also assess how well these results overlap with the gene-$\varepsilon$ -EN findings that were reported in the main text. Significant genes were determined by using a Bonferroni-corrected $P$-value threshold (in our analyses, $P = 0.05/13029$ autosomal genes $= 3.84 \times 10^{-6}$). Enriched dbGAP categories were those with Enrichr $Q$-values (i.e., false discovery rates) less than 0.05.

| | Trait | OLS | Ridge Regression | LASSO | Elastic Net |
|---|---|---|---|---|---|
| # Sig. Genes | Height | 859 | 21 | 90 | 71 |
| | BMI | 770 | 9 | 14 | 73 |
| | MCV | 564 | 90 | 86 | 104 |
| | MPV | 595 | 119 | 83 | 80 |
| | PLC | 517 | 73 | 75 | 69 |
| | WHR | 721 | 4 | 25 | 4 |
| % Sig. Gene Overlap w/ Elastic Net | Height | 6.82% | 42.86% | 70.00% | — |
| | BMI | 1.69% | 11.11% | 64.29% | — |
| | MCV | 13.83% | 46.67% | 83.72% | — |
| | MPV | 12.77% | 50.42% | 84.34% | — |
| | PLC | 11.99% | 42.47% | 85.33% | — |
| | WHR | 0.28% | 0.00% | 12.00% | — |
| # Enriched dbGaP Categories | Height | 3 | 1 | 1 | 1 |
| | BMI | 30 | 7 | 0 | 0 |
| | MCV | 2 | 2 | 4 | 1 |
| | MPV | 9 | 4 | 2 | 3 |
| | PLC | 5 | 1 | 1 | 1 |
| | WHR | 10 | 0 | 0 | 0 |
| % Enriched dbGaP Overlap w/ Elastic Net | Height | 33.33% (Body Height) | 100.00% (Body Height) | 100.00% (Body Height) | — |
| | BMI | 0.00% | 0.00% | 0.00% | — |
| | MCV | 50.00% (Erythrocyte Indices) | 50.00% (Erythrocyte Indices) | 25.00% (Erythrocyte Indices) | — |
| | MPV | 11.11% (Platelet Count) | 75.00% (Platelet Count; Hearing Loss; Face) | 100.00% (Platelet Count; Face) | — |
| | PLC | 20.00% (Platelet Count) | 100.00% (Platelet Count) | 100.00% (Platelet Count) | — |
| | WHR | 0.00% | 0.00% | 0.00% | — |

**Table S27. Comparison of the different gene-$\varepsilon$ approaches on the six quantitative traits assayed in European-ancestry individuals from the UK Biobank un-imputed genotyped data with gene boundaries augmented by a 50 kilobase (kb) buffer.** Traits include: height; body mass index (BMI); mean corpuscular volume (MCV); mean platelet volume (MPV); platelet count (PLC); and waist-hip ratio (WHR). Here, we list the number of significant genes found when using gene-$\varepsilon$ with various regularization strategies, as well as the number of dbGAP categories enriched for significant genes identified by gene-$\varepsilon$. We also assess how well these results overlap with the gene-$\varepsilon$ -EN findings that were reported in the main text. Significant genes were determined by using a Bonferroni-corrected $P$-value threshold (in our analyses, $P = 0.05/17680$ autosomal genes $= 2.83 \times 10^{-6}$). Enriched dbGAP categories were those with Enrichr $Q$-values (i.e., false discovery rates) less than 0.05.
